# Developmental variation in pterygoid segmentation clarifies patterns of avian bony palate evolution

**DOI:** 10.64898/2026.03.24.713852

**Authors:** Annabel K. Hunt, Juan Benito, Olivia Plateau, Adam Urantówka, Daniel J. Field

## Abstract

The morphology of the palate has long constituted the primary basis for differentiating between the two deepest clades of crown group birds, Neognathae and Palaeognathae. However, published literature on the bird palate is dominated by classical anatomical descriptions pre-dating the advent of contemporary three-dimensional imaging techniques, hindering our understanding of bird palate disparity and development. Pterygoid segmentation, the process by which the rostral portion of the pterygoid separates and fuses with the palatine during ontogeny in neognathous birds, remains a poorly understood aspect of avian cranial development despite giving rise to an important component of the cranial kinetic system. Here, we use micro-computed tomography to explore ontogenetic change of the palate during the process of pterygoid segmentation across an unprecedentedly broad taxonomic sample of immature and mature birds. With this large dataset, we present direct evidence for the full process of post-hatching pterygoid segmentation being exclusively restricted to the major avian subclade Neoaves. Further, our investigation helps refine several hypotheses of homology related to the pterygoid-palatine contact in crown birds and their close stem-group relatives: we hypothesise that the rostral projection of the pterygoid observed in Anseres (and potentially Anhimidae and Megapodiidae) is homologous with the hemipterygoid of neoavians, and that the discrete and unfused hemipterygoids of some crownward stem-birds suggest that a discrete hemipterygoid originated prior to the origin of the process of pterygoid segmentation. However, whether pterygoid segmentation represents a synapomorphy of Neornithes, Neognathae, or Neoaves remains ambiguous in light of elusive data from the fossil record and early stages of cranial development. Overall, our study clarifies the process of pterygoid segmentation in crown birds and raises new questions regarding the major morphological modifications that have characterized the evolutionary history of the avian bony palate.

## 1. INTRODUCTION

Neornithes (crown group birds) are classified into two major clades: Palaeognathae and Neognathae, with neognaths comprising the bulk of extant species diversity with over 11,000 species (Billerman et al., 2025). By contrast, extant Palaeognathae comprise only 60 species of volant tinamous and flightless ratites (Widrig & Field, 2022). The organization and morphology of the palate constitutes a key feature distinguishing both clades, as first documented over 150 years ago by Huxley (1867) (fig. 1), but it was only in 1900 that Pycraft introduced the terms ‘Palaeognathae’ (“ancient jaws”) and ‘Neognathae’ (“new jaws”), explicitly tying assumptions about the relative antiquity of each clade’s palatal arrangement to their names (Pycraft, 1900).

**Figure 1:**
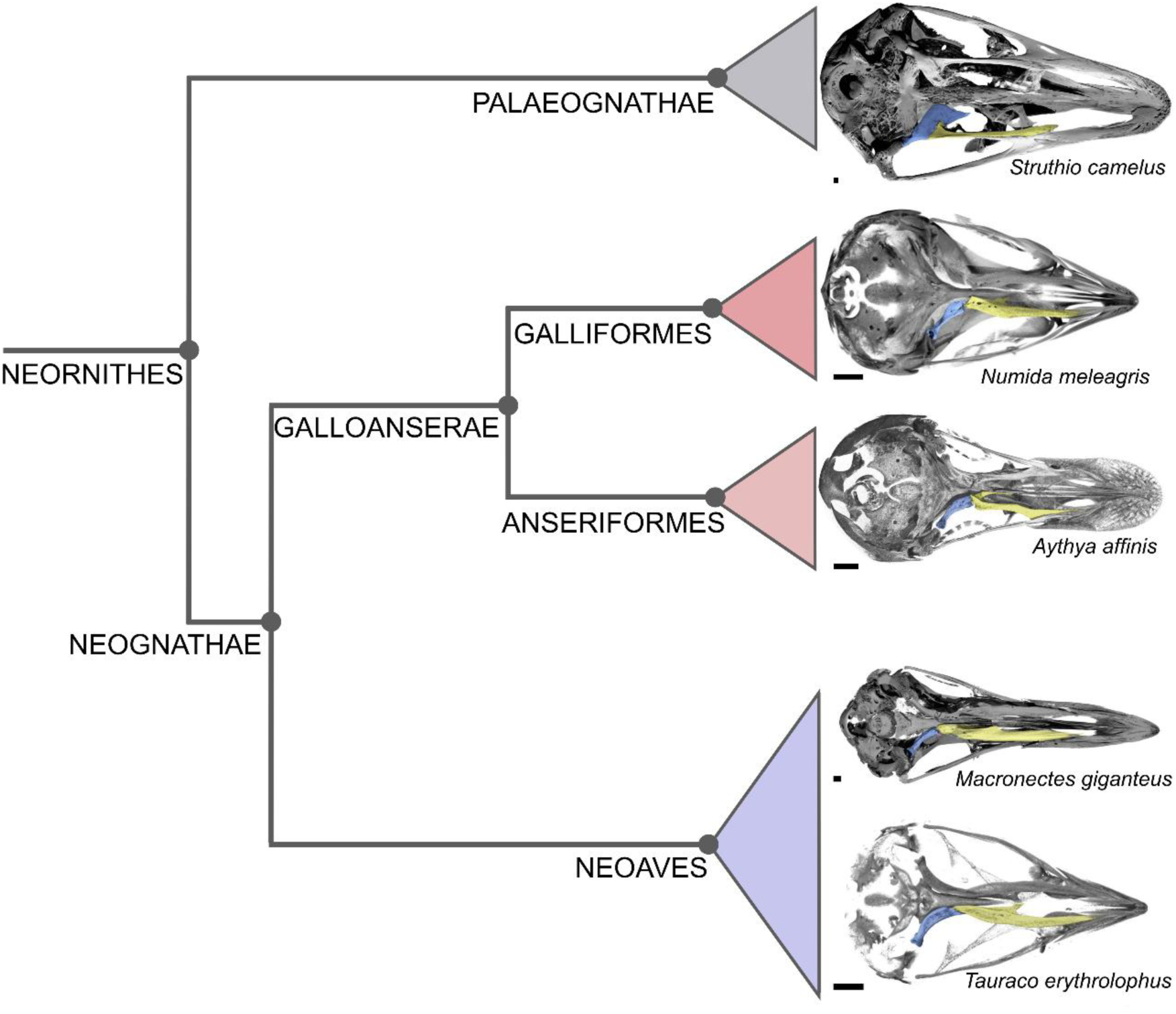
Simplified phylogeny of crown birds (adapted from Prum et al. (2015)) with crania illustrated in ventral view for selected representatives of major clades, from top to bottom: *Struthio camelus* (Common Ostrich); *Numida meleagris* (Helmeted Guineafowl); *Aythya affinis* (Lesser Scaup); *Macronectes giganteus* (Southern Giant Petrel) and *Tauraco erythrolophus* (Red-Crested Turaco). All individuals illustrated are immatures, apart from *Struthio camelus* (subadult). Pterygoid illustrated in blue and palatine illustrated in yellow. All scale bars represent 2.5 mm.

The avian bony palate is a complex structure consisting of paired pterygoids and palatines, and an unpaired vomer (which itself is formed through the fusion of independent precursors during development; e.g., Zusi (1993)), with all of these elements exhibiting variation in morphology, size and relative position among major neornithine subclades. In addition to differences related to the proportions of the vomer and its relationships with other components of the palate (e.g., Pycraft (1900); Pycraft (1901); Hu et al. (2019)) an important feature distinguishing the palates of palaeognaths and neognaths is the relationship between the pterygoid and palatine (figs. 1, 2). The pterygoid and palatine fuse into an immobile complex in palaeognaths, while in neognaths these elements articulate at a mobile joint (e.g., Pycraft (1901); Bock (1963); Widrig and Field (2022)). The mobile articulation between the pterygoid and palatine contributes to the enhanced kinetic capacity of the neognath skull relative to palaeognaths (Simonetta, 1960), which has been suggested to represent a factor underpinning the astonishing taxonomic disparity between these clades (Hu et al., 2019).

**Figure 2:**
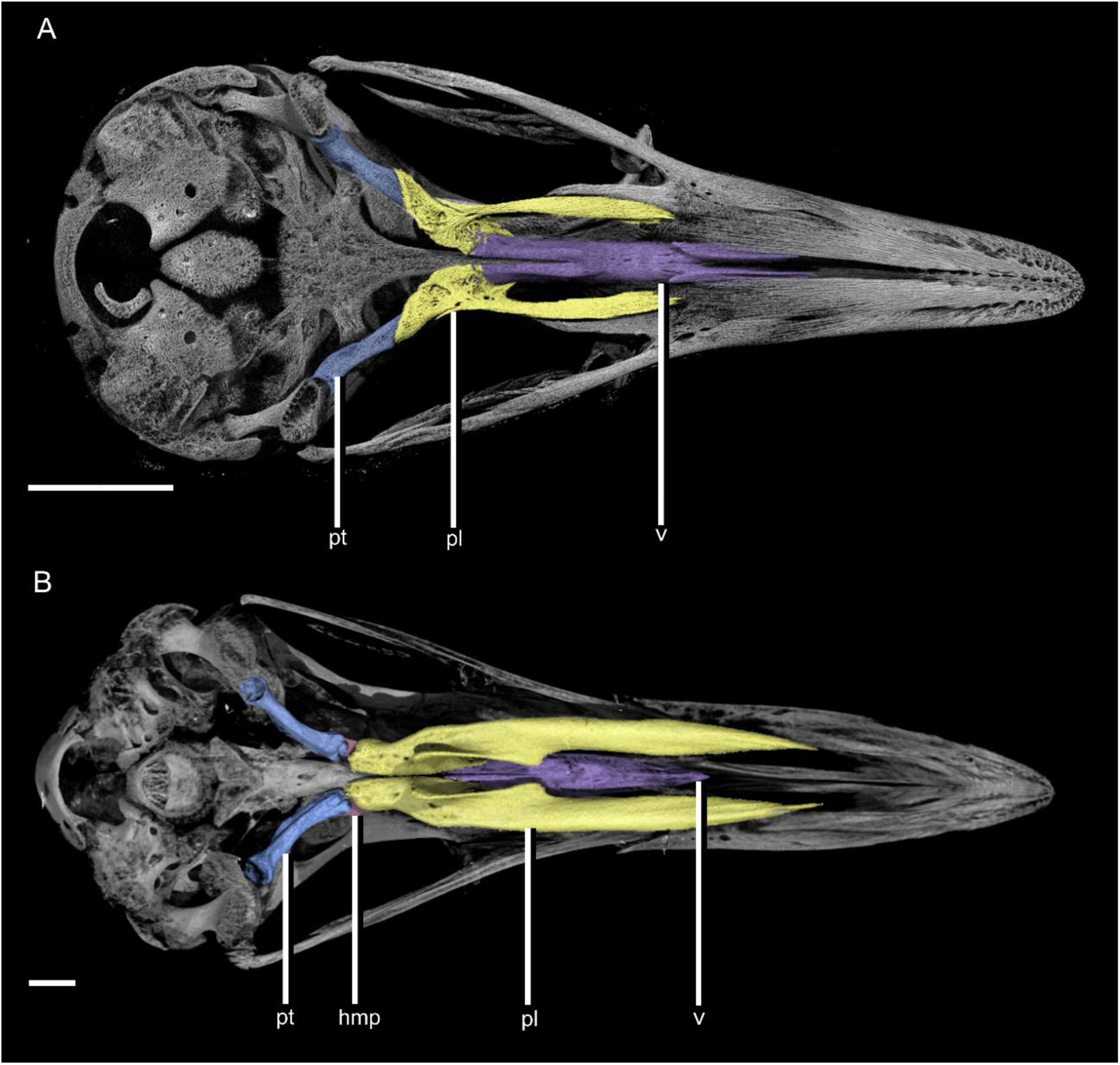
Ventral view of the cranium of an immature palaeognath and an immature neognath with palatal elements digitally segmented and coloured. **A**, palaeognath (*Tinamus solitarius*); **B**, neognath (*Macronectes giganteus*). Abbreviations: hmp, hemipterygoid (pink – largely obscured); pl, palatine (yellow); pt, pterygoid (blue); v, vomer (purple). Both scale bars represent 5 mm.

Here, we focus on the remarkable developmental origin of the mobile pterygoid-palatine joint characterising Neognathae. In most neognaths, this mobile articulation arises in part through a process termed ‘pterygoid segmentation’ ((Pycraft, 1898); (Pycraft, 1901)) - in which the rostral portion of the pterygoid detaches from the main pterygoid body (the ‘hemipterygoid’) and fuses with the caudal portion of the palatine, such that in mature birds the pterygoid represents only the posterior component of the original element. During this process, which has been considered a neognath synapomorphy (e.g., Balouet (1983); Witmer and Martin (1987)), an ‘intrapterygoid’ articulation forms between the rostral end of the remnant pterygoid body (now reduced in size as a result of the loss of the hemipterygoid), and the caudal end of the palatine, which is augmented by fusion with the hemipterygoid (fig. 3) ((Pycraft, 1898); (Pycraft, 1901); (Zusi & Livezey, 2006)).

**Figure 3:**
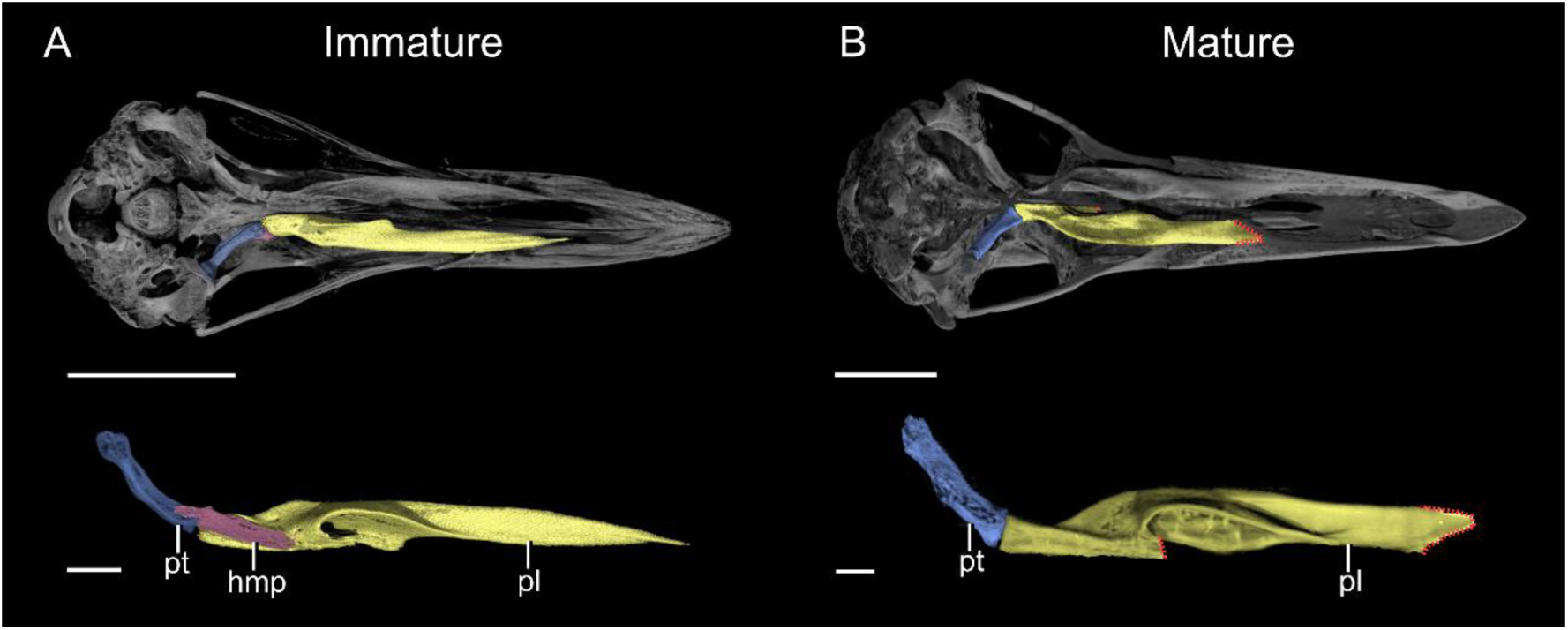
Ontogenetic change in palate morphology of *Macronectes* (Neoaves: Procellariidae). **A**, immature *Macronectes giganteus.* Top: ventral view of the cranium with palatal elements digitally segmented and coloured. Bottom: Dorsal view of these digitally segmented palatal elements; **B**, mature *Macronectes halli.* Top: ventral view of the cranium with palatal elements digitally segmented and coloured. Bottom: Dorsal view of these digitally segmented palatal elements. Red dotted lines represent areas where bone margins cannot be precisely delineated due to fusion. Abbreviations: hmp, hemipterygoid (pink); pl, palatine (yellow); pt, pterygoid (blue). Scale bars in A: top scale bar represents 25 mm; bottom scale bar represents 5 mm; B: top scale bar represents 25 mm; bottom scale bar represents 5 mm.

Despite the important role played by pterygoid segmentation in the formation of the mobile pterygoid-palatine joint and in facilitating powered palatal kinesis, the developmental process of segmentation remains poorly understood. Variable anatomical terminology in previous studies has contributed to our incomplete understanding of pterygoid segmentation (and the associated hemipterygoid) among neognath subclades (e.g., Parker (1872); Parker (1875); Parker (1876); Parker (1878); Pycraft (1898); Balouet (1983); Bühler et al. (1988); Pycraft (1901); Pycraft (1903); Pycraft (1907a); de Beer (1937); Jollie (1957); Jollie (1958); Zusi and Livezey (2006); Livezey and Zusi (2006)). Moreover, literature on the process of pterygoid segmentation is dominated by classical comparative studies in which the small size of the elements involved, their ontogenetically variable nature, and their topological positions have, at times, impeded unambiguous anatomical interpretation. For instance, classical investigations of avian palate morphology typically illustrate the palate in ventral or lateral view, obstructing interpretation of the dorsally positioned hemipterygoid. Three-dimensional digital reconstructions of palate morphology therefore have potential to allow more detailed insights into the topological relationships of palatal elements throughout the process of pterygoid segmentation.

To clarify the process of avian pterygoid segmentation, we use micro-computed tomography (μCT) to illustrate ontogenetic change of the pterygoid-palatine contact across a broad sample of extant avian diversity, and, based on these observations, propose more precise anatomical terminology to describe the process. Our results illustrate the post-hatching ontogenetic origin of the mobile pterygoid-palatine joint in neognathous birds, with implications for informing avian palate evolution in crown birds and their closest stem-group relatives.

## 2. MATERIALS AND METHODS

We used micro-computed tomography (µCT) to examine the palate morphology of 69 extant bird specimens representing 50 unique species-level taxa (43 immature and 26 comparatively mature individuals), across the three major extant clades of Neornithes (Palaeognathae, Galloanserae and Neoaves) and 21 ordinal-level subclades (suppl. table 1). We also present data from the subadult subfossil holotype specimen of the extinct moa *Megalapteryx didinus* ((Owen, 1883); (Rawlence et al., 2013)) to allow us to more fully characterise palaeognath palatal disparity, though we were unable to incorporate data from additional ontogenetic stages for this taxon. Further, for the purposes of our discussion, we also examined the pterygoids of two total-group anseriform fossils: *Nettapterornis* and *Presbyornis* (suppl. table 1).

We focus primarily on immature specimens (late-stage embryos, hatchlings, or juveniles; suppl. table 1) to enable identification of the hemipterygoid as a distinct element prior to its eventual fusion with the palatine, and to observe other individual palate bones prior to bone fusion. For extant taxa, we additionally examined at least one mature representative within each sampled taxonomic order to characterise the process of pterygoid segmentation and illustrate changes in palatal configuration during ontogeny. We also visually examined additional immature and mature specimens for comparative purposes (suppl. table 2).

We used VGSTUDIOMAX software version 3.4.5 and version 2023.2 (VolumeGraphics) to digitally segment the palatines, pterygoids and, when present, hemipterygoids manually across our sample. To describe the process of pterygoid segmentation we focused on the points of articulation among the palatine, pterygoid and hemipterygoid. We therefore describe the caudalmost portion of the palatine (particularly the pterygoid process (*processus pterygoideus,* (Baumel & Witmer, 1993))), the rostral portion of the pterygoid, and the hemipterygoid itself. For other relevant regions of the palatine that we describe, we use the term rostral process (*processus rostralis palatini* ((Zusi & Livezey, 2006); (Livezey & Zusi, 2006))) to refer to a distinct laterally positioned rostral projection of the palatine, while we use the term rostromedial process (*processus rostromedialis palatini* ((Zusi & Livezey, 2006); (Livezey & Zusi, 2006))) to refer to a distinct medially positioned rostral projection of the palatine. We also use the term ‘caudolateral corner’ as a simplification of “*angulus caudolateralis palatini*” of Zusi and Livezey (2006) (see their fig. 7).

We apply the term ‘hemipterygoid’ to the distinct, identifiable element that forms during segmentation of the pterygoid ((Pycraft, 1898); (Pycraft, 1901)). Further, throughout this study we discuss the potential homologies of the rostral projection of the pterygoid (present across much of extant neornithine diversity) with this distinct hemipterygoid element.

Here, the distinct hemipterygoid may be identifiable as a discernible element directly attached to the pterygoid body, present as an entirely separate (discrete) element unfused to any other, or as a separate element in the process of fusion to the palatine. We therefore use the terms ‘partially segmented’, and ‘fully segmented’ to describe the relationship between the distinct hemipterygoid and the pterygoid body. We use the term ‘partially segmented’ to refer to the situation where the hemipterygoid remains attached to the pterygoid, but is readily distinguishable from it in µCT scans due to the presence of a suture separating it from the remainder of the pterygoid, as well by differences in bone texture and density (fig. 4). The suture between the rostral pterygoid and the caudal hemipterygoid was particularly challenging to distinguish in scans that were of lower resolution or of relatively weakly ossified specimens (fig. 4). ‘Fully segmented’ hemipterygoids, by contrast, represent elements entirely separate from the pterygoid body, which were therefore straightforward to identify. ‘Fully segmented’ hemipterygoids that had begun fusing with the caudal palatine (and thus lack a clearly identifiable rostral margin) (fig. 4) are specifically noted in the relevant figure captions. Further, where bone margins of other palatal regions were not readily distinguishable (either due to scan resolution or bone fusion), we use red dotted lines to mark these regions in our figures.

**Figure 4:**
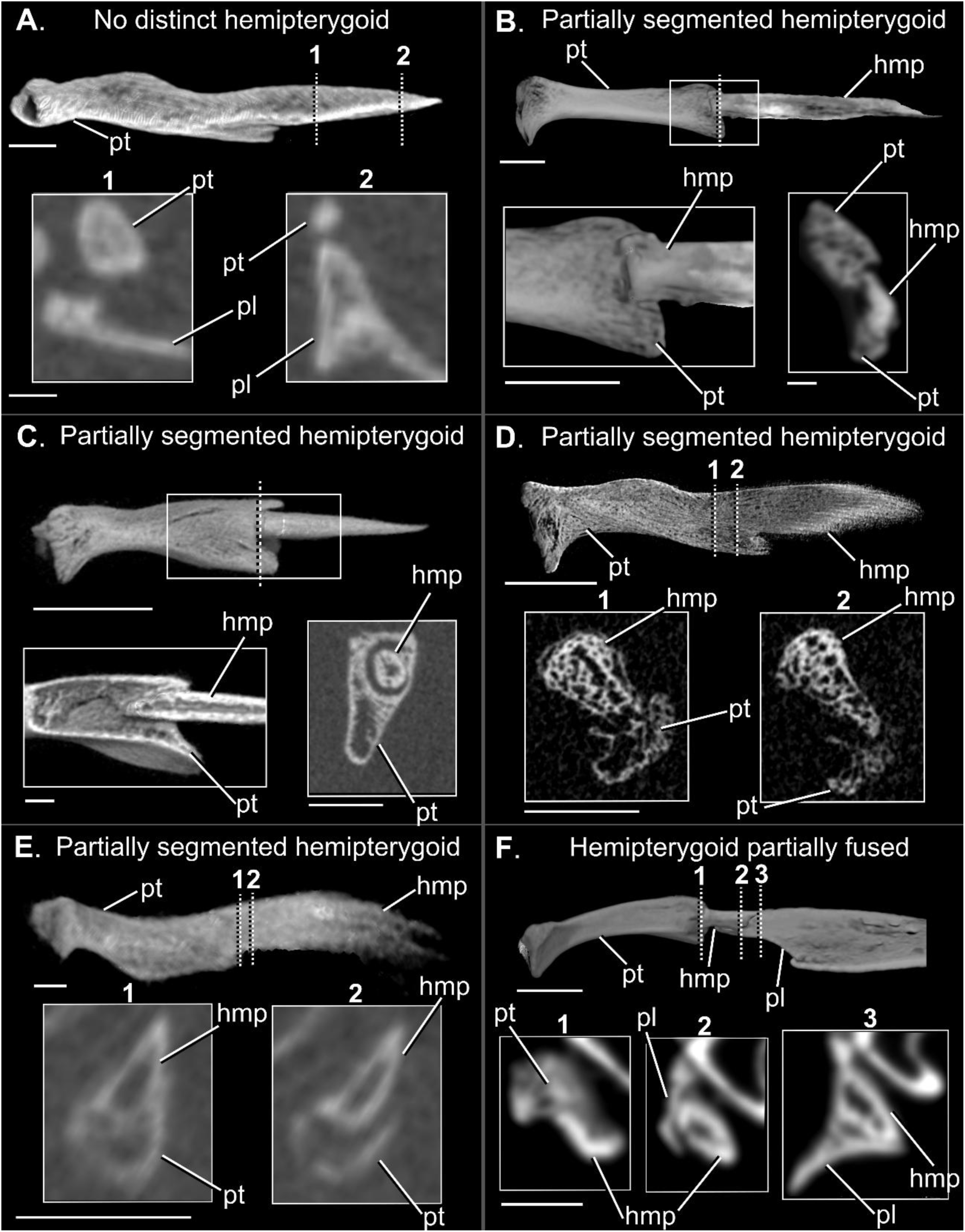
Identification of the hemipterygoid based on observation of 3D models and μCT cross sections. **A**. lateral view of the left pterygoid (reflected) of embryonic *Pteroglossus viridis* (Neoaves: Ramphastidae); **B**. lateral view of the right pterygoid and hemipterygoid of juvenile *Nycticorax nycticorax* (Neoaves: Ardeidae); **C**. lateral view of left pterygoid and hemipterygoid (reflected) of hatchling-stage *Spheniscus demersus* (Neoaves: Spheniscidae); **D**. lateral view of right pterygoid and hemipterygoid of juvenile *Morus bassanus* (Neoaves: Sulidae); **E**. lateral view of right pterygoid and hemipterygoid of hatchling-stage *Columba livia* (Neoaves: Columbidae); **F**. oblique lateral view of right pterygoid, hemipterygoid and part of the palatine of juvenile *Egretta garzetta* (Neoaves: Ardeidae). Dashed vertical lines in images correspond to cross-sections (coronal/oblique coronal) and white boxes on the 3D rendering denote regions displayed as separate inset images, which in the case of *Spheniscus* represents a 3D clipping plane illustrating internal morphology. The unlabelled element in cross sections for part F represent the parabasisphenoid. Abbreviations: hmp, hemipterygoid; pl, palatine; pt, pterygoid. Scale bars: A. 3D model and cross section scales represent 0.5 mm; B. 3D model scale (including inset image) represent 2.5 mm and cross section scale represents 0.5 mm; C. 3D model scale represents 2.5 mm, cross section scale represents 1.25 mm and scale by the inset image represents 0.5 mm; D. 3D model scale represents 2.5 mm and the cross section scale represents 1.25 mm; E. 3D model scale represents 0.5 mm and cross section scale represents 1.25 mm; F. 3D model scale represents 2.5 mm and cross section scale represents 1 mm.

## 3. RESULTS

Below we describe the morphology and observed ontogenetic patterns across our entire sample, grouped by major clade (Palaeognathae, Anseriformes, Galliformes, and Neoaves). In each case, descriptions are organised at a taxonomic level appropriate to summarise major patterns of variation and to convey the degree of intraclade variation observed; in cases where all surveyed family-level clades exhibit strikingly divergent patterns (e.g., Palaeognathae, Anseriformes) family-level patterns are described separately, but in cases where several or all family-level clades show broadly congruent patterns (e.g., non-megapodiid Galliformes, Neoaves), family-level patterns are described together.

### PALAEOGNATHAE

We examined six immature and four comparatively mature palaeognath specimens representing Tinamidae (tinamous), Casuariidae (cassowaries and emu), Apterygidae (kiwis), and Struthionidae (ostriches) (suppl. table 1). A larger sample of comparatively mature specimens was consulted for comparative purposes (suppl. table 2). Additionally, we examined a subadult subfossil specimen of the moa *Megalapteryx didinus*. All the surveyed palaeognath family-level clades exhibit major differences in both the morphology of the pterygoid-palatine complex (PPC) and the degree of post-hatching ontogenetic change it undergoes, and as such are described separately below (figs. 5-7).

**Figure 5:**
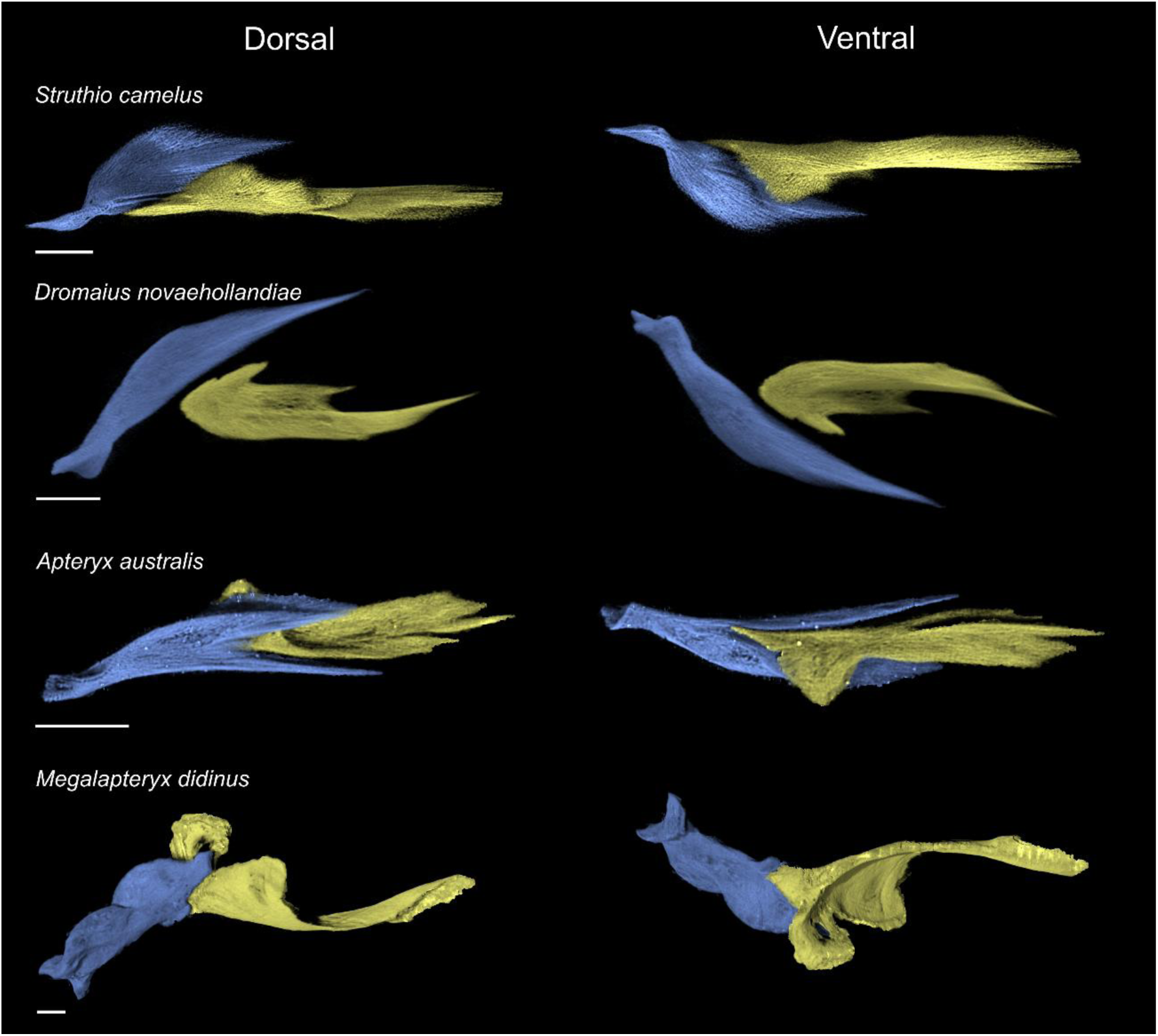
Comparative morphology of the palatine (yellow) and pterygoid (blue) in articulation in select immature palaeognaths, figured for the right side of the skull (*Struthio*, *Apteryx* and *Megalapteryx* elements are reflected) (part 1). ‘Dorsal’ and ‘ventral’ orientations are with respect to the skull. All scale bars represent 2.5 mm.

**Figure 6:**
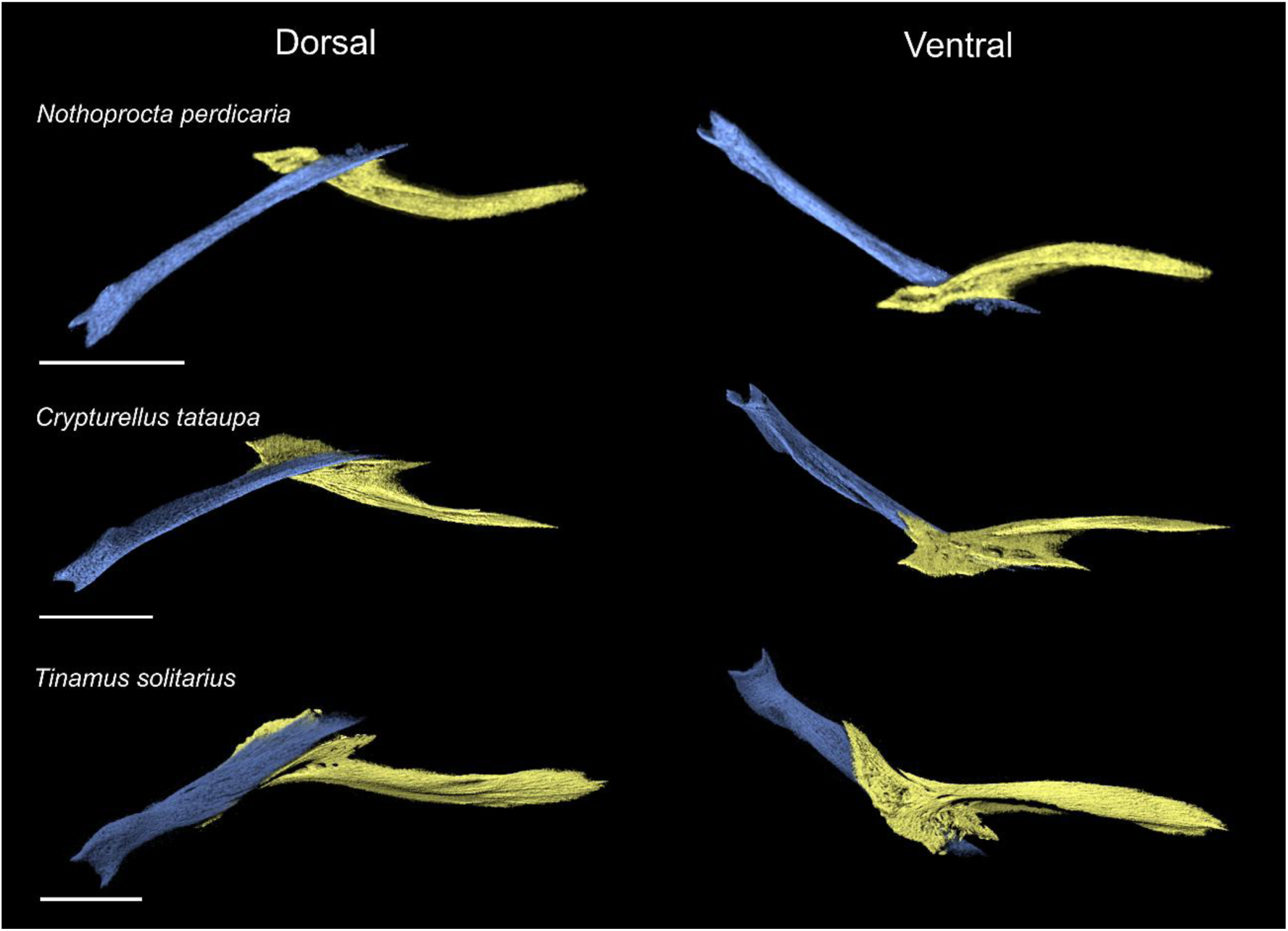
Comparative morphology of the palatine (yellow) and pterygoid (blue) in articulation in select immature palaeognaths, figured for the right side of the skull (part 2). ‘Dorsal’ and ‘ventral’ orientations are with respect to the skull. All scale bars represent 2.5 mm.

**Figure 7:**
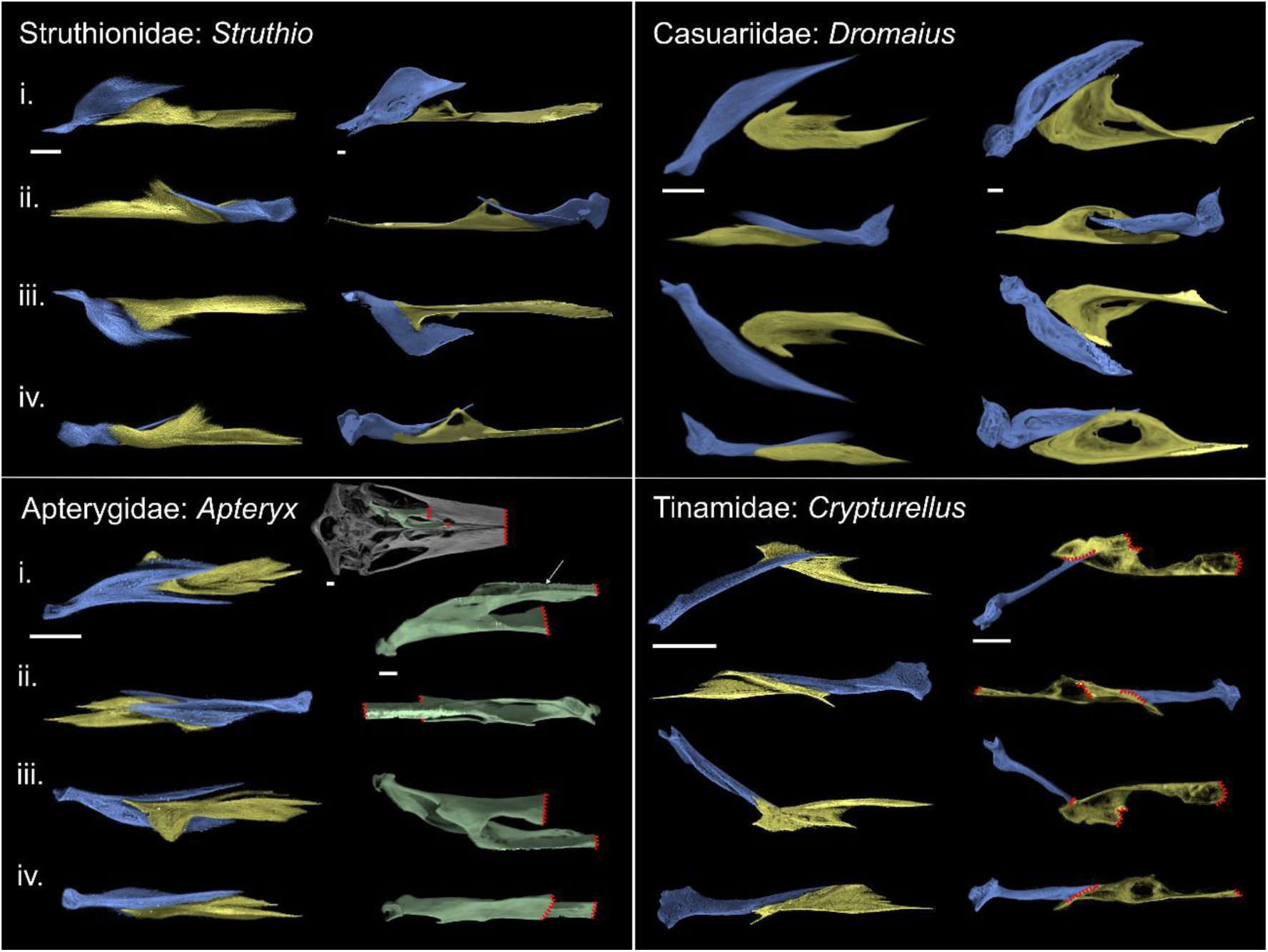
Comparison of immature (left columns of each ‘box’) and mature (right columns of each ‘box’) palaeognath palatine (yellow) and pterygoid (blue) morphology, figured for the right side of the skull (immature and mature *Struthio* and immature and mature *Apteryx* elements are reflected). Red dotted lines represent areas where bone margins cannot be precisely delineated due to fusion. Struthionidae pairing: *Struthio camelus* (Common Ostrich) immature and subadult; Casuariidae pairing: *Dromaius novaehollandiae* (Emu) immature and subadult; Apterygidae pairing: *Apteryx australis* (Southern Brown Kiwi) immature and subadult; Tinamidae pairing: *Crypturellus tataupa* (Tataupa Tinamou) immature and adult. Due to extensive fusion in adult *Apteryx*, we figure the palatine and pterygoid as a single complex (green), which also includes part of the vomer (arrowed). The position of this complex is indicated on the skull in ventral view. i. - iv. represent the following orientations: i.- dorsal, ii.- medial, iii.- ventral and iv.- lateral (all with respect to the skull). All scale bars represent 2.5 mm.

#### STRUTHIONIDAE

##### Immature

The caudal region of the palatine of near-hatching embryonic *Struthio camelus* is already well ossified and bears a pterygoid process (*processus pterygoideus*) that is clearly distinct from the rest of the palatine (fig. s1). The pterygoid process is mediolaterally wide and includes a flat, dorsocaudally oriented shelf forming most of the pterygoid’s articular surface, which is subtriangular in dorsal view. The lateral margin of the pterygoid process extends into a dorsoventrally deep and mediolaterally narrow caudally directed flange, which tapers caudally into a caudodorsally pointed tip. Although the palatine and pterygoid are not yet in contact, presumably as a result of incomplete ossification, the pterygoid appears to lie flat over the subtriangular contact surface, with its lateral surface contacting the entire caudolateral flange of the pterygoid process (fig. 5).

The wing-shaped pterygoid extends into a pronounced and tapered rostral projection, which terminates in a pointed rostral tip (fig. s2). This rostral projection is rostromedially directed and extends medial to the body of the palatine (fig. 5), exhibiting a marked dorsal deflection towards its rostral terminus. Caudal to this projection, the main body of the pterygoid is mediolaterally wide and dorsoventrally compressed; its dorsal surface is slightly concave while its ventral surface is moderately convex. The lateral margin of the pterygoid rises into a dorsoventrally deep and mediolaterally narrow ridge, which extends caudally into an elongated projection. This lateral ridge is aligned with the orientation of the corresponding lateral flange extending from the pterygoid process of the palatine, which it contacts (fig. 5).

##### Mature

The near-mature specimen of *Struthio camelus* examined is a subadult, and although certain sutures are not fully obliterated (such as the palatine-pterygoid contact), the specimen is of almost adult size and illustrates the mature condition well. Comparison of the near-mature specimen with the near-hatching specimen documents a limited degree of ontogenetic change across the caudal region of the palatine and the rostral portion of the pterygoid, and the relationships between these elements is mostly unchanged (fig. 7).

In the subadult, the pterygoid and palatine are fully in contact, with the pterygoid process of the palatine wrapping around the lateral and ventral surfaces of the rostrolateral margin of the pterygoid. The dorsoventrally narrow and dorsocaudally oriented shelf of the pterygoid process extends further medially than in the more immature specimen examined, such that it becomes mediolaterally rather than rostrocaudally aligned and is more rectangular than subtriangular in shape. The lateral flange extending from the pterygoid process of the subadult is more rounded than caudally tapered and contacts the lateral surface of the pterygoid (fig. 7).

In the subadult, the rostral projection of the pterygoid is more strongly dorsally deflected, with its tip appearing slightly more rounded than in the embryonic specimen examined. In contrast with the less mature specimen, this projection does not lie immediately medial to the body of the palatine, such that a large gap exists between both structures. The wing-like surface extending caudally from this rostral projection is relatively mediolaterally wider, such that the pterygoid process of the palatine only wraps around its ventrolateral margin, rather than around much of its ventral surface as in the younger specimen, and has a marked and well-defined dorsal concave depression. This wing-like surface is also more clearly differentiated from the thickened and dorsoventrally deeper ridge forming the lateral margin of the pterygoid than in the younger specimen.

#### CASUARIIDAE

##### Immature

The palatine of the examined immature *Dromaius novaehollandiae* does not bear a pronounced, caudally directed pterygoid process, in contrast to the more immature *Struthio camelus* (fig. s1). Instead, the caudalmost end of the palatine bears a slightly pointed, dorsoventrally compressed tip, that is not distinct from the rest of the palatine. The dorsal and ventral surfaces of the caudal region of the palatine are smooth and flat. The palatine and pterygoid are not in direct contact (as is also the case for the more immature *Struthio*), presumably as a result of incomplete ossification (fig. 5). Unlike in *Struthio*, the relative position of these elements indicates that the two bones are positioned side-by-side, rather than with the pterygoid overlying the palatine. Moreover, the medial surface of the pterygoid process is slightly longer and thicker than the lateral surface, indicating the potential contact zone with the pterygoid. Similar to the condition in *Struthio*, the pterygoid is wing-shaped and exhibits a rostromedially-directed rostral projection (fig. 5) that narrows mediolaterally towards its rostral end (fig. s2). The wing-shaped portion of the pterygoid represents a much larger proportion of the entire length of the element than in *Struthio*, yet it is relatively mediolaterally narrower. This region is dorsoventrally compressed, with a somewhat concave dorsal surface; its lateral margin is straight and its medial margin is curved. Caudal to the rostromedially directed process, the medial margin of the pterygoid is slightly thickened, outlining the contact zone with the palatine, yet, unlike in *Struthio,* it only becomes a dorsoventrally deep ridge at the caudalmost end of the pterygoid.

##### Mature

The older specimen of *Dromaius novaehollandiae* examined is not fully osteologically mature, instead corresponding to an older immature or subadult, yet although certain sutures are not fully obliterated, it is of almost adult size and illustrates the adult condition well. There is substantial ontogenetic change between the immature and comparatively mature *Dromaius* specimens, particularly regarding the shape of the palatine, although the general arrangement of both elements does not change (fig. 7).

Compared to the immature *Dromaius*, the palatine of the more mature specimen is substantially wider mediolaterally, and its caudalmost portion extends into a more distinct pterygoid process, though it lacks the mediolaterally wide contact shelf observed in *Struthio*. Instead, the pterygoid process of *Dromaius* is subtriangular, caudally pointed and slightly medially deflected. The dorsal region of the pterygoid process is somewhat convex, while the ventral portion is flat. In contrast to the immature, the lateral margin of the pterygoid process is thicker than the medial margin, but the latter extends into a small ridge that forms much of the medial surface of the palatine. The rostralmost portion of this ridge, close to the rostromedial process of the palatine, includes a short side-to-side longitudinal contact between the palatine and the pterygoid. This rostrally situated contact between both bones contrasts with the condition in *Struthio* and most other surveyed taxa, where it occurs at or near the pterygoid process (fig. 7).

The pterygoid of the more mature *Dromaius* is very reminiscent of the general condition of the immature, although its wing-like rostral portion is proportionally wider, and the dorsally curved rostral projection is medially deflected. The dorsal surface of the pterygoid is flatter than in the immature, yet it exhibits a previously absent rounded pneumatic fenestra towards its caudal end, and the ventral surface of the element is more strongly convex. The lateral margin of the pterygoid is narrow and sharp across the rostralmost third of the element, but becomes rounded and dorsoventrally thicker at the point where the pterygoid and the palatine contact; this contact is short, and the lateral margin of the pterygoid narrows again caudal to this point.

#### APTERYGIDAE

##### Immature

The palatine of the immature *Apteryx australis* extends into a distinct pterygoid process which, similar to the condition in *Struthio*, bears a mediolaterally wide articular shelf and a prominent caudolateral flange (fig. s1). The articular shelf is markedly medially expanded and subtriangular in dorsal view, and its dorsal surface is slightly concave. In contrast to the condition in *Struthio,* the caudolateral flange of *Apteryx* is dorsoventrally low and narrows in mediolateral width caudally, terminating in a caudally directed, pointed tip. The pterygoid approaches both structures (fig. 5), and the caudolateral flange of the pterygoid process of the palatine cleanly fits within a marked groove on the ventral surface of the pterygoid, while the wider shelf is overlain by the ventral surface of the medial process of the pterygoid.

The rostral portion of the pterygoid forks into distinct lateral and medial processes, with both processes tapering rostrally to form pointed tips (fig. s2). Both processes are separated by a narrow ventral groove. The medial process of the pterygoid is deeply concave medially, and laterally is convex and angular. This process overlies the articular shelf of the palatine, and has a close contact with the caudolateral flange of the palatine, which slightly wraps over the curved ventrolateral surface of the medial process, inserting into the groove separating both processes of the pterygoid. The lateral process of the pterygoid is blade-like, and dorsoventrally narrower than the medial process. It lacks any curvature and extends along most of the lateral edge of the palatine without contacting it.

##### Mature

The examined subadult specimen of *Apteryx australis* exhibits an extreme degree of fusion, preventing the precise delineation of the contact between the palatine and pterygoid. As a result, we figure the palatine and pterygoid as a single complex, which also includes the vomer, which we label the ‘palatine-pterygoid-vomer complex’ (fig. 7). The resulting structure is strongly bifurcated, with two rostral prongs. The medial prong of the complex is clearly formed by the merger of most of the pterygoid process of the palatine and the ventral surface of the medial process of the pterygoid, as well as fusion of the rostralmost portion of both structures to the caudal portion of the vomer. The lateral prong of the complex, in contrast, is formed mostly by the main body of the palatine and the lateral process of the pterygoid, which, unlike in the immature specimen, not only contacts but fully fuses with the lateral surface of the palatine.

#### MEGALAPTERYGIDAE

In addition to extant palaeognaths, we also examined the palate elements of the holotype of the Upland Moa, *Megalapteryx didinus* ((Owen, 1883); (Rawlence et al., 2013)), to more fully characterise the diversity of palatal morphologies in Palaeognathae. Although no precise ontogenetic assessment of this specimen is available, its large size—close or equal to that of previously reported adult specimens of *Megalapteryx* (Worthy & Scofield, 2012) and the closed sutures between several skull elements suggest that it corresponds to a nearly mature individual. However, in this specimen the palatine, pterygoid, and vomer remain unfused to one another, unlike in published adult specimens of *Megalapteryx* (Worthy & Scofield, 2012) and we therefore consider it likely that it represents a subadult individual that was very close to skeletal maturity (fig. 5).

The pterygoid process of the palatine forms a distinct tapered and pointed caudal projection from the body of the palatine, with a rugose caudomedially aligned subtriangular dorsal articular surface (fig. s1). This surface is divided mediolaterally by a low ridge, which defines two smaller and slightly concave lateral and medial depressions both of which are contacted by the pterygoid; the medial depression is rostrocaudally longer and extends further rostrally than the lateral depression. Medial to the pterygoid process, and extending rostrally to it, the palatine of *Megalapteryx* exhibits a bizarre disc-shaped lobe, pierced by a large fenestra. The rostral portion of this lobe contacts the vomer and lobes of both palatines meet medially, yet this structure does not participate in the pterygoid-palatine contact, despite being medially adjacent to it.

As in *Struthio* and *Dromaius*, the main body of the pterygoid is wing-shaped, extending rostrally into a pointed rostral projection (fig. s2). The rostral tip of this projection is strongly tapered and slightly dorsally deflected, and does not contact the palatine in the examined specimen. Beyond these superficial similarities, however, the pterygoid of *Megalapteryx* is strikingly divergent with respect to that of *Struthio* and *Dromaius*. In *Megalapteryx* the wing-like portion of the pterygoid represents less than half of the total length of the element, while it comprises the majority of its rostrocaudal length in *Struthio* and *Dromaius*. Furthermore, in contrast to these taxa, the wing-like portion of the *Megalapteryx* pterygoid appears to be twisted, such that it is dorsoventrally deep and mediolaterally compressed rather than mediolaterally wide and dorsoventrally shallow, with a strongly concave medial surface and convex lateral surface. The pterygoid contacts the palatine along its narrow ventral margin, which exhibits two low, rostrocaudally elongate ridges separated by a shallow groove; these cleanly match the pterygoid process of the palatine which bears two corresponding concave articular surfaces separated by a low ridge.

#### TINAMIDAE

##### Immature

The pterygoid process of the palatine of all examined immature tinamous forms a flat or slightly concave surface (fig. s1), yet the surveyed taxa differ in the variable branching of this structure. In *Crypturellus tataupa,* the pterygoid process bifurcates into two pointed dorsolateral and ventromedial branches separated by a subtriangular notch. The pterygoid forms a very long rostrocaudal overlap with the palatine across the dorsolateral branch of the pterygoid process, extending further rostrally onto the body of the palatine; however, the pterygoid and palatine are not fully in contact across the entire area of overlap (fig. 6). The ventromedial branch (identified as the caudomedial process by Bertelli et al. (2014)), in contrast, does not participate in the pterygoid-palatine contact.

The pterygoid process of immature *Nothoprocta perdicaria* exhibits a structure equivalent to the ventromedial branch of *Crypturellus,* which does not participate in the pterygoid-palatine contact (fig. 6). No dorsolateral branch is present in *Nothoprocta,* and the pterygoid only overlies the palatine rostral to the pterygoid process where they are not in contact. Contrastingly, *Tinamus solitarius* instead lacks a ventromedial branch, while an elongated dorsolateral branch is present, bearing a long contact with the pterygoid, mirroring the long overlap between both elements in *Crypturellus* (fig. 6).

The pterygoid of all surveyed immature tinamous is elongate and rod-like (fig. s2). The rostralmost portion of the pterygoid is slightly compressed mediolaterally, yielding a blade-like appearance, with a moderately concave medial surface extending into a narrow and tapered rostral projection which is slightly laterally deflected. The ventral margin of this rostral region of the pterygoid contacts the matching articular surface of the palatine (including the dorsolateral branches of the pterygoid process in *Crypturellus* and *Tinamus*), with this contact extending towards the rostromedial process of the palatine. The rostralmost portion of this projection contacts the dorsal surface of the underlying vomer.

##### Mature

Beyond fusion between the palatine and the pterygoid, the examined comparatively mature tinamids (suppl. tables 1 and 2) generally show little ontogenetic change with respect to their corresponding immatures. The dorsolateral branch of the pterygoid process present in immature *Crypturellus* is completely fused to the pterygoid in the adult, such that both elements form a dorsoventrally deep, narrow flange lacking clear boundaries between its component elements (fig. 7). In both adult *Crypturellus* and *Nothoprocta*, the ventromedial branch of the pterygoid process is mediolaterally wider than in the immature, with a rounder medial edge; its caudal end terminates in a slightly laterally hooked point, but otherwise remains unchanged in its size and proportions. In contrast to the immature *Nothoprocta*, whose pterygoid process only bears a ventromedial branch, the surveyed subadult and adult representatives also exhibit a broad ascending dorsolateral branch similar to that of *Crypturellus*, which in the adult completely fuses with the ventral margin of the pterygoid. The pterygoid of adult *Tinamus* also develops a marked ventromedial branch, while the immature only exhibits a dorsolateral branch. Although the order in which these branches appears is variable among surveyed tinamous (dorsolateral first in *Tinamus*, ventromedial first in *Nothoprocta*), all surveyed adult tinamous (see suppl. tables 1 and 2) exhibit both dorsolateral and ventromedial branches of the pterygoid process.

The rostral projection of the pterygoid completely fuses to both the palatine and the vomer in adult tinamous, such that the rostrally tapering blade-like morphology of immature tinamous is no longer observable. Once fused, the pterygoid forms a pronounced dorsoventrally deep ridge extending across much of the rostrocaudal length of the palatine-pterygoid complex, which maintains the lateral curvature observed in immature tinamou pterygoids.

### GALLOANSERAE

#### ANSERIFORMES

We directly examined eight immature and three adult specimens representing Anhimidae and Anatidae (suppl. table 1). A larger sample of adult Anseriformes across both families (as well as Anseranatidae) were included for comparative purposes (suppl. table 2). We found substantial differences in the morphology and the degree of ontogenetic change between Anhimidae and Anatidae, which are therefore described separately below (figs. 8-10).

**Figure 8:**
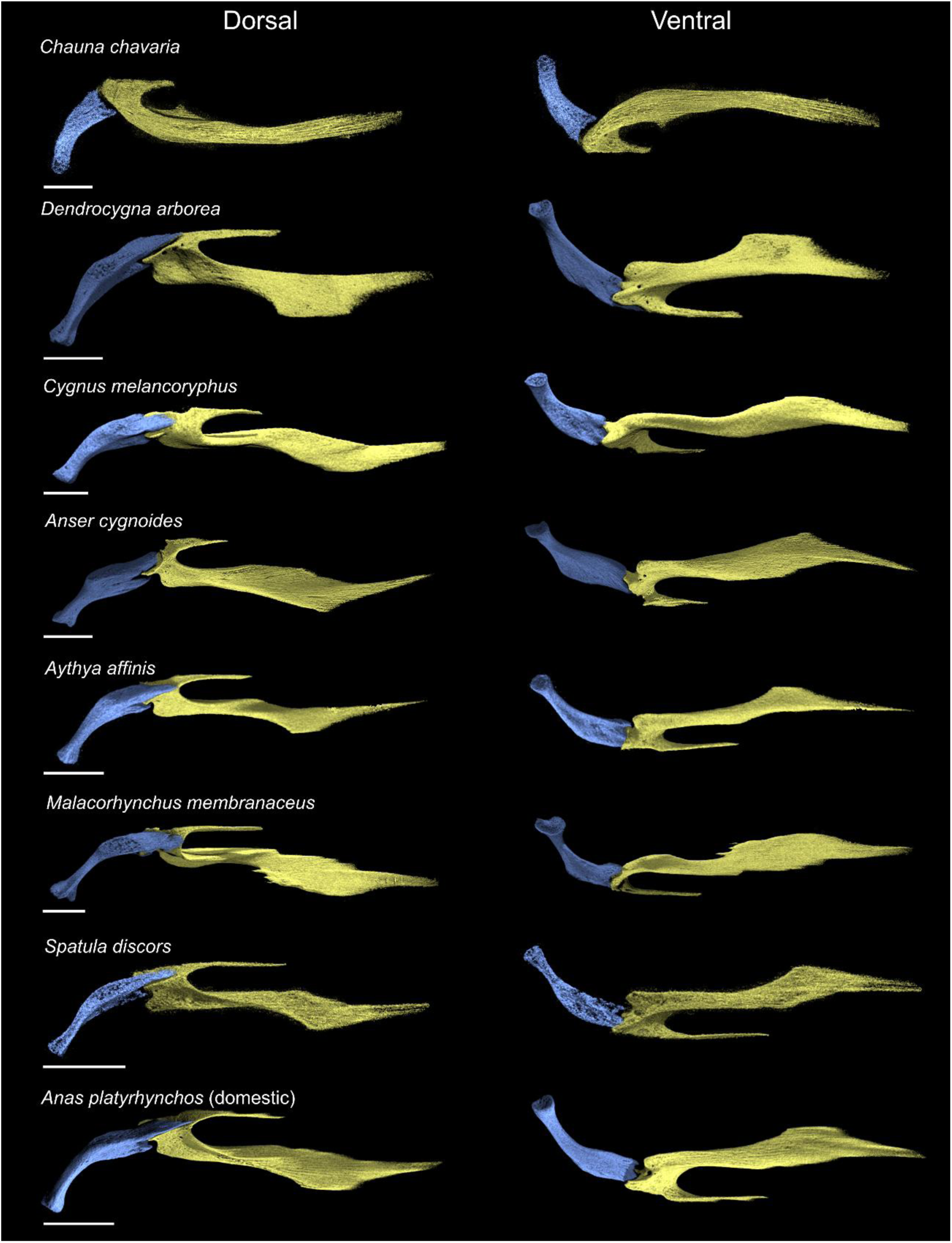
Comparative morphology of the palatine (yellow) and pterygoid (blue) in articulation in select immature anseriforms, figured for the right side of the skull (*Chauna*, *Anser*, *Aythya*, *Malacorhynchus*, *Spatula* and *Anas* elements are reflected). ‘Dorsal’ and ‘ventral’ orientations are with respect to the skull. All scale bars represent 2.5 mm.

**Figure 9:**
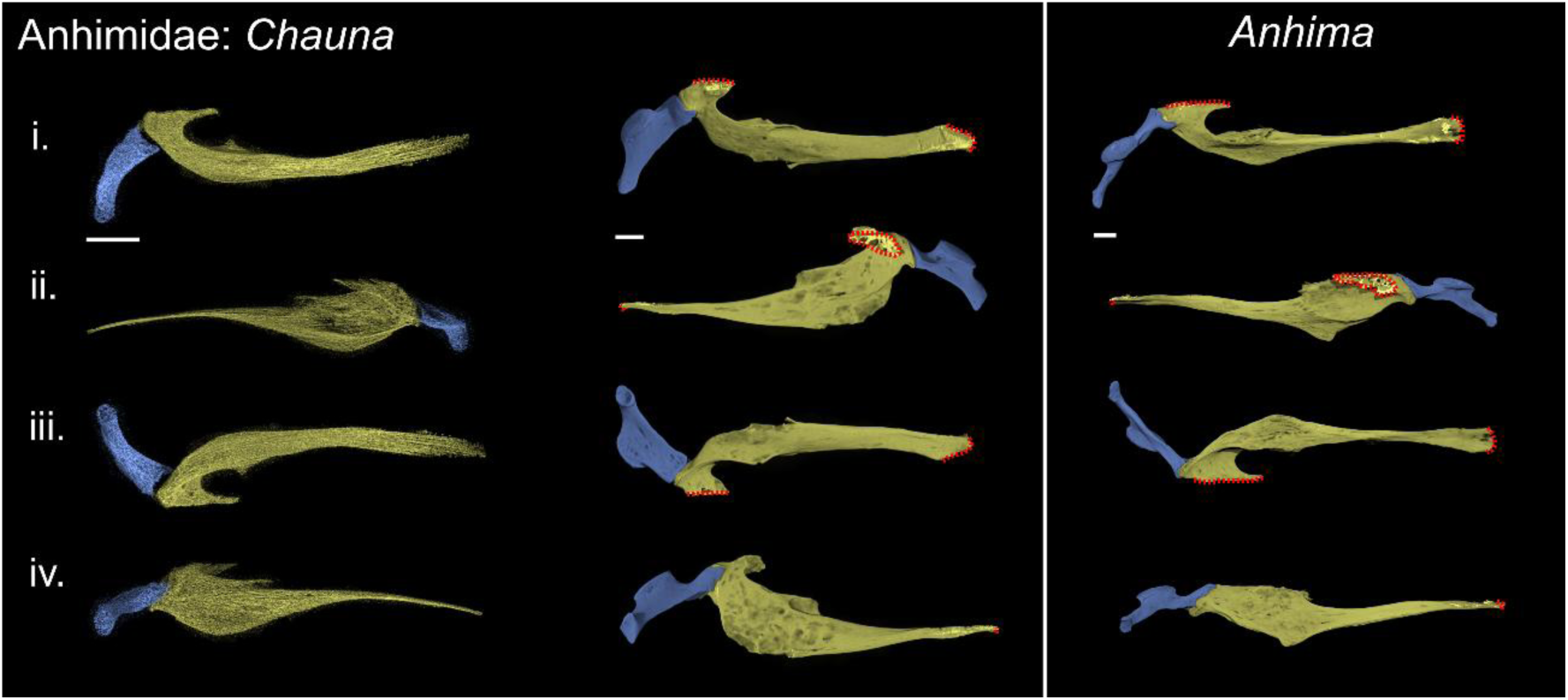
Comparison of Anhimidae palatine (yellow) and pterygoid (blue) morphology, figured for the right side of the skull (*Chauna* immature and mature elements are reflected). Left box, *Chauna chavaria* (left column – immature); *Chauna chavaria* (right column – mature), right box mature *Anhima cornuta*. Red dotted lines represent areas where bone margins cannot be precisely delineated due to fusion. i.- iv. represent the following orientations: i.- dorsal, ii.- medial, iii.- ventral and iv.- lateral (all with respect to the skull). All scale bars represent 2.5 mm.

**Figure 10:**
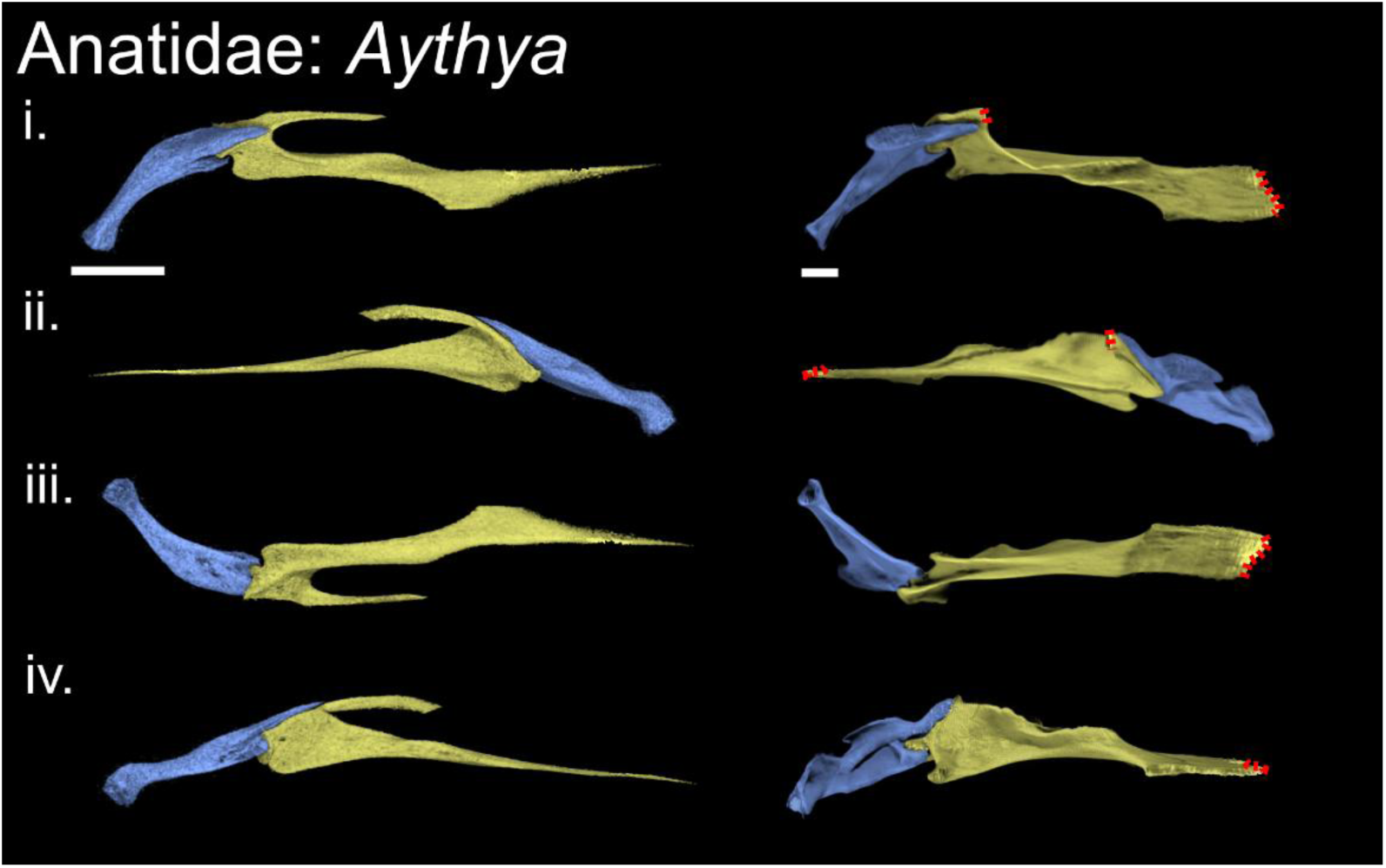
Anatidae, comparison of immature *Aythya affinis* (Lesser Scaup; left column) and mature *Aythya ferina* (Common Pochard; right column) palatine (yellow) and pterygoid (blue), figured for the right side of the skull (immature *Aythya* elements are reflected). Red dotted lines represent areas where bone margins cannot be precisely delineated due to fusion. i.- iv. represent the following orientations: i.- dorsal, ii.- medial, iii.- ventral and iv.- lateral (all with respect to the skull). All scale bars represent 2.5 mm.

#### ANHIMIDAE

##### Immature

The caudal region of the palatine of the Screamer *Chauna chavaria* is incompletely ossified and does not bear a caudally projected pterygoid process clearly distinct from the rest of the palatine (fig. s3). Instead, at the caudalmost end is a poorly defined, short and sub-rounded protuberance. The short rostromedial process of the palatine is aligned with the caudal end of the element, but the dorsoventrally deep main body of the palatine, including the elongated rostral process, is laterally displaced and curved, defining a convex lateral outline of the palatine (fig. s3).

The pterygoid of the immature *Chauna* is simple in shape, with a broadly cylindrical outline, widening mediolaterally towards its rostral end (fig. s4). The rostral region of the element is incompletely unossified, yet an incipient cotyle for the pterygoid process of the palatine is apparent, forming a poorly defined shallow depression. The dorsolateral margin of the rostral end of the pterygoid is slightly rostrally projected and tapered, extending along (though not contacting) a small portion of the caudal palatine. Although poorly ossified and diffuse, an incipient facet for the basipterygoid process appears to be developed on the dorsomedial surface of the pterygoid, immediately caudal to the rostral end of the element. The incomplete ossification of both palatine and pterygoid results in a noticeable space between both palatal elements which are therefore not in direct contact (fig. 8).

##### Mature

In contrast to the poorly ossified condition of the immature specimen, the adult *Chauna chavaria* has a very short, and well-defined pterygoid process, with a sub-triangular caudal end defining a dorsoventrally deep condyle that tapers into a short, narrow ventrolateral projection (fig. 9). The ventromedial surface of the pterygoid process forms a deep undercut or depression just rostral to this projection. Remarkably, the pterygoid process of *C. torquata* is much more robust and rounded, lacking any ventromedial projection or undercut. The broad morphology of the palatine shows a moderate degree of change between immature and adult *C. chavaria*, with a robust rostromedial process that appears to be co-ossified with the vomer, and a notable change in the shape of the caudolateral corner of the palatine, which is rounded in immatures but becomes squared in adults (fig. 9). Interestingly, the morphology of both aspects in the adult *C. torquata* is more reminiscent of that of the immature *C. chavaria*.

The rostral portion of the pterygoid, which is poorly defined in the immature *C. chavaria*, develops into a rostral projection that includes the cotyle for the palatine at its rostral tip. This rostral projection is short and mediolaterally compressed; a pronounced ridge runs along its lateral surface, extending rostrally slightly further than the rest of the rostral projection, forming a short tapering triangular tip at its dorsolateral end which contacts the palatine (see below). The only major observed ontogenetic change in the pterygoid corresponds to the relative position of the facet for the basipterygoid process, which, contrary to the condition in the immature *C. chavaria*, is not adjacent to the rostral end of the pterygoid, and is instead much more caudally situated (fig. 9). The entire rostral end of the rostral projection forms a mediolaterally narrow but dorsoventrally deep concave articular cotyle that wraps around the lateral and caudal surfaces of the pterygoid process of the palatine. As in the immature *C. chavaria*, the dorsolateral margin of the rostral projection extends into a short, tapering triangular process which overlaps the dorsolateral surface of the pterygoid process of the palatine. The morphology of the pterygoid of adult *C. torquata* is quite divergent from that of either immature and adult *C. chavaria*: the rostral projection of the pterygoid shows a marked mediolateral constriction at its caudal end, rather than being of constant mediolateral width across its length, and is proportionally shorter, such that the facet for the basipterygoid process is much more rostrally situated than in the adult *C. chavaria*.

Furthermore, the rostral projection of *C. torquata* extends over both the dorsolateral and ventromedial surfaces of the pterygoid process of the palatine, developing a distinct though short ventromedial tip that wraps around the ventral palatine, entirely absent in either immature or adult *C. chavaria*. Finally, the dorsolateral margin of the rostral pterygoid projection of *C. torquata* is short and rounded, rather than tapered and pointed as in *C. chavaria*. A similar mediolaterally compressed rostral projection of the pterygoid bearing a rostral cotyle is also present in a mature specimen of *Anhima cornuta* (fig. 9), although it is substantially longer than in either *Chauna* species, representing approximately half of the entire length of the pterygoid. Despite the pterygoid of both taxa being otherwise quite distinct, this rostral projection in *Anhima* resembles that of *C. chavaria* more closely, with no ventromedial overlap over the palatine like that of *C. torquata*, and with a pointed dorsolateral margin wrapping over the pterygoid process. Importantly, the condition of this pointed dorsolateral overlap in *Anhima,* which is mediolaterally narrower and rostrocaudally much more elongated than in *Chauna*, is reminiscent of the shape, position and articulation of the rostral projection of the pterygoid in Anseres (see below).

#### ANATIDAE

##### Immature

All surveyed immature anatids exhibit a similar morphology in both the caudal region of the palatine and the rostral portion of the pterygoid, which, in contrast to the condition observed in *Chauna*, is more completely ossified and better defined in all sampled specimens (fig. 8). The pterygoid process is relatively short rostrocaudally, dorsoventrally flattened, and deflected medially with respect to the main body of the palatine and its rostral process, roughly aligned with the rostromedial process. The caudal end of the pterygoid process bifurcates into a pair of projections, which are incipiently developed in *Aythya affinis*, *Spatula discors* and *Malacorhynchus membranaceus*, and marked and well-defined in *Dendrocygna arborea*, *Cygnus melancoryphus* and *Anas platyrhynchos*; the pterygoid process appears not to bifurcate in the immature *Anser cygnoides* (fig. s3). Each projection extends from the lateral and medial margins of the pterygoid process, respectively, defining a wide concavity between them of variable interspecific depth (e.g., deep in *Aythya*, shallow in *Cygnus*). The lateral projection is shorter, slightly more dorsally situated and generally more rounded, and the medial projection is dorsoventrally compressed, slightly more ventrally positioned, and extends further caudally.Both projections participate in the articular contact for the pterygoid. The rounded, laterally positioned condyle-like projection fits within the palatine cotyle of the pterygoid, while the more blade-like medial projection extends beneath the ventral surface of the rostral pterygoid. The concavity between both projections continues rostrally, forming an articular surface for the rostral projection of the pterygoid, which extends along the dorsomedial margin of the caudal palatine until it reaches the base of the rostromedial process (fig. s3). This articular surface is mostly flat rostrally but becomes progressively more concave towards the bifurcated caudal end of the pterygoid process. On the lateral surface of the palatine, all examined immature anatids (apart from *Cygnus*) bear a well-defined caudolateral corner which is less caudally projected than the pterygoid process in all cases, yet is variably projected across the surveyed taxa, ranging from rounded and small in *Dendrocygna* to squared and elongated in *Malacorhynchus*.

The dorsal surface of the rostral pterygoid extends into a pronounced rostral projection which represents between ∼ 1/4 (*Cygnus, Anser*) and ∼ 1/3 (*Dendrocygna, Aythya, Spatula, Malacorhynchus* and *Anas*) of the total pterygoid length (figs. s4, s5). This rostral projection is blade-like and tapers rostrally. It is dorsoventrally compressed, dorsomedially flat and ventrolaterally flat to concave, overlapping much of the caudal portion of the palatine where it covers its dorsomedial articular surface. An incipient facet for the basipterygoid process extends caudal to the rostral projection and across much of the dorsomedial surface of the pterygoid in some immature anatids (*Malacorhynchus*), but is absent or less defined in others (*Dendrocygna, Cygnus*). Ventrolateral to the rostral projection, the rostral surface of the pterygoid is dominated by a deep, concave depression which is variably defined in the surveyed immature anatids (e.g., it is well delimited in *Malacorhynchu*s, but poorly defined in *Spatula*). This depression corresponds to the cotyle for the palatine and articulates with the lateral projection of the pterygoid process. Although some degree of post-mortem rotation cannot be excluded in the examined immatures, the cotyle appears to be open dorsally and laterally, but is bounded medially by the rostral projection and ventrally by a pronounced lip that wraps around the lateral projection of the pterygoid process; this lip is very short in *Dendrocygna* but longer in the other immature anatids examined (figs. s4, s5). No distinct hemipterygoid is discernible in any of the surveyed immature anatid specimens.

##### Mature

The sampled adult anatids (suppl. tables 1 and 2) exhibit a limited degree of ontogenetic change across the caudal region of the palatine and the rostral portion of the pterygoid. The spatial relationship between the elements does not change ontogenetically, but both exhibit a noticeable degree of rotation. The bifurcate morphology of the pterygoid process does not change in the adult anatids, but both projections become mediolaterally aligned rather than being noticeably dorsoventrally offset; this ontogenetic change is best exemplified in *Aythya* in our sample (fig. 10). The concavity between both projections, as in the immatures, extends rostrally into a long, flat, dorsomedial contact surface for the rostral projection of the pterygoid. The caudolateral corner in adult anatids is generally distinct, squared, and caudally projected, compared with the rounded caudolateral corner of most immature anatids. It is only slightly less caudally projected than the pterygoid process.

The rostral portion of the pterygoid similarly maintains its general morphology across ontogeny, though the shape of the rostral projection is ontogenetically variable. The relatively broad, tapering, blade-like rostral projection of immature anatids becomes roughly cylindrical, dorsally convex and ventrally flat, with a marked dorsal deflection across its length. It bears a rounded (e.g., *Aythya*) to sharply pointed (e.g., *Anas*) rostral tip. Indeed, this rostral projection of adult anatids resembles the rostral projection of the adult anseranatid *Anseranas* (suppl. table 2). The basipterygoid process articular facet is poorly developed in immature anatids, but becomes a distinct, well-defined medially projected facet in adult anatids positioned immediately caudal to the rostral projection (as also present in *Anseranas*).

The variably ossified cotyle for the palatine of immature anatids is a well-defined cup-shaped concavity in all examined adults, receiving the rounded lateral projection of the pterygoid process of the palatine. The dorsoventrally flattened ventral lip of the cotyle is slightly more rostrally projected in adult anatids, wrapping around the ventral surface of the aforementioned palatine condyle. The ventromedial surface of the pterygoid immediately caudal to the rostral projection and medial to the pterygoid cotyle bears a subtriangular contact surface for the flattened, medially positioned projection of the pterygoid process of the palatine; the extent of this contact is variable (e.g., short in *Aythya*, long in *Anas*). Although the arrangement of this contact is similar in immature anatids, no facet in the pterygoid is yet developed. As in immature specimens, no distinct hemipterygoid is found in any surveyed adult anatids.

### GALLIFORMES

We directly examined seven immature and five adult galliform specimens (see suppl. table 1) representing Megapodiidae, Numididae and Phasianidae (figs. 11-13). A larger sample of adult Galliformes, including Cracidae, was also consulted for comparative purposes (suppl. table 2). We found substantial differences in the morphology and the degree of ontogenetic change between megapode and non-megapodiid galliforms, which are therefore described separately below.

**Figure 11:**
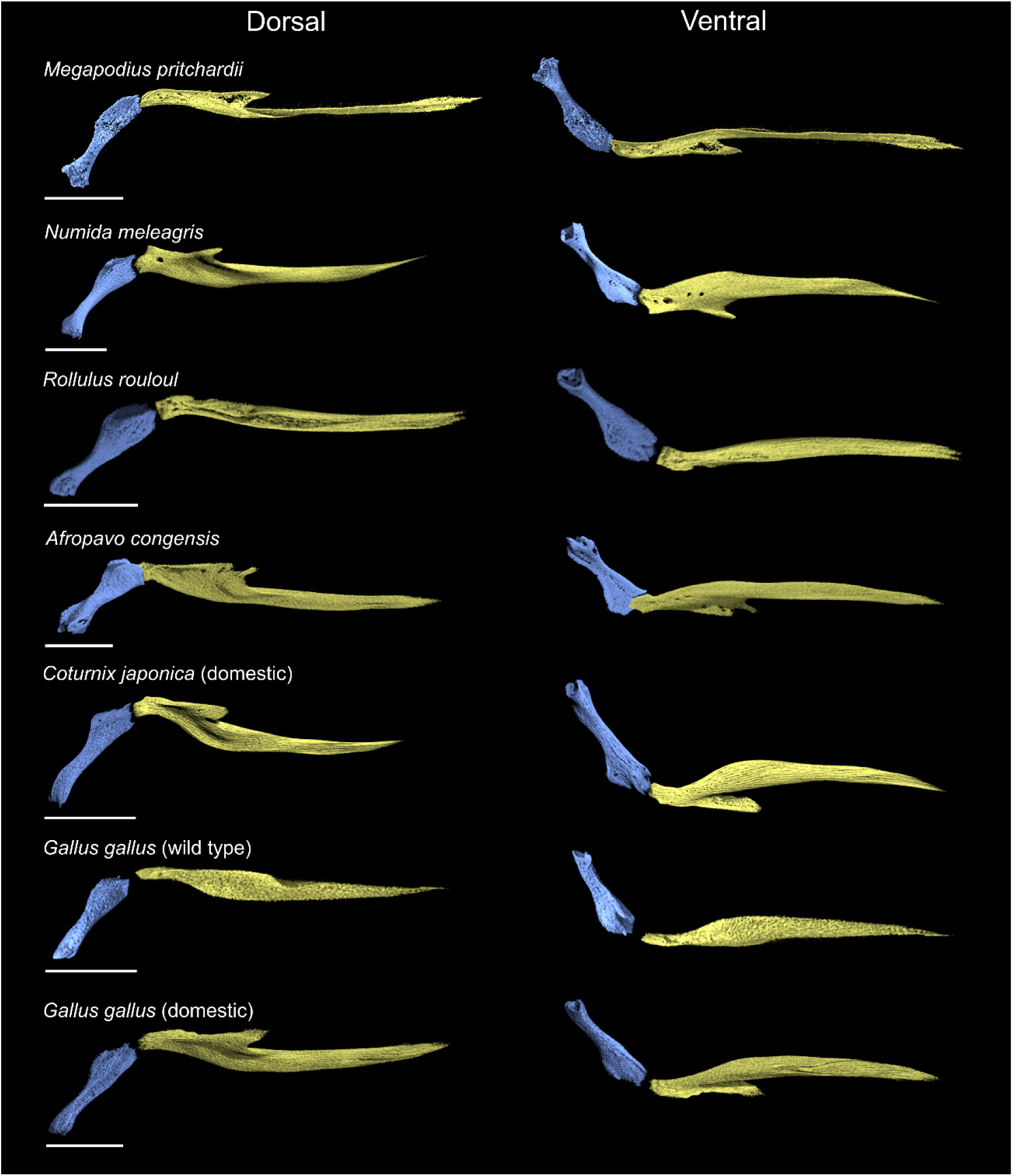
Comparative morphology of the palatine (yellow) and pterygoid (blue) in articulation in select immature galliforms, figured for the right side of the skull (*Megapodius*, *Numida*, *Rollulus*, *Coturnix* and *Gallus* (domestic) elements are reflected). ‘Dorsal’ and ‘ventral’ orientations are with respect to the skull. All scale bars represent 2.5 mm.

**Figure 12:**
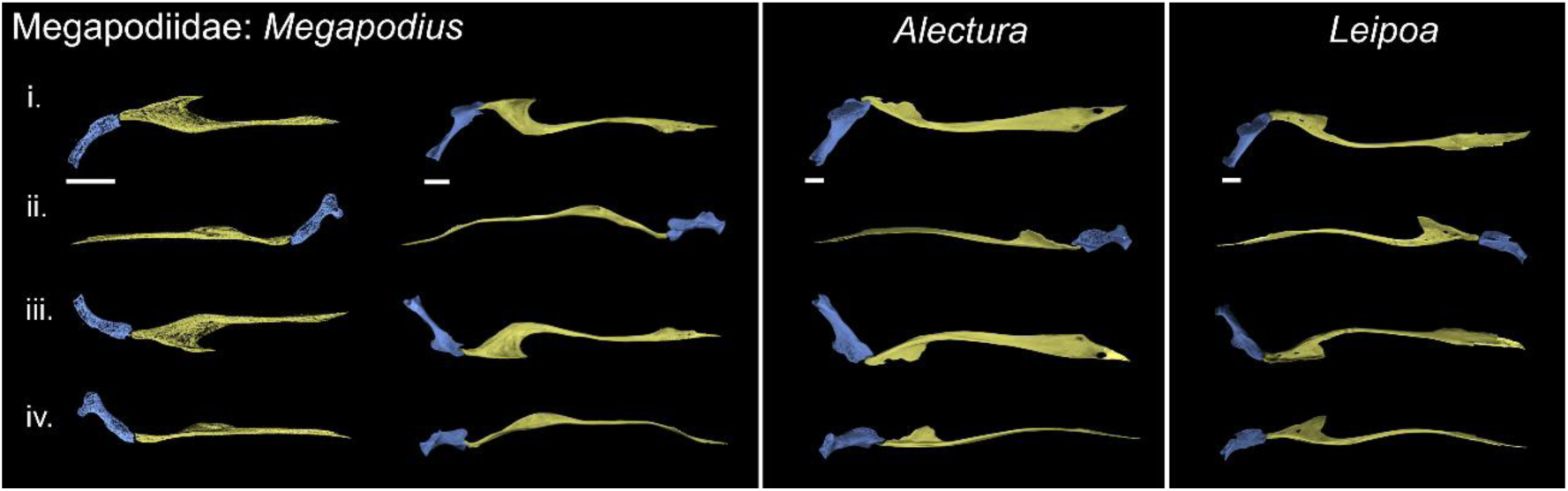
Comparison of Megapodiidae palatine (yellow) and pterygoid (blue) morphology, figured for the right side of the skull (*Megapodius* immature and *Alectura* elements are reflected). Left box, *Megapodius pritchardii* (left column – immature); *Megapodius nicobariensis* (right column – mature), middle box mature *Alectura lathami*, right box mature *Leipoa ocellata*. i.- iv. represent the following orientations: i.- dorsal, ii.- medial, iii.- ventral and iv.- lateral (oriented with respect to the skull itself). All orientations of the immature of *Megapodius* are slightly oblique to enhance comparability of morphology between the immature and mature. All scale bars represent 2.5 mm.

**Figure 13:**
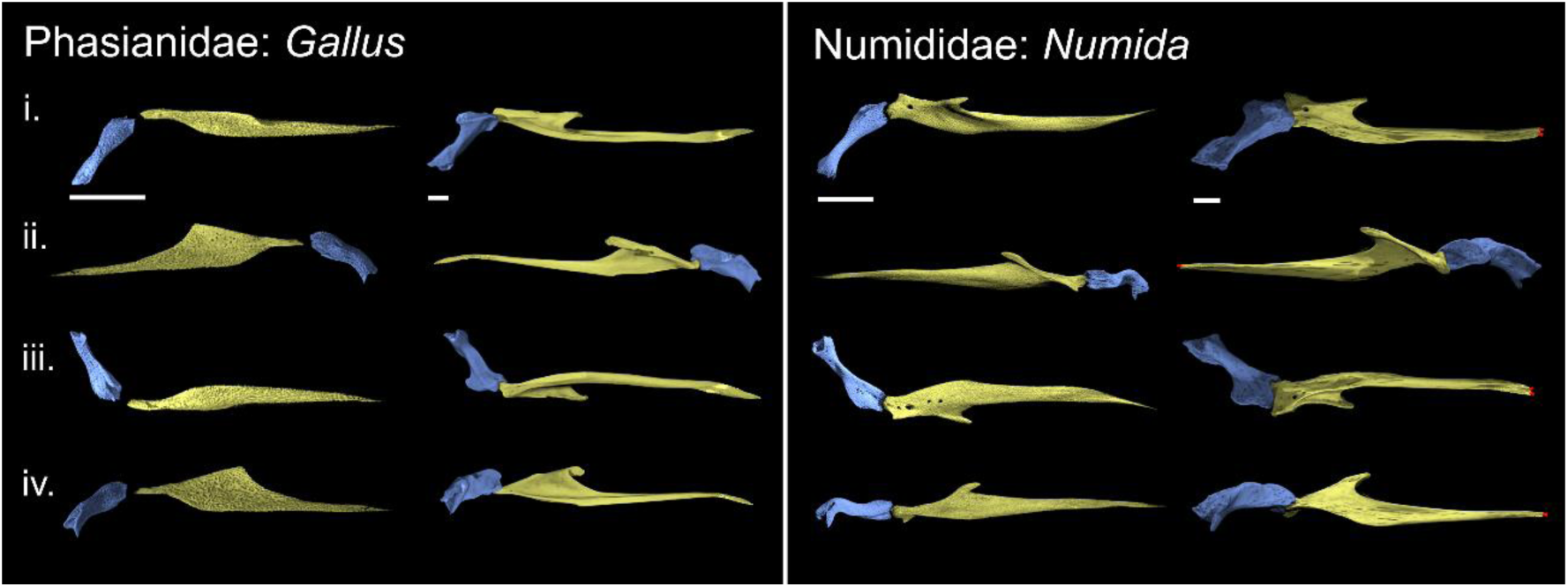
Comparison of non-megapodiid galliform palate morphology through ontogeny. immature (left column) and mature (right column) comparison of the palatine (yellow) and pterygoid (blue), figured for the right side of the skull (*Numida* immature elements are reflected). Red dotted lines represent areas where bone margins cannot be precisely delineated due to fusion. Phasianidae pairing: immature *Gallus gallus* (Red Junglefowl, wild type), mature *Gallus gallus* (Red Junglefowl, domestic); Numididae pairing: immature and mature *Numida meleagris* (Helmeted Guineafowl). i.- iv. represent the following orientations: i.- dorsal, ii.- medial, iii.- ventral and iv.- lateral (all with respect to the skull). All scale bars represent 2.5 mm.

#### MEGAPODIIDAE

##### Immature

The caudal region of the palatine of the hatchling-stage *Megapodius* extends into a rounded, dorsoventrally flattened and elongate pterygoid process, which is poorly ossified. The pterygoid process is broadly aligned with the main axis of the palatine in dorsal view, but with a slight lateral deflection at its caudal end (figs. 11, s6).

The rostralmost end of the pterygoid is incompletely ossified, and as such little relevant osteology is discernible. However, a shallow, poorly defined socket sits at the dorsolateral surface of the rostral end of the pterygoid. The pterygoid and palatine are not in contact, likely as a consequence of the incomplete ossification of the articular regions of both elements. An incipient basipterygoid process articular facet is present at the rostralmost end of the pterygoid, immediately adjacent to the shallow socket mentioned above (fig. s7).

##### Mature

The palatine appears to undergo a notable degree of rotation during ontogeny in *Megapodius*, such that the rostromedial process is more dorsally positioned in the immature compared to the adult (figs. 11, 12). The limited ossification of the sampled hatchling *Megapodius* precludes detailed morphological comparisons with the adult, but overall, the palatine and pterygoid of *Megapodius* do not appear to show substantial changes in morphology across ontogeny, apart from becoming more complex in structure.

The palatine of adult *Megapodius* bears a mediolaterally narrow and rostrocaudally elongate pterygoid process, which extends into a short mediolaterally compressed and laterally deflected projection at its tip. This projection, whose dorsomedial surface is slightly concave, appears to function as a condyle, inserting deeply into the articular cotyle of the pterygoid, and is absent or unossified in the hatchling specimen. Altogether, the pterygoid process of the mature *Megapodius* specimen is comparatively gracile relative to the immature specimen (fig. 12). A similarly elongate and dorsoventrally flattened pterygoid process is present in *Alectura lathami,* although in *Alectura* the pterygoid process terminates caudally at a rounded, dorsally deflected strap-like condyle, rather than a pointed tip. In *Leipoa ocellata* the pterygoid process is also relatively elongate, yet it terminates caudally into a mediolaterally wide rectangular facet, exhibiting a shallow notch at its centre (fig. 12).

In contrast to the incompletely ossified condition of the hatchling, the rostral region of the pterygoid of the adult *Megapodius* bears a short but robust rostral projection. The rostral surface of this structure encloses a deeply concave cup-like cotyle for the palatine, which is open dorsally and ventrally but bounded by well-developed margins on its medial and lateral sides. This cotyle-bearing process is strongly reminiscent of that present in Anhimidae (fig. 9). The pterygoids of both *Alectura* and *Leipoa* bear similar short rostral projections, yet these do not bear clear cotyles for the palatine (fig. 12). In *Alectura* this rostral projection dorsolaterally overlies a portion of the pterygoid process of the palatine, and the caudal condyle of the pterygoid process articulates with a shallow concavity just ventral to the pterygoid’s rostral projection. In *Leipoa,* the rostral end of the rostral projection forms a round condyle-like surface which abuts the rectangular and slightly notched caudal end of the pterygoid process of the palatine, lacking any overlap or deep insertion as in *Megapodius* and *Alectura*.

We do not observe a distinct hemipterygoid or significant changes in the shape of the palatine and pterygoid throughout ontogeny that would be consistent with pterygoid segmentation in Megapodiidae.

#### NON-MEGAPODIID GALLIFORMS

##### Immature

The surveyed immature non-megapode galliforms vary substantially in developmental stage and degree of ossification. These range from poorly ossified specimens at very early ontogenetic stages (e.g., near-hatching embryos) where the palatine and the pterygoid do not contact and do not show distinct morphologies (*Gallus gallus*, domestic and wild type), early post-hatching immatures where the palatine and the pterygoid are more ossified and complex, yet still do not come into contact (*Numida, Coturnix*, *Rollulus, Ortalis*), and more mature juveniles where both elements articulate with each other (*Afropavo*). All surveyed non-megapode galliforms appear to show comparable patterns of palate development.

The caudal portion of the palatine of the examined immature non-megapodiid galliforms is comparatively simple, extending into a rostrocaudally elongated and mediolaterally narrow sub-cylindrical or sub-rectangular pterygoid process (fig. s6). These are dorsoventrally compressed, simple, and mostly straight in both domestic and wild type *Gallus*, but the more mature representatives of Phasianoidea examined exhibit distinct laterally deflected condyles at their caudal ends. In *Numida, Coturnix* and *Rollulus*, these articular condyles are bulbous and robust, mediolaterally wide and terminate into a squared and flat caudal articular surface; in *Coturnix* and *Rollulus* a shallow notch on the dorsomedial surface of the pterygoid process separates two distinct dorsolateral and ventromedial sub-condyles. By contrast, the pterygoid process of *Afropavo* extends into a bifurcated articular surface, with a mediolaterally compressed and rostrocaudally elongated ventrolateral lip and a shorter medially directed dorsomedial lip, separated by a shallow concavity which wraps around the rostral surface of the pterygoid. Additionally, a short, caudomedially directed process extends from the medial surface of the pterygoid process in both *Numida* and *Coturnix*, terminating in a point (fig. s6).

Despite varying degrees of development, the rostral end of the pterygoid is incompletely ossified and indistinct in all surveyed immature non-megapodiid galliforms, forming a wide cup-like concavity that does not contact the palatine, with the exception of *Afropavo* (fig. 11). In *Afropavo,* the rostral surface of the pterygoid forms a wide saddle-shaped articular surface that is contacted by the pterygoid. The articular surface for the basipterygoid process is only incipiently developed in some of the immature galliforms examined (most notably in *Numida*), and is situated immediately caudal to the rostral end of the pterygoid, adjacent to the palatine articular socket (fig. s7).

##### Mature

The caudal region of the palatine of mature *Gallus* and *Numida* closely mirrors the condition in the immature specimens, illustrating little change in morphology through post-hatching ontogeny (fig. 13). Both mature *Gallus* and *Ortalis* exhibit a comparatively simple, dorsoventrally compressed and rostrocaudally elongated pterygoid process. Contrastingly, that of mature *Numida* is relatively short, robust and squared, and, unlike the laterally deflected process of immature *Numida*, is broadly aligned with the main axis of the palatine. The caudal end of the pterygoid process of *Gallus* bears a laterally offset and dorsally projected articular condyle, which forms a distinct rounded protrusion extending from the main body of the palatine (fig. 13) (this rounded protrusion is absent in *Ortalis*). In mature *Numida,* the caudal condyle is strap-like and mediolaterally narrow, without a pronounced dorsal extension. The caudomedially oriented pointed process present in the palatine of the immature *Numida* develops into a large and sharp process contacting a portion of the rostralmost margin of the basipterygoid process facet of the pterygoid. The mature *Gallus* develops a very small, rounded process in this region which is comparable in relative size to that of immature *Coturnix*.

The rostral end of the pterygoid of non-megapode galliforms does not undergo substantial morphological change across ontogeny, although the rostral articular surface becomes more completely ossified. In contrast to the condition in megapodes, no non-megapode galliforms examined exhibit or develop any sort of rostral pterygoid projection, and the rostral surface of the pterygoid is dominated by a conspicuously deep, cup-shaped cotyle for the palatine. In *Numida,* this cotyle is unbounded ventrally, but is delimited by pronounced margins on all other sides, and envelops much of the palatine condyle. In *Gallus*, the cotyle is open both medially and ventrally (fig. 13). In all surveyed adult non-megapode galliforms, the cotyle is situated approximately level with the rostral end of the basipterygoid process facet.

As in Megapodiidae, we find no evidence of a distinct hemipterygoid or significant changes in the shape of the palatine and pterygoid through ontogeny consistent with pterygoid segmentation in non-megapodiid galliforms.

#### NEOAVES

We directly examined 22 immature and 14 adult neoavian specimens (see suppl. table 1; figs. 14-20). This sample includes immature-adult pairs representing most major neoavian subclades: Strisores (Apodidae), Otidimorphae (Musophagidae), Columbimorphae (Columbidae), Gruiformes (Rallidae), Mirandornithes (Phoenicopteridae), Charadriiformes (Recurvirostridae), Aequornithes (Spheniscidae, Procellariidae, Sulidae, and Ardeidae), Opisthocomidae, and Telluraves (Cathartidae, Ramphastidae, and Psittacidae) (figs. 17-20). As our sample incorporates immature specimens at varying ontogenetic stages (ranging from late-stage embryos to older immature individuals) and mature representatives, we were able to observe a range of discrete points in the process of pterygoid segmentation. We observed evidence for the presence of a hemipterygoid in all surveyed immature representatives of Neoaves, which was variably connected to either the pterygoid or the palatine (or present as an entirely discrete element), while this element was entirely confluent with the palatine in all sampled adults, becoming part of the pterygoid process of the palatine. Hence we infer that the rostral projection of the pterygoid of neoavians is homologous with the hemipterygoid element (e.g., see Pycraft (1901)). To facilitate the characterization of this process, we grouped the specimens below into logical, hypothetical stages based on the observed relationships between the hemipterygoid and the other main palatal bones with reference to Pycraft (1901). Stage 1 corresponds to the hemipterygoid and pterygoid forming a single structure – thus there is no distinct, identifiable, hemipterygoid (i.e. ‘unsegmented hemipterygoid’); Stage 2 corresponds to a distinct hemipterygoid that is attached to the pterygoid (i.e. ‘partially segmented hemipterygoid’); Stage 3 corresponds to a discrete hemipterygoid that is fully independent and therefore detached from the pterygoid without having begun fusion to the palatine (i.e. ‘fully segmented hemipterygoid’); Stage 4 corresponds to a distinct hemipterygoid partially fused to the palatine (associated with a ‘fully segmented hemipterygoid’); and Stage 5 to the absence of a recognisable hemipterygoid distinct from the palatine (i.e. complete fusion of the hemipterygoid to the caudal palatine) (figs. 21, 22). Despite some degree of variation in the morphology of the different palatal elements, the entire process of pterygoid segmentation and major associated ontogenetic changes are present across all sampled representatives of Neoaves, regardless of subclade, and they are thus described together. Considering the general similarities among all sampled adult neoavians in the arrangement of their palatine-pterygoid contacts, and the pronounced variation among sampled immatures in the observed stage of segmentation progression, we begin the neoavian descriptions below by establishing the condition in mature individuals to better contextualize our observations of stage-by-stage post-hatching ontogenetic change, which follow.

##### Adult condition (figs. 21, 22, STG. 5)

All examined mature representatives of Neoaves exhibit a similar arrangement of the palatine-pterygoid contact (apart from the highly divergent morphologies of parrots (fig. 20; suppl. table 2), with a relatively simple condylar articulation between both elements, and only moderate variation across our taxon sample (figs. 17-20). Generally, the caudal end of the palatine extends into a rostrocaudally elongate pterygoid process that terminates in a dorsoventrally deep and mediolaterally narrow caudal articular surface for the pterygoid in most surveyed neoavians. The dorsal and ventral margins of the pterygoid process are continuous with the respective dorsal and ventral crests of the palatine (*margo dorsalis palatini* and *margo ventralis palatini* see Zusi and Livezey (2006)). Although the caudal end of the pterygoid process appears to form a single continuous vertical hinge-like articular surface in certain taxa (e.g., *Tauraco*) (fig. 17), in some other cases the dorsal and ventral margins of the articular surface form two distinct dorsal and ventral mediolaterally narrow strap-like condyles separated by a variably developed incisure (e.g., *Gallirallus, Recurvirostra*) (figs. 17, 18, respectively). Certain taxa exhibit a less complex pterygoid process terminating caudally into a distinct, rounded single condyle, as exemplified by *Apus* and *Opisthocomus* (figs. 17, 19, respectively). Only in parrots (e.g., *Ara*) does the pterygoid process, like the rest of the palatine, differ markedly from that of other neoavians (fig. 20). In parrots, the pterygoid process is rostrocaudally shorter than in other neoavians, and positioned further rostrally on the palatine, rather than at its caudal end. The articulation surface for the pterygoid is squared and flat or slightly concave, rather than forming a condyle.

The pterygoid of most surveyed neoavians is comparatively simple, generally forming an elongated rod-like structure that is often mediolaterally compressed, with marked medial and lateral depressions and pronounced crests or ridges whose orientation can vary across taxa. Notably, both *Spheniscus* and *Opisthocomus* exhibit a broad, dorsoventrally compressed flange or wing that extends laterally from the main body of the pterygoid. The rostral end of the pterygoid encompasses the articulation for the palatine on its medial margin, as well as a broad subtriangular articular surface for the parasphenoid rostrum (see below). The articulation for the palatine is generally dorsoventrally deep and mediolaterally narrow, and can range from a featureless hinge-like ridge oriented dorsoventrally, as in *Tauraco* (fig. 17) and *Columba* (fig. 17) to a more complex structure with distinct dorsoventrally arranged shallow cotylae or facets which articulate with the dorsal and ventral condyles of the pterygoid process of the palatine, as in *Gallirallus* and *Vultur* (figs. 17, 19, respectively). In certain taxa, either the dorsal or the ventral portions of the articular surface for the palatine develop into a large cup-like cotyle, matching a rounded condyle of the palatine, while the opposite end of the palatine articular surface remains simple and hinge-like. In *Nycticorax* the dorsal articular surface is cup-like and the ventral joint hinge-like (fig. 19), while *Pteroglossus* exhibits the opposite condition, with a cup-like ventral surface and a hinge-like dorsal joint (fig. 20). An additional variably developed articular facet between the palatine and the pterygoid is present on the rostral end of the pterygoid in some surveyed neoavians, situated lateral to the main palatine-pterygoid hinge. This structure, when present, ranges from a small subtriangular flat or slightly concave surface which contacts a mediolateral broadening of the pterygoid process of the palatine, as in *Recurvirostra* (fig. 18), to a more complex interlocking mechanism. The latter encompasses a deep cup-like cotyle which bears a lateral hook-shaped projection wrapping around the lateral side of the palatine as in *Gallirallus, Phoenicopterus, Opisthocomus* and *Vultur* (figs. 17-19, respectively), though the specific nature of this additional contact appears to be highly clade-specific.

In contrast to the condition amongst Galloanserae, only a few mature neoavians exhibit an articular facet for the basipterygoid process, including *Columba*, *Vultur* and *Recurvirostra*. In all sampled neoavians in which the basipterygoid process articulates with the pterygoid, this facet is generally situated further caudally than in galloanserans and is not adjacent to or involved in the palatine-pterygoid articulation. However, it is worth noting that an additional contact between the palatal bones and the parasphenoid rostrum is present in the examined mature Neoaves. This contact, variable in rostrocaudal length but always overlapping with the palatine-pterygoid articulation, involves both the palatine and the pterygoid, with both elements wrapping around the parasphenoid rostrum along a variably defined flat surface on the medial side of both bones.

##### Immature condition

###### STAGE 1 - UNSEGMENTED HEMIPTERYGOID (figs. 21, 22, STG. 1)

Among the examined immature specimens of **Musophagiformes** (*Tauraco* and *Musophaga*), **Columbiformes** (*Otidiphaps*), **Piciformes** (*Pteroglossus*), and **Psittaciformes** (*Ara* and *Lorius*) (figs. 14, 16), we observed an elongate rostral projection extending from the rostralmost region of the pterygoid that is not present in any of the adults examined (figs. 17, 20). We did not observe any sutures or bony separations distinguishing this rostral projection from the rest of the pterygoid. Therefore, we hypothesise that these immature specimens represent **Stage 1** of pterygoid segmentation, characterized by an **unsegmented hemipterygoid** (figs. 21, 22, STG.1). The caudal region of the palatine extends into a caudally projected process which is caudally tapered and dorsoventrally narrow (figs. s8, s9). This process exhibits a simple flattened dorsomedial surface over which the rostral projection of the pterygoid, or ‘unsegmented hemipterygoid’, extends. This flattened morphology strongly contrasts with that of the more complex pterygoid process of the palatine of the relevant adults, which exhibit marked dorsal and ventral extensions of the palatine margins (i.e. *margo dorsalis palatini* and *margo ventralis palatini*, see Zusi and Livezey (2006)) and a dorsoventrally deep and mediolaterally narrow palatine-pterygoid contact, sometimes divided into two distinct condyles (see above). This major change in morphology is best exemplified by the immature-adult pairings of *Tauraco* and *Pteroglossus* (figs. 17, 20). Importantly, as with adults, the immature parrots examined exhibit highly divergent morphologies of both the palatine and pterygoid compared to other immature neoavians.

**Figure 14:**
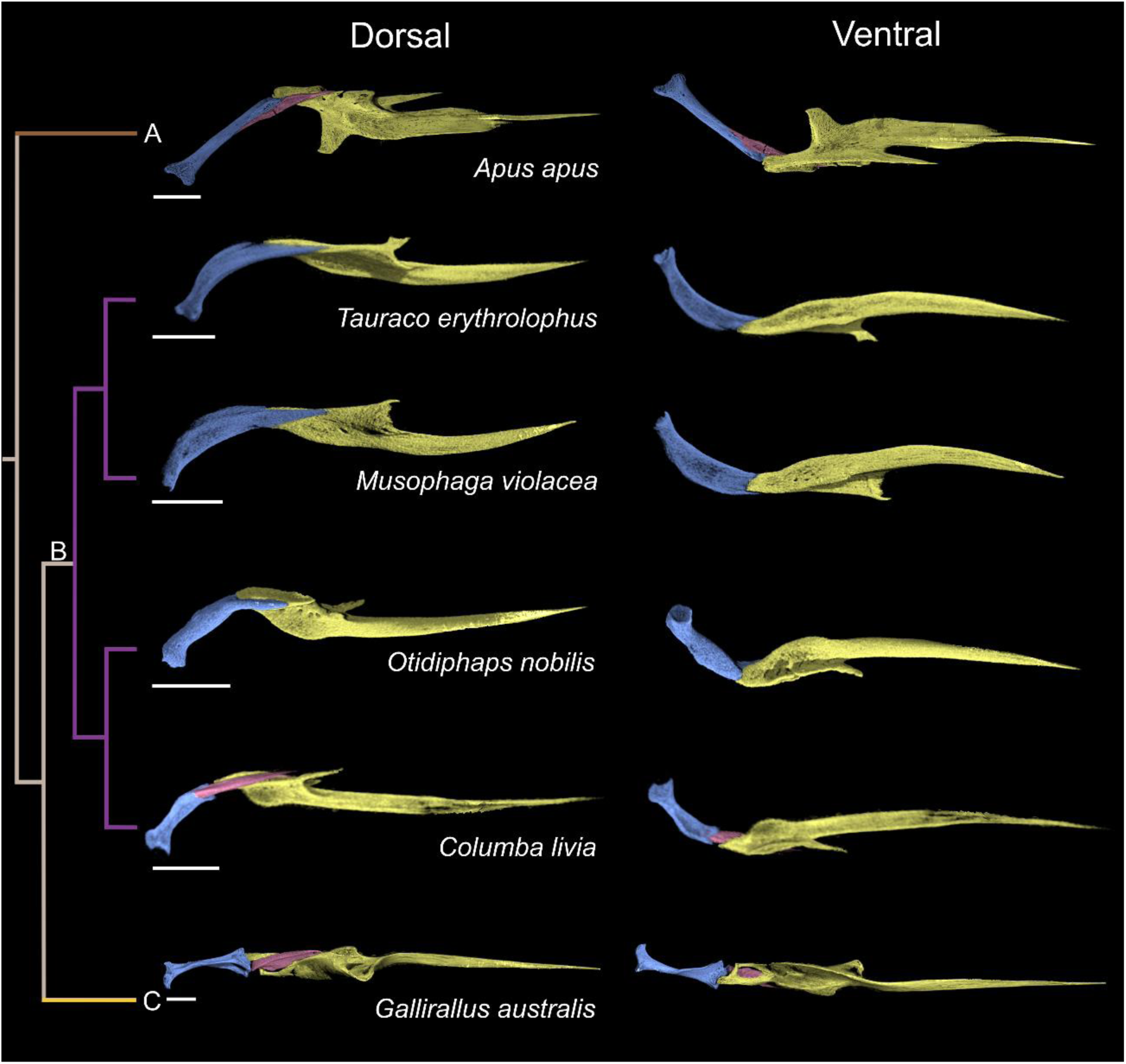
Comparative morphology of the palatine (yellow), pterygoid (blue) and hemipterygoid (pink) (partially/fully segmented) in select immature neoavians, figured for the right side of the skull (*Apus* and *Otidiphaps* elements are reflected). ‘Dorsal’ and ‘ventral’ orientations are with respect to the skull. Left, cladogram illustrating the phylogenetic relationships of these taxa (higher order relationships follow Prum et al. (2015); detailed interrelationships among subclades follow Cooney et al. (2017)). **A** – Strisores; **B** – Columbaves; **C** – Gruiformes. All scale bars represent 2.5 mm.

**Figure 15:**
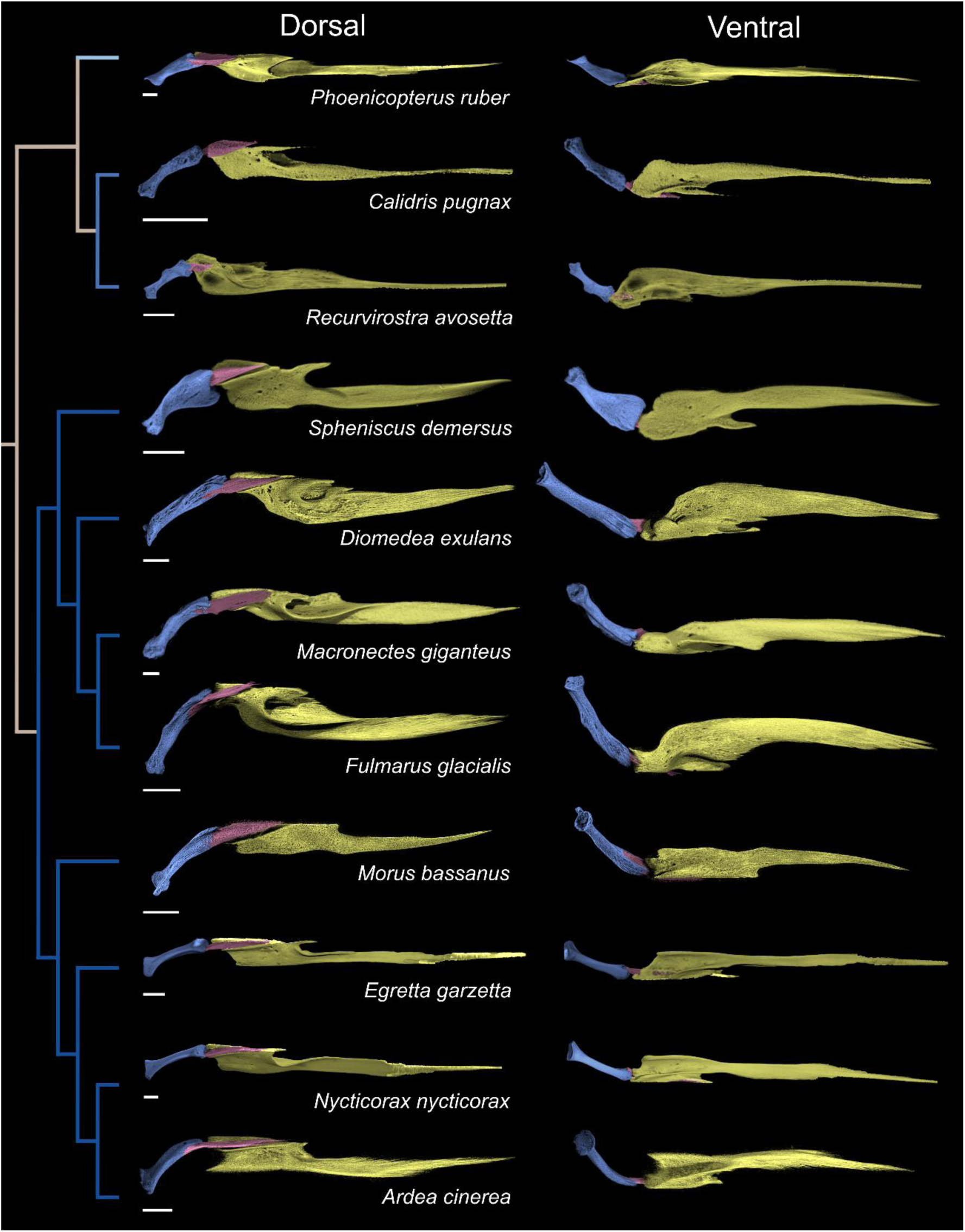
Comparative morphology of the palatine (yellow), pterygoid (blue) and hemipterygoid (pink) (partially/fully segmented) in select immature members of the neoavian subclade Aequorlitornithes (Prum et al., 2015), figured for the right side of the skull (*Phoenicopterus*, *Calidris*, *Spheniscus* and *Macronectes* elements are reflected). ‘Dorsal’ and ‘ventral’ orientations are with respect to the skull. Left, cladogram illustrating the phylogenetic relationships of these taxa (higher order relationships follow Prum et al. (2015); detailed interrelationships among subclades follow Cooney et al. (2017)). In *Egretta* and *Recurvirostra* the rostral regions of the hemipterygoid are partially fused to the palatine. All scale bars represent 2.5 mm.

**Figure 16:**
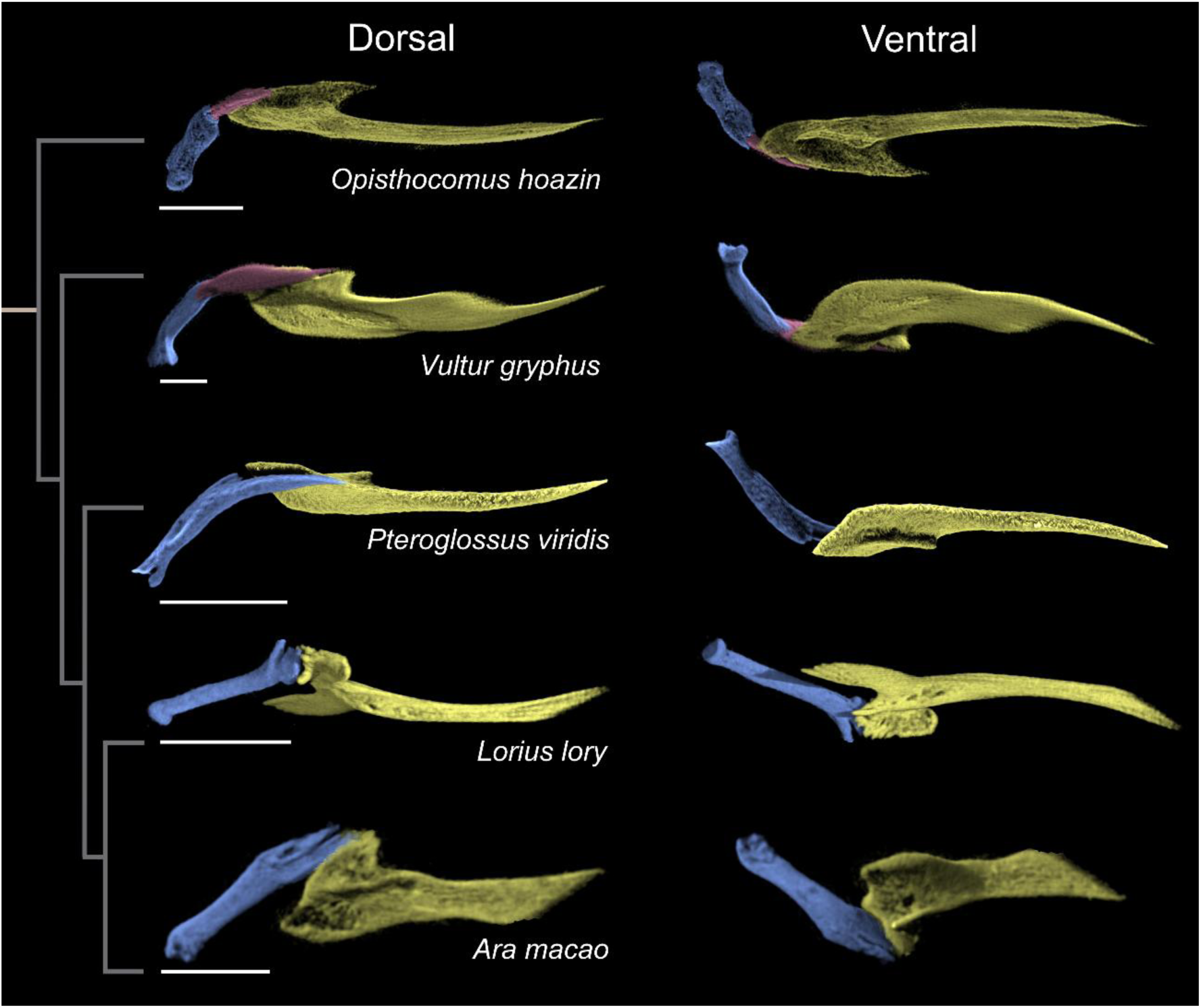
Comparative morphology of the palatine (yellow), pterygoid (blue) and hemipterygoid (pink) (partially/fully segmented) in select immature members of the neoavian subclade Inopinaves (Prum et al., 2015), figured for the right side of the skull (*Opisthocomus*, *Pteroglossus* and *Lorius* elements are reflected). ‘Dorsal’ and ‘ventral’ orientations are with respect to the skull. Left, cladogram illustrating the phylogenetic relationships of these taxa (higher order relationships follow Prum et al. (2015); detailed interrelationships among subclades follow Cooney et al. (2017)). All scale bars represent 2.5 mm.

**Figure 17:**
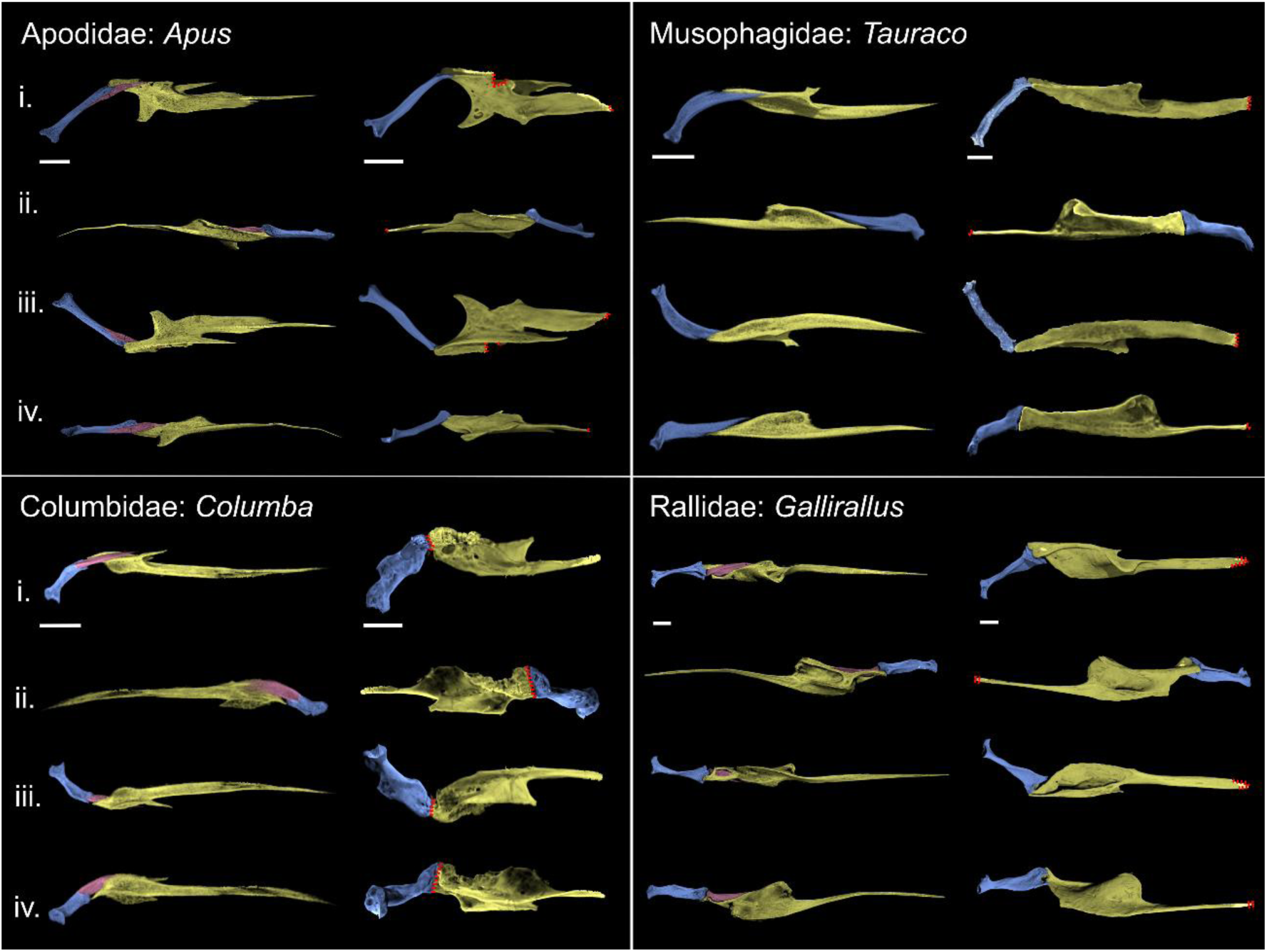
Comparison of immature (left columns of each ‘box’) and mature (right columns of each ‘box’) neoavian palatine (yellow), pterygoid (blue) and hemipterygoid (pink) (partially/fully segmented) (part 1), figured for the right side of the skull (immature *Apus* and mature *Columba* elements are reflected). Red dotted lines represent areas where bone margins cannot be precisely delineated due to fusion. Apodidae pairing: *Apus apus* (Common Swift) immature and mature; Musophagidae pairing: *Tauraco erythrolophus* (Red-Crested Turaco) immature and mature; Columbidae pairing: *Columba livia* (Rock Dove) immature and *Columba livia domestica* (Domestic pigeon) mature; Rallidae pairing: *Gallirallus australis* (Weka) immature and mature. i - iv. represent the following orientations: i.- dorsal, ii.- medial, iii.- ventral and iv.- lateral (all with respect to the skull). All scale bars represent 2.5 mm.

**Figure 18:**
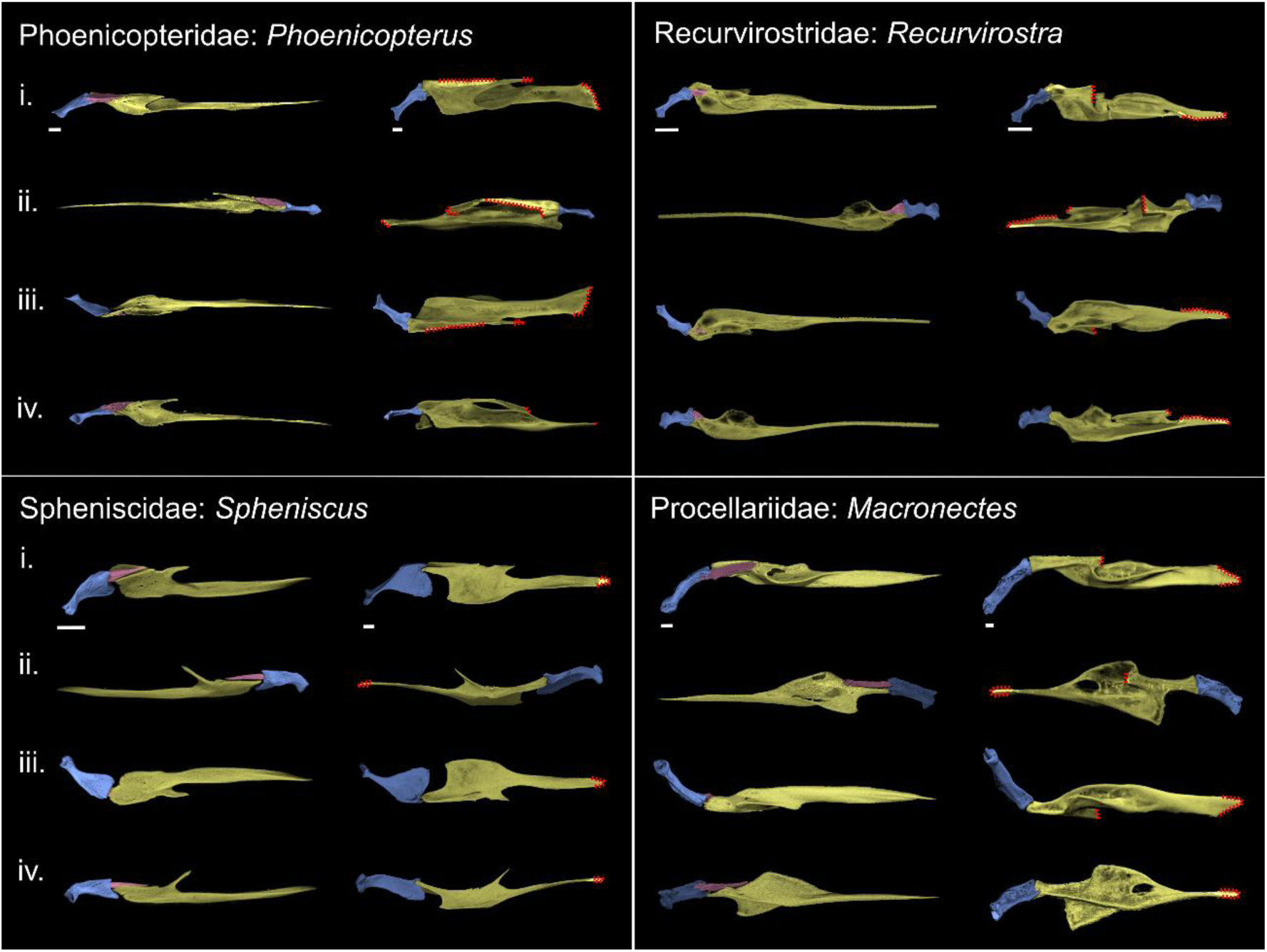
Comparison of immature (left columns of each ‘box’) and mature (right columns of each ‘box’) neoavian palatine (yellow), pterygoid (blue) and hemipterygoid (pink) (partially/fully segmented) (part 2), figured for the right side of the skull (immature and mature *Phoenicopterus*, immature *Spheniscus* and immature and mature *Macronectes* elements are reflected). Red dotted lines represent areas where bone margins cannot be precisely delineated due to fusion. In immature *Recurvirostra* the rostral region of the hemipterygoid is partially fused to the palatine. Phoenicopteridae pairing: *Phoenicopterus ruber* (American Flamingo) immature and mature; Recurvirostridae pairing *Recurvirostra avosetta* (Pied Avocet) immature and mature; Spheniscidae pairing: *Spheniscus demersus* (African Penguin) immature and mature, Procellariidae pairing: *Macronectes giganteus* (Southern Giant Petrel) immature and *Macronectes halli* (Northern Giant Petrel) mature. i - iv. represent the following orientations: i.-dorsal, ii.- medial, iii.- ventral and iv.- lateral (all with respect to the skull). All scale bars represent 2.5 mm.

**Figure 19:**
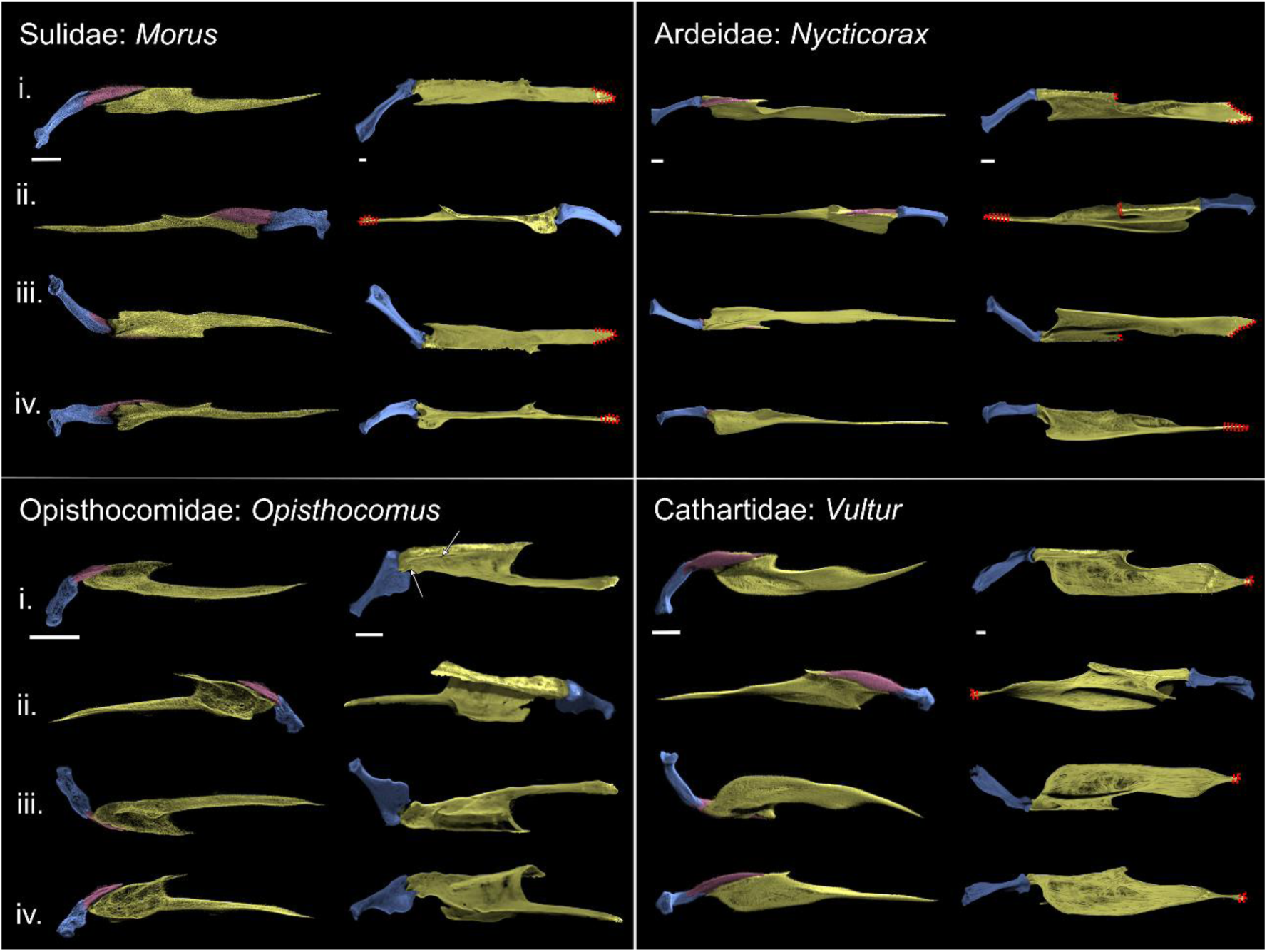
Comparison of immature (left columns of each ‘box’) and mature (right columns of each ‘box’) neoavian palatine (yellow), pterygoid (blue) and hemipterygoid (pink) (partially/fully segmented) (part 3), figured for the right side of the skull (immature *Opisthocomus* elements are reflected). Red dotted lines represent areas where bone margins cannot be precisely delineated due to fusion. White arrows denote an apparent remnant suture line separating the fused hemipterygoid and palatine in an adult skull of *Opisthocomus hoazin*. Sulidae pairing*: Morus bassanus* (Northern Gannet) immature and mature; Ardeidae pairing: *Nycticorax nycticorax* (Black-Crowned Night Heron) immature and mature; Opisthocomidae pairing: *Opisthocomus hoazin* (Hoatzin) immature and mature; Cathartidae pairing: *Vultur gryphus* (Andean Condor) immature and mature. i - iv. represent the following orientations: i.- dorsal, ii.- medial, iii.- ventral and iv.- lateral (all with respect to the skull). All scale bars represent 2.5 mm.

**Figure 20:**
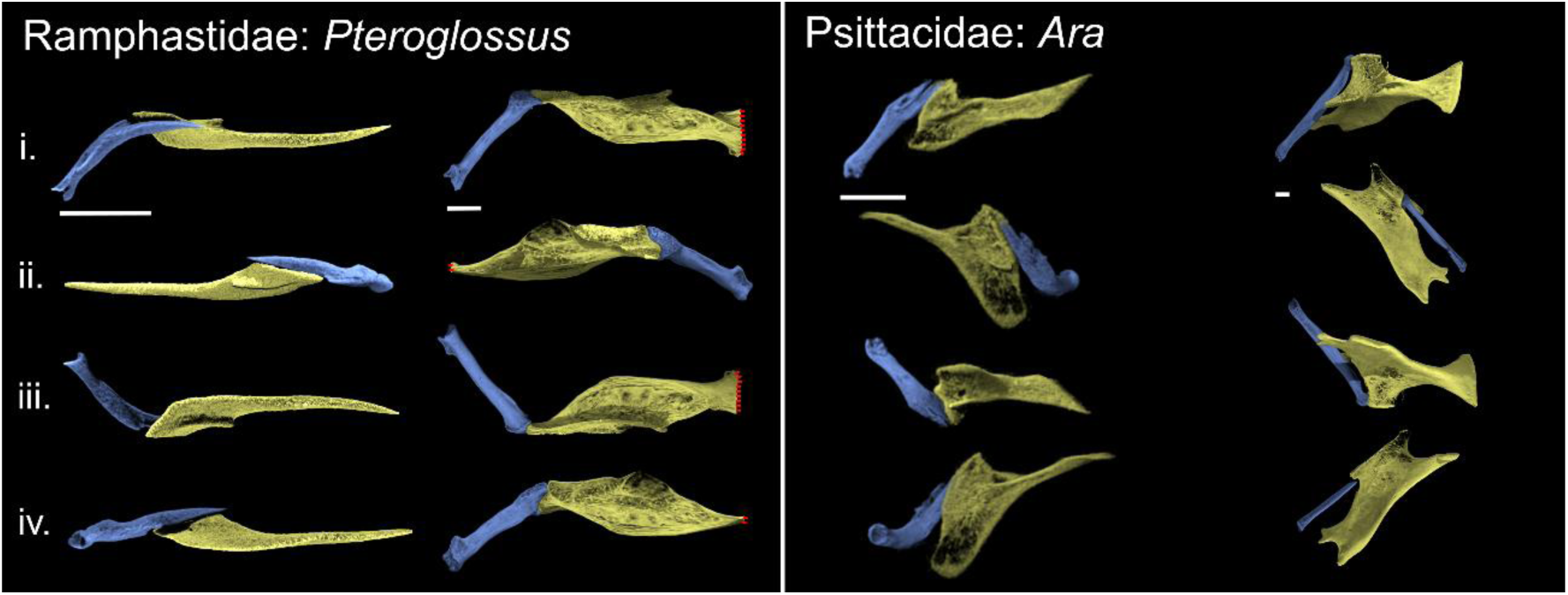
Comparison of immature (left columns of each ‘box’) and mature (right columns of each ‘box’) neoavian palatine (yellow) and pterygoid (blue) (part 4), figured for the right side of the skull (immature *Pteroglossus* elements are reflected). Red dotted lines represent areas where bone margins cannot be precisely delineated due to fusion. Ramphastidae pairing: *Pteroglossus viridis* (Green Aracari) immature and mature; Psittacidae pairing: *Ara macao* (Scarlett Macaw) immature and *Ara ambiguus* (Great Green Macaw) mature. i - iv represent the following orientations: i.- dorsal, ii.- medial, iii.- ventral and iv.- lateral (all with respect to the skull itself). All scale bars represent 2.5 mm.

Pterygoids of the Stage 1 immatures are short, robust and cylindrical, with a long projected and tapered rostral projection (unsegmented hemipterygoid) which forms a broad curve relative to the main body of the pterygoid (figs. s10, s11). We observed a marked difference in shape compared to adults, both in the length of the pterygoid and in its morphology, with adult pterygoids being longer and more complex in shape, generally mediolaterally compressed, and exhibiting a marked crest along their length as well as narrow, dorsoventrally elongated contacts with the palatine. In all these taxa, the palatine-pterygoid contact in immatures resembles that of adult Anseres; that is, the rostral, blade-shaped unsegmented hemipterygoid extends across the dorsal surface of the pterygoid process of the palatine, while the dorsoventrally narrow, caudal end of the pterygoid process of the palatine contacts the pterygoid.

###### STAGE 2 - PARTIALLY SEGMENTED HEMIPTERYGOID (figs. 21, 22, STG. 2)

**Figure 21:**
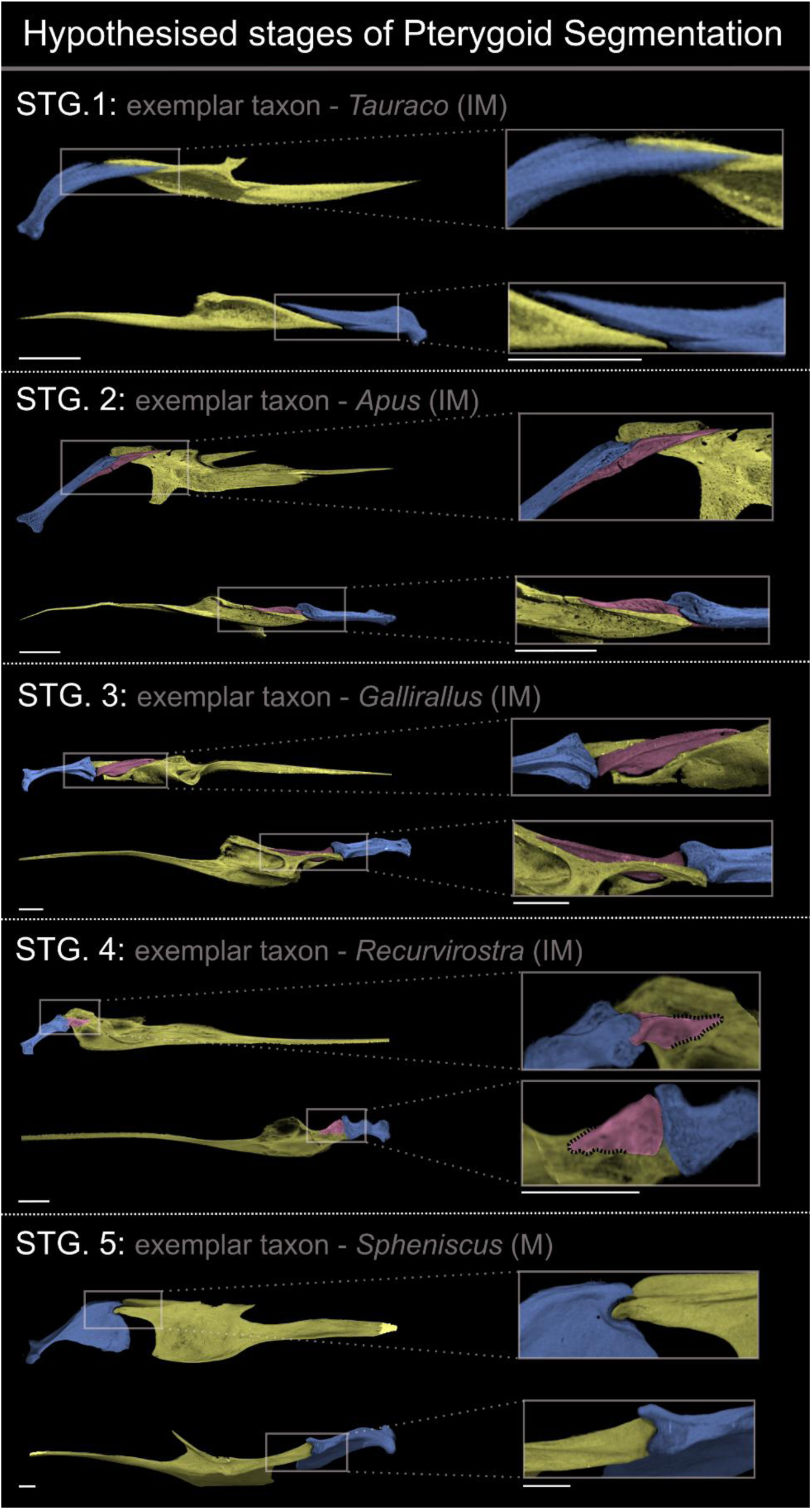
The process of pterygoid segmentation discretized into five hypothetical stages, illustrated by various immature (and one mature) neoavian specimens. Inset images show details of the relationship between the palatine (yellow), pterygoid (blue) and hemipterygoid (pink) (partially/fully segmented), figured for the right side of the skull (*Apus* elements are reflected). Upper images for each stage are in dorsal view and lower images for each stage are in medial view (oriented with respect to the skull). Black striped lines at Stage 4 denote partial fusion of the hemipterygoid with the palatine. Abbreviation: IM, immature; M, mature; STG, stage. All scale bars represent 2.5 mm.

**Figure 22:**
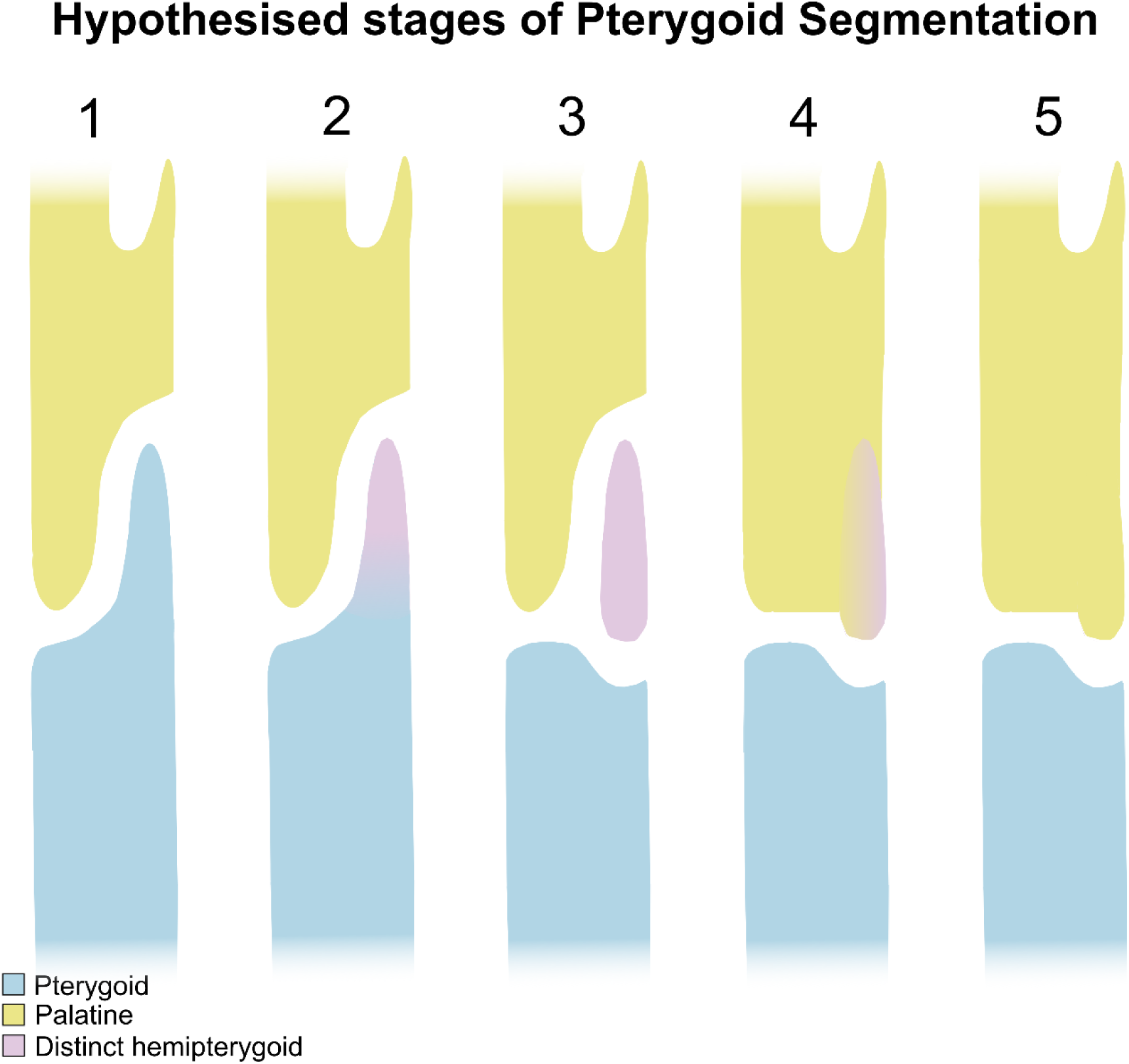
Schematic illustration of the process of pterygoid segmentation, discretized into five hypothetical stages. **1** – **Unsegmented hemipterygoid**: pterygoid bears a rostral projection homologous with the hemipterygoid. The rostral projection remains indistinguishable from the main body of the pterygoid; **2** – **Partially segmented hemipterygoid**: pterygoid bears a rostral projection that has begun to segment from the pterygoid body to form a distinct hemipterygoid; **3** – **Fully segmented hemipterygoid**: the distinct hemipterygoid has detached from the pterygoid, now representing a ‘discrete’ and therefore entirely separate element; **4** – **Partial fusion of hemipterygoid to caudal palatine**: discrete hemipterygoid has begun fusing to the palatine. **5** – **Complete fusion of the hemipterygoid to the caudal palatine:** no visible contact surface between the hemipterygoid and palatine. This stage is associated with the presence of an intrapterygoid articular surface.

Several sampled immature representatives of **Strisores** (*Apus*), **Columbiformes** (*Columba*), **Phoenicopteriformes** (*Phoenicopterus*), **Charadriiformes** (*Calidris*), **Sphenisciformes** (*Spheniscus*), **Suliformes** (*Morus*), **Procellariiformes** (*Macronectes, Diomedea, Fulmarus*), **Pelecaniformes** (*Nycticorax, Ardea*), **Opisthocomiformes** (*Opisthocomus*), and **Accipitriformes** (*Vultur*) (figs. 14-16), also exhibit an elongated blade-like rostral projection situated at the rostralmost region of the pterygoid, which is absent in all examined adults (figs. 17-19). However, in contrast with the Stage 1 specimens described above, the rostral projection (the hemipterygoid) is clearly distinguished from the pterygoid by a clear suture present along most of their contact (e.g., *Apus*, *Spheniscus*, *Nycticorax*), yet it remains attached or partially fused to the main body of the pterygoid (fig. 4). Therefore, we identify these immature specimens as being at **Stage 2**, characterized by a **partially segmented hemipterygoid** (figs. 21, 22, STG. 2).

All Stage 2 immature specimens observed exhibit a roughly subtriangular and rostrally tapered partially segmented hemipterygoid (figs. s10-s13). The length of the hemipterygoid is variable, which may be directly related to the rostrocaudal length of the caudalmost portion of the palatine. The position of the partially segmented hemipterygoid with respect to the main body of the pterygoid is variable across the surveyed members of Neoaves. In some species, the hemipterygoid lies primarily lateral to the main body of the pterygoid, forming a longitudinal overlapping contact (e.g., *Ardea, Diomedea*), often within a variably deep groove on the lateral surface of the pterygoid (e.g., *Fulmarus*). In other taxa, the hemipterygoid is set within a deep cavity or depression at the rostral end of the pterygoid, so that the main body of the pterygoid almost completely encloses the caudal end of the hemipterygoid (e.g., *Spheniscus* (figs. 4, 15)). Finally, in some taxa, the hemipterygoid shows a small flat contact with the rostral end of the pterygoid, such that little overlap between both elements exists (e.g., *Nycticorax*) (figs. 4, 15).

In contrast to Stage 1 immatures, we observe very little change in pterygoid morphology between Stage 2 immatures and adults (figs. 17-19). In Stage 2 immatures the pterygoid already exhibits an adult-like morphology: they are generally elongated, mediolaterally or dorsoventrally compressed, with various crests or ridges running rostrocaudally along the pterygoid body. The most notable exception amongst the sampled taxa is *Opisthocomus*, in which the Stage 2 pterygoid lacks a broad lateral flange present in the adult (fig. 19). Although less ossified, the morphology of the rostral surface of the pterygoid is generally already recognizable in Stage 2 immatures, despite the hemipterygoid still being attached to it. The ventral portion of the rostral pterygoid, when distinguishable, contacts the caudal end of the palatine in some taxa (e.g. *Apus*) but not in others (e.g., *Calidris*). The dorsal portion of the rostral articular surface of the pterygoid is attached to the hemipterygoid, yet nonetheless exhibits a similar morphology to the dorsal portion of the palatine-pterygoid articulation in adults. Similarly, in taxa where the partially segmented hemipterygoid is positioned within a deep cavity in the rostral pterygoid, the cavity resembles the round cotylar articulation (i.e., “ball-and-socket”) found in adults (e.g., *Spheniscus* (fig. 4)).

Unlike in Stage 1 immatures, the dorsal and ventral crests or extensions of the palatine margins seen in adults are already present in all surveyed Stage 2 immatures (except *Phoenicopterus* which lacks a well-developed ventral crest); both palatine crests are separated by a shallow medial depression. The hemipterygoid dorsomedially overlaps and fits within this depression or facet, with most of its lateral border lying along the dorsal crest of the palatine (figs. 17-19). When observed together, the combined morphology of the immature caudal palatine and the hemipterygoid strongly resemble the condition of the pterygoid process in adult Neoaves, with the hemipterygoid and dorsal palatine crest ultimately fusing to form the dorsal portion or condyle of the pterygoid process. Furthermore, examined adults lack the medial depression for the hemipterygoid between both palatine crests, this region is instead flat or concave, similar to the medial surface of the hemipterygoid. The ventral condyle (where present) is formed exclusively by the palatine. Notably, the sampled mature *Opisthocomus* specimen exhibits an incompletely fused suture across the dorsal portion of the pterygoid process (fig. 19), in a similar position to where the overlap between the hemipterygoid and the palatine are in the immature.

###### STAGE 3 - FULLY SEGMENTED HEMIPTERYGOID (figs. 21, 22, STG. 3)

Only one of the sampled immature specimens, the gruiform *Gallirallus australis*, exhibits a discrete, elongated hemipterygoid positioned between the palatine and the pterygoid, completely independent and detached from both (fig. 14).

Therefore, we identify this immature specimen as representing **Stage 3**, characterized by a **fully segmented hemipterygoid** (figs. 21, 22, STG. 3). The morphology of all elements in this specimen is extremely similar to the adult, such that no major subsequent ontogenetic change, beyond the fusion between the hemipterygoid and the palatine, occurs (fig. 17). Despite being a wholly detached element, the morphology of the hemipterygoid closely matches that of the dorsal portion of the pterygoid process of the palatine in adult *Gallirallus*, and its shape and the arrangement of the articulation between the hemipterygoid and the pterygoid is strongly reminiscent of that between the dorsal condyle of the palatine’s pterygoid process and the pterygoid in adults.

###### STAGE 4 – HEMIPTERYGOID PARTIALLY FUSED TO THE PALATINE (figs. 21, 22, STG. 4)

The sampled immature specimens of the charadriiform *Recurvirostra* and the pelecaniform aequornithine *Egretta* exhibit a distinct, elongated hemipterygoid positioned between the palatine and the pterygoid, yet attached to the dorsomedial surface of the palatine, with which it forms a diffuse but clear suture (figs. 4, 15). We identify these immature specimens as being at **Stage 4**, characterized by a **fully segmented hemipterygoid that is partially fused with the palatine** (figs. 21, 22, STG. 4). In *Egretta*, this observation is combined with tentative evidence for the incomplete detachment of the hemipterygoid from the pterygoid body, suggesting the possibility that some taxa may bypass Stage 3. However, scan resolution precludes a definitive assessment of whether this immature *Egretta* hemipterygoid remains attached to the pterygoid (fig. 4).

Similar to Stage 3, the hemipterygoid at Stage 4 remains a discrete element separated from the pterygoid, and the nature of their contact is essentially indistinguishable from that of the relevant adult specimens. However, the hemipterygoid appears to be in the process of fusing to the palatine along most of its length, making it difficult to distinguish clearly between both elements, particularly towards the rostral terminus of the hemipterygoid. A contact suture is more readily visible between the ventral surface of the caudalmost region of the hemipterygoid and the dorsal surface of the palatine in both immature specimens, suggesting that fusion of the hemipterygoid and the palatine progresses rostrocaudally.

## 4. DISCUSSION

In light of our examination of immature and mature individuals from across the extant diversity of birds, in conjunction with the early observations of Pycraft (1901)), we outline five hypothetical stages for the process of pterygoid segmentation (figs. 21, 22; table 1). Our data suggest that the complete process of post-hatching pterygoid segmentation (involving the formation of a distinct hemipterygoid, its detachment from the pterygoid body, and its fusion with the palatine), is exclusive to Neoaves among extant birds. We found no evidence of either a transient discrete hemipterygoid nor broad changes in the morphology and arrangement of the palatine and the pterygoid across ontogeny in Galloanserae or Palaeognathae consistent with the complete process of pterygoid segmentation. These observations are in agreement with some previous explorations (e.g., Pycraft (1901); de Beer (1937); Witmer and Martin (1987); Bühler et al. (1988); Zusi and Livezey (2006)) indicating the absence or truncation of the process of pterygoid segmentation in certain groups. Nonetheless, the lack of advanced three-dimensional imaging to illustrate these small elements in detail across a large sample of neornithine diversity has thus far precluded an unambiguous assessment of the phylogenetic distribution of pterygoid segmentation across major neornithine subclades.

**Main text table 1:** Table showing our five hypothetical stages of pterygoid segmentation

|  |  |
| --- | --- |
| **Stage 1** | Unsegmented hemipterygoid |
| **Stage 2** | Partially segmented hemipterygoid |
| **Stage 3** | Fully segmented hemipterygoid |
| **Stage 4** | Partial fusion of hemipterygoid to caudal palatine |
| **Stage 5** | Complete fusion of hemipterygoid to caudal palatine |

### Palaeognathae

Despite consensus that pterygoid segmentation does not occur in palaeognaths (e.g., Pycraft (1901); Balouet (1983); Bühler et al. (1988); Elzanowski (1995)), the term ‘hemipterygoid’ sometimes appears in descriptions of the palaeognath palate, including in works suggesting that neoteny drove morphological transitions in the palaeognath palate (e.g., de Beer (1956)). More recent studies suggest that neoteny does not explain the peculiarities of the palaeognath palate, as the morphology of the pterygoid-palatine complex (PPC) of adult palaeognaths does not resemble that of developing neognaths (e.g., Gussekloo and Bout (2002); Zusi and Livezey (2006); Plateau et al. (2026) (but see the discussion of the tinamou in Gussekloo and Bout (2002))). The non-neotenic nature of the palaeognath palate is further supported by evidence from the ossification sequence of the palaeognath skull (Maxwell, 2009).

None of the palaeognaths surveyed as part of our study exhibited any direct evidence for pterygoid segmentation such as an identifiable, distinct hemipterygoid, nor significant ontogenetic change in the shape of the pterygoid and palatine consistent with pterygoid segmentation. However, all the examined palaeognaths bore various forms of a rostral projection of the pterygoid, which, based on our observation of mature specimens, does not detach from the pterygoid during ontogeny. Indeed, in some palaeognaths, such as tinamous, this rostral projection of the pterygoid is somewhat reminiscent of the first stage of pterygoid segmentation (figs. 6, 7, 21, 22); however, the rostral projection of palaeognaths is highly morphologically variable (figs. 5-7), which contrasts with the relatively conservative morphology of the hemipterygoid of neoavians (figs. 14-16).

Both the rostral end of the palaeognath pterygoid (e.g., de Beer (1956); McDowell (1978)) and the portion of the palaeognath pterygoid contacting the vomer (Núñez León, 2015) have occasionally been inferred to be homologous with the hemipterygoid of neognaths. However, until a more detailed assessment of the pre-hatching development of the pterygoid and associated palatal regions of a wider taxonomic sample of palaeognaths and neognaths is undertaken, we regard the potential homology of the rostral projection of palaeognaths with the unsegmented hemipterygoid of neoavians as incompletely substantiated.

Pterygoid morphology is ontogenetically conservative across our palaeognath taxon sample (fig. 7). Conversely, the morphology of the pterygoid exhibits a high degree of interspecific variability (fig. s2) ((Benito et al., 2022); (Plateau et al., 2026)), particularly with regard to the pterygoid-palatine contact and the morphology of the rostral region of the pterygoid (figs. 5, 6, s2). In tinamous and the moa *Megalapteryx*, the rostral projection of the pterygoid contacts the caudal region of the palatine, possibly representing a morphological synapomorphy of the hypothesised tinamou-moa clade proposed on the basis of molecular sequence data (e.g., Phillips et al. (2010); Baker et al. (2014); Kimball et al. (2019)). By contrast, in *Struthio* and immature *Dromaius* this rostral projection does not contact the caudal region of the palatine, instead diverging from it medially.

The pterygoid of the immature kiwi (*Apteryx australis*) is especially distinct from that of other immature palaeognaths (fig. s2) exhibiting a conspicuous rostral bifurcation previously reported by Parker (1891), Archey (1941) and McDowell (1948). Jollie (1957) interpreted the medial process of the kiwi pterygoid as the anteropterygoid (hemipterygoid), and we agree that it does resemble the unsegmented hemipterygoid of immature neoavians in several ways. Firstly, the medial position of this process (fig. 5) resembles the position of the hemipterygoid of neoavians (figs. 14-16); in addition, the medial process is blade-like in appearance and narrows rostrally, reminiscent of the morphology of neoavian hemipterygoids. Likewise, a section of the medial process of the pterygoid of *Apteryx* overlies part of the caudal region of the palatine, in a similar manner to how the hemipterygoid and the palatine contact one-another in neoavians. Nevertheless, the nature of the contact between the pterygoid, palatine and vomer in *Apteryx* contrasts with that of neoavians, where the hemipterygoid largely contacts the dorsolateral region of the palatine (with a short contact with the caudal region of the vomer in some cases), while the medial process of the pterygoid in *Apteryx* largely overlaps the caudalmost region of the vomer. The potential homology of these structures thus remains ambiguous at present, leaving open an interesting avenue for future developmental studies to explore.

The caudal region of the palaeognath palatine is fairly conservative through ontogeny (fig. 7); however, in *Dromaius* the palatine undergoes a considerable degree of ontogenetic dorsoventral expansion during ontogeny (fig. 7). As noted by others (e.g., Plateau et al. (2026)), palatine morphology is highly variable across palaeognaths (fig. s1), ranging from more gracile, elongate palatines such as that of the tinamou *Nothoprocta perdicaria*, to more complex, robust morphologies such as that of the moa *Megalapteryx*.

Unusually among crown-group birds, palaeognaths exhibit immobile palatine-pterygoid articulations resisting prokinesis, with both elements, as well as the vomer, fused into a single solid structure in adult birds (e.g., Pycraft (1900); McDowell (1948); Widrig and Field (2022)). Although palatal fusion appears to occur relatively late in ontogeny, particularly in large-bodied palaeognath taxa (exemplified by the lack of fusion in the subadult *Struthio* and *Dromaius* sampled here), it is conceivable that this fusion and the loss of palatal mobility could be directly related to the absence of a discernible hemipterygoid and pterygoid segmentation. For instance, the onset of palatal fusion may prevent pterygoid segmentation in palaeognaths; alternatively, the loss of pterygoid segmentation may have facilitated the evolution of palaeognath palatal fusion. Assessing whether a causal relationship exists between these processes—or whether they are related at all—remains challenging to ascertain, not least because of the extraordinary palatal disparity exhibited within palaeognaths.

The morphology of the palatine-pterygoid contact surface of palaeognaths is highly variable (figs. 5, 6). For instance, in *Struthio* the pterygoid process of the palatine wraps around the convex ventral pterygoid, whereas in *Dromaius* the palatine and pterygoid lie alongside one-another, forming a rostrocaudal longitudinal contact. Contrastingly, in *Apteryx* the condition is more complex. The medial articular shelf of the palatine wraps around the ventral surface of the pterygoid’s medial process, and the caudolateral flange of the palatine inserts into the groove separating the medial and lateral processes of the pterygoid. Considerable variation is even evident among tinamous (Tinamidae): in *Tinamus*, the slightly concave dorsal surface of the palatine wraps around the overlying pterygoid, while *Nothoprocta* and *Crypturellus* exhibit relatively flat and elongate contact surfaces between the pterygoid and palatine (fig. 6). Notably, the latter condition is shared with the moa *Megalapteryx*, constituting another potential morphological synapomorphy of the tinamou-moa clade hypothesised on the basis of genomic data (e.g., Phillips et al. (2010); Baker et al. (2014); Kimball et al. (2019)).

These and previous observations of the highly variable morphologies of the palatine and pterygoid of palaeognaths challenge the very concept of a single ‘palaeognathous’ palatal condition that accurately encompasses the palatal architecture of Palaeognathae. This conclusion is concordant with previous investigations highlighting morphological disparity in the palaeognath palate (e.g., Benito et al. (2022); Plateau et al. (2026)) and recall earlier observations highlighting pronounced morphological variation in this clade (e.g., McDowell (1948)). It has been proposed that the reduced palatal kinesis in palaeognaths might be related to their high palatal disparity, as the release of functional constraints on pterygoid and palatine shape related to supporting palatal kinesis may have allowed these bones to vary more freely (Plateau et al., 2026). Our observation of negligible ontogenetic change of the rostral region of the pterygoid and the caudal region of the palatine is consistent with the apparent absence of pterygoid segmentation in crown group palaeognaths, yet whether segmentation represents a crown bird plesiomorphy lost along the palaeognath stem lineage remains unclear, and will necessitate future work on pre-hatching palate development (see Núñez León (2015)).

### Anseriformes

Previous literature has been ambiguous regarding the existence of a hemipterygoid associated with pterygoid segmentation in Anseriformes. For example, Pycraft (1901) described the hemipterygoid of Anseres as a rostral projection of the pterygoid that does not segment (yet, in the same publication also referred to the hemipterygoid as being lost by atrophy in Anseres) and later Pycraft (1902) described the hemipterygoid as completely supressed in Anseres. No specific ontogenetic information was provided in support of these statements. Bühler et al. (1988) stated that pterygoid segmentation is suppressed in anatids, but did not explicitly comment on the presence or absence of a ‘hemipterygoid’. More recently, in a study that included both immature and mature specimens, Zusi and Livezey (2006) interpreted the hemipterygoid (their ‘*pars palatina pterygoidei’*) as absent in Galloanserae, finding no evidence for a reduced or vestigial hemipterygoid.

None of the anseriforms surveyed as part of our study exhibit any direct evidence for post-hatching pterygoid segmentation such as an identifiable, distinct hemipterygoid or significant changes in the shape of the pterygoid and palatine during ontogeny. This is concordant with developmental studies that have not documented the presence of a hemipterygoid, or identified evidence for pterygoid segmentation (e.g., Maxwell (2008a); Mitgutsch et al. (2011); Arnaout et al. (2025); but see Núñez León (2015)).

Amongst Anseriformes, the articular regions of the palatine and the pterygoid of Anatidae undergo very minor morphological change across post-hatching ontogeny, and beyond some limited rotation of both elements relative to one-another and to the skull, their arrangement is conserved across post-hatching development. This is consistent with the absence of a distinct hemipterygoid and pterygoid segmentation, and indicates that, unlike in Neoaves, the pterygoid process of Anatidae (and presumably Anseres more generally) is formed exclusively by the palatine. Furthermore, the conspicuous rostral projection of the pterygoid present in immatures is retained across ontogeny and is found in adult anatids, yet it changes from a blade-like tapered process in immatures, to a wider cylindrical process in adults. Whether the retention of this rostral projection through ontogeny is of any biomechanical relevance may represent a topic of interest for future functional research. Importantly, both the morphology and position of this rostral projection strongly resemble that of the unsegmented hemipterygoid of young Stage 1 neoavians (figs. 14, 16). Based on the morphological resemblance and topological similarities between these structures in Anseres and Neoaves, and given the very limited degree of post-hatching ontogenetic change in the palate of Anatidae, we hypothesise that the rostral projection of anatids is homologous with the unsegmented hemipterygoid of neoavians, with anatids failing to undergo the process of pterygoid segmentation. This observation adds further support to previous inferences regarding the absence of separation of the rostral projection of the pterygoid in anatids (e.g., Pycraft (1901); Bühler et al. (1988)). However, we acknowledge that future developmental research is needed to rigorously evaluate this hypothesis of homology.

Palate ontogeny amongst the surveyed members of Anhimidae appears to be clearly distinct from that of Anatidae, exhibiting more marked changes in the morphology of the pterygoid and the palatine. Although obscured by incomplete ossification in the examined immature *Chauna chavaria* specimen, it is apparent that the rostral portion of the pterygoid experiences a marked elongation across post-hatching ontogeny, though the arrangement and morphology of the palatine-pterygoid contact remain mostly unchanged.

Importantly, despite their limited diversity, anhimids exhibit a notable degree of disparity in palatal anatomy, especially compared to other extant Anseriformes. Both the palatine and the pterygoid of the two adult *Chauna* specimens surveyed (belonging, respectively, to *C. chavaria* and *C. torquata*) are markedly different, particularly regarding their palatine-pterygoid articulation, while the pterygoid of *Anhima* is considerably more elongate than that of *Chauna* or indeed any other galloanseran observed.

Unlike the case in Anseres, anhimids lack a distinct elongated rostral projection overlapping the dorsolateral surface of the palatine. Instead, their pterygoids exhibit a short-stalked and robust rostral projection (fig. 9) which incorporates the articular surface for the palatine and wraps around most of the pterygoid process (or its entirety in *C. torquata*). In *Chauna*, this dorsolateral margin of the pterygoid’s rostral projection extends into a very short overlap on the lateral surface of the palatine, while this overlap is narrower and much longer rostrocaudally in *Anhima*, reminiscent of the rostral projection of Anseres. In addition, the basipterygoid process facet of all surveyed adult anhimids is markedly caudal to the palatine-pterygoid articulation, rather than immediately adjacent to it, as in all other Anseriformes examined. Remarkably, it is positioned much closer to the rostral end of the pterygoid in the immature *C. chavaria* examined, which could suggest the rostral projection of anhimids lengthens through ontogeny.

Whether any portion of the anhimid pterygoid’s rostral projection is homologous with the rostral projection of Anseres and the unsegmented hemipterygoid of Neoaves is unclear. In both immature Neoaves and Anatidae the hemipterygoid (in the former) and rostral projection of the pterygoid (in the latter) is adjacent to, but does not contain the palatine-pterygoid articulation, unlike in anhimids, and does not change in its relative length across ontogeny. However, it is possible that only the dorsal portion of the rostral projection of anhimids is homologous with the rostral projection of other Anseriformes, as it shares a similar position and, especially in *Anhima*, is similar in its general morphology. If this interpretation is accurate, the morphology of the rest of the anhimid rostral projection might correspond to a unique rearrangement of the rostral region of the pterygoid. Importantly, the presence of similar rostral projections in megapode galliforms raises new questions about the ancestral condition for both Galliformes and Anseriformes, and the specific evolutionary transformations that have given rise to the distinctive palates of extant galloanserans. Our results suggest that the anseriform condition may represent a truncated or incomplete version of pterygoid segmentation, which could be explained by a heterochronic shift. Whether this interpretation proves accurate, and if so, determining whether the anseriform condition is paedomorphic—or the neoavian condition peramorphic with respect to the ancestral condition for crown birds or Neognathae—will require further developmental work.

#### Potential homology of the anseriform rostral projection and neoavian hemipterygoid

Given the evidence for homology of the rostral projection of the pterygoid of anatids with the hemipterygoid of neoavians as outlined in the previous section, we recommend the term ‘hemipterygoid process’ to describe the rostral projection of the pterygoid of Anseres, and especially Anatidae. This term is also potentially applicable to at least part of the rostral projection of the pterygoid of anhimids (see above), yet whether the entire structure is homologous with the hemipterygoid remains unclear. We suggest refraining from simply using the term ‘hemipterygoid’ to describe the rostral projection of anatids (and potentially part of the rostral projection of anhimids), because according to our data it does not undergo full pterygoid segmentation during ontogeny (i.e. it terminates at a point equivalent to Stage 1 in Neoaves (see figures 21, 22)). Our application of the term ‘hemipterygoid process’ therefore differs from the ‘mesopterygoid process’ (Parker, 1872), which referred to the unsegmented portion of the hemipterygoid (his mesopterygoid) that was capable of splitting off during pterygoid segmentation.

### Galliformes

Differing views on the presence of a hemipterygoid and pterygoid segmentation in Galliformes have been documented in the literature. For example, de Beer (1937) described the hemipterygoid of *Gallus* as a process projecting forwards from the rostral end of the pterygoid that does not detach. While no specific information was provided regarding the ontogenetic stage to which this observation pertains, de Beer had earlier stated that most membrane-bones arise during the middle of the second week of incubation, perhaps providing some indication. By contrast, Parker (1869) described the hemipterygoid (his ‘mesopterygoid’) as absent in *Gallus* based on observations of a two-day-old post-hatching specimen. Without providing any ontogenetic information, Pycraft (1901) described the hemipterygoid of the “Galli” as a peg-like structure projecting from the rostrodorsal edge of the pterygoid that does not segment (yet, in this same publication also stated that the hemipterygoid was lost by atrophy in the “Galli”). Subsequently, Pycraft (1902) described the hemipterygoid as completely supressed in the “Galli”, though again did not provide specific information on the ontogenetic stage of the specimens observed. Notably, Jollie (1957) documented the pterygoid (‘posteropterygoid’) and the hemipterygoid (‘anteropterygoid’) as ossifying separately in *Gallus*, with the hemipterygoid already being fused to the palatine upon its first appearance (observed on the eleventh day of incubation). This scenario would indicate an embryonic origin of the hemipterygoid formed without post-hatching pterygoid segmentation. Zusi and Livezey (2006) described the hemipterygoid (their *‘pars palatina pterygoidei’*) of Galloanserae as absent (their study included immature and adult specimens), while Núñez León (2015) identified a possible hemipterygoid during the pre-hatching development of *Gallus gallus* (the latter implying an embryonic origin of the hemipterygoid). However, other recent developmental studies of the embryonic skull in galliforms do not explicitly discuss the presence or absence of the hemipterygoid or pterygoid segmentation (e.g., Maxwell (2008b); Arnaout et al. (2021); Arnaout et al. (2025)). Thorough imaging of pre-hatching embryos at relatively early stages of development will be necessary to unambiguously resolve the question of whether a transient hemipterygoid is present in at least some members of Galliformes (see Núñez León (2015)).

Megapodes (Megapodiidae) represent the sister taxon to all other extant Galliformes (e.g., Prum et al. (2015)). They are distinct from other examined galliforms in that mature specimens bear a short rostral projection of the pterygoid, the morphology of which is remarkably variable among the megapode taxa surveyed (fig. 12). This projection is particularly conspicuous in *Leipoa,* in which it is robust and narrows subtly to produce a rounded rostral tip abutting the pterygoid process of the palatine, while those of *Alectura* and *Megapodius* are rostrocaudally shorter. In *Alectura*, this rostral projection partially overlies the dorsal surface of the pterygoid process of the palatine, similar to the relationship between the pterygoid process of the palatine and the hemipterygoid process (in Anatidae) and the unsegmented hemipterygoid (Stage 1, Neoaves) (figs. 8, 12, 14, 16). By contrast, in *Megapodius* this projection bears a rostrally positioned cup-like cotyle which almost completely encircles the caudal palatine, strongly reminiscent of the cotyle-bearing rostral projections of adult anhimids.

The previously overlooked disparity in pterygoid morphology in both megapodes and anhimids is intriguing, as is the close morphological similarity between the rostral projection of *Megapodius* to that of *Chauna* and *Anhima* (figs. 9, 12). This is especially notable considering that both megapodes and anhimids represent the sister taxa to the remainder of Galliformes and Anseriformes, respectively (e.g., Prum et al. (2015)). These shared morphologies suggest that megapodes and anhimids may capture several aspects of the plesiomorphic galloanseran palatal condition that were convergently lost along the stem lineages of the clade of non-megapodiid galliforms (recently named Orbigalli by Chen et al. (in review)) and the non-anhimid anseriform clade (i.e. Anseres). However, strong inferences about the morphology of the pterygoid in the last common ancestor of Galloanserae are complicated by the observation that the hemipterygoid processes of Anseres and the unsegmented hemipterygoid of Stage 1 Neoaves exhibit similarities that are unshared with either Anhimidae or Megapodiidae. Moreover, several aspects of the morphology of the rostral pterygoid projection that are also reminiscent of neoavian unsegmented hemipterygoids are exclusively shared by *Anhima* and Anseres (but not *Chauna*), and *Alectura* and Anseres (but not other megapodes). Regardless of the specific plesiomorphic pterygoid morphology for Galloanserae, the observed lack of pronounced ontogenetic change in the caudal region of the palatine of *Megapodius* (fig. 12) supports the absence of pterygoid segmentation in megapodes and we consider it likely that at least part of the rostral projection of the megapode pterygoid is homologous with the unsegmented hemipterygoid of neoavians (similar to the case discussed above for Anhimidae). We therefore suggest applying the term ‘hemipterygoid process’ to refer to the rostral projections of megapode pterygoids.

We only observed moderate ontogenetic changes in the palatal morphology of non-megapodiid Galliformes, generally restricted to the caudal palatine. Specifically, while the shape of the pterygoid process of the palatine among the examined immature representatives of Phasianoidea was conservative and generally cylindrical, the pterygoid processes of the surveyed adults exhibit greater morphological disparity (fig. 13). In contrast, the morphology of the rostral articular surface of the pterygoid was conserved across ontogeny, with a cup-like articular cotyle for the pterygoid process of the palatine present early in development and retained in adults. This cotyle becomes more complex in mature specimens, developing clear raised margins: in *Gallus*, it is clearly enclosed on at least its dorsal and lateral margins, whereas in *Numida* it is enclosed on all margins apart from the ventral margin (fig. 13).

In a broad taxonomic sample of non-megapodiid galliforms, we did not identify evidence for pterygoid segmentation or any distinct structure diagnosable as a hemipterygoid. Unlike in Megapodiidae, Anhimidae, and Anatidae, none exhibited a rostral projection of the pterygoid potentially homologous with a hemipterygoid at any point of their post-hatching development, and the morphology of the rostral pterygoid was essentially unchanged through post-hatching ontogeny (fig. 13). As such, non-megapodiid galliforms apparently lack a stage equivalent to Stage 1 of pterygoid segmentation. As mentioned above, Jollie (1957) reported that the hemipterygoid in *Gallus* forms initially as an independent element which immediately fuses to the palatine at an early embryonic stage, never forming part of the pterygoid. If this scenario is accurate and holds true for other non-megapodiid galliforms, the pterygoid process of the palatine would already incorporate the hemipterygoid at hatching, and would be homologous with this structure in Neoaves (see discussion below). Indeed, if this process is restricted to an early point in pre-hatching ontogeny (Jollie (1957)), it may simply be unobservable in the late-stage embryos and post-hatching immature specimens examined here. Crucially, further embryological work will be required to confidently assess whether a structure homologous with the neoavian hemipterygoid is present at any point during the pre-hatching development of non-megapodiid Galliformes.

### Neoaves

The transient existence of a discrete hemipterygoid has previously been documented in various neoavian taxa, most extensively in the works of Parker and Pycraft (e.g., Parker (1872); Parker (1875); Pycraft (1903); Pycraft (1907b)). Though these studies present detailed images of palatal morphology, commonly in ventral/lateral views, interpretation of the hemipterygoid is hindered by its dorsal position relative to the pterygoid and palatine. The present study therefore represents the first attempt to illustrate hemipterygoid morphology in detail using a broad sample of extant bird species.

In immature neoavians in which the hemipterygoid is observable as a distinct element prior to its fusion with the palatine—either partially segmented (Stage 2) or discrete and fully segmented (Stage 3) (table 2)—we observed that the rostral region of the hemipterygoid predominantly contacts the dorsomedial surface of the caudal region of the palatine. Sometimes this caudal region of the palatine bears a pronounced, albeit shallow, facet to receive the hemipterygoid (best exemplified by the immature penguin *Spheniscus demersus* (fig. s14)). Although the nature of the hemipterygoid-palatine contact and the arrangement of palatal elements is remarkably similar among surveyed immature neoavians (figs. 14-16), the morphology of the hemipterygoid at Stages 2 and 3 is variable among neoavian subclades. The taxa surveyed exhibit hemipterygoids ranging from relatively elongate, slender elements in Ardeidae to more robust, mediolaterally broad elements in Procellariidae and Sulidae. We also observed a high degree of variability in the relative sizes of the hemipterygoid and pterygoid body, with the hemipterygoid of the condor *Vultur* being longer than the entire pterygoid body, while that of the avocet *Recurvirostra* was proportionally short, approximately one third of the entire pterygoid body length. Variability was also observed in the extent to which the hemipterygoid extends rostrally across the palatine, ranging from the hemipterygoid covering ∼15% of the palatine in the flamingo *Phoenicopterus* to ∼25% in the condor *Vultur*.

**Main text table 2:**
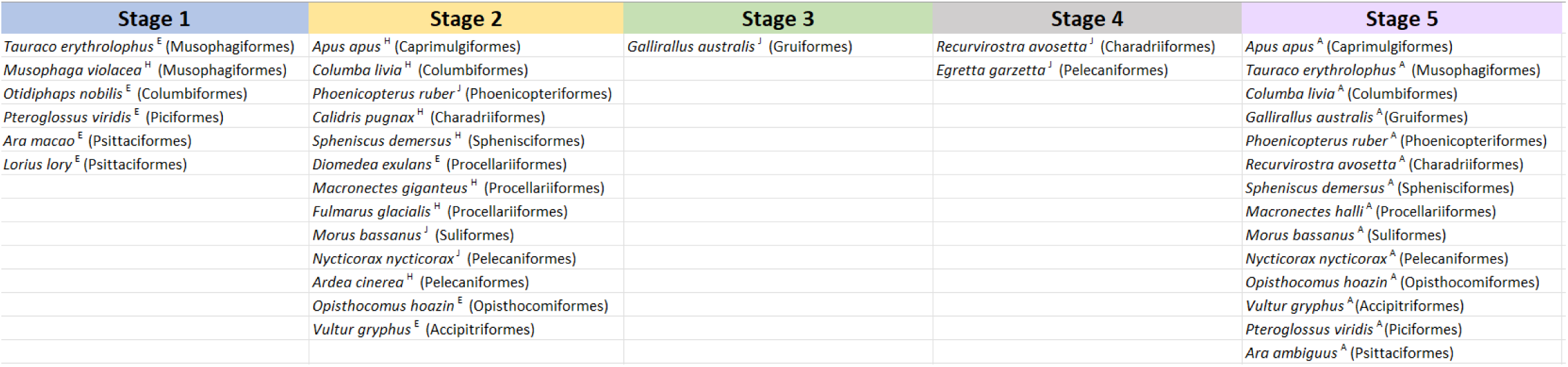
Table showing categorizations of each examined neoavian specimen into one of five distinct, hypothetical stages of pterygoid segmentation. 1: unsegmented hemipterygoid; 2: partially segmented hemipterygoid; 3: fully segmented hemipterygoid; 4: partial fusion of hemipterygoid to caudal palatine: 5: complete fusion of the hemipterygoid to the caudal palatine. Footnotes: Superscript E, H, J and A correspond to embryo, hatchling, juvenile and adult, respectively.

Our ontogenetic observations illustrate the process of pterygoid segmentation, capturing all stages from the hemipterygoid’s detachment from the pterygoid to its fusion with the caudal palatine during ontogeny (figs. 21, 22). These observations reveal that, among extant birds, the process of pterygoid segmentation—at least as a post-hatching phenomenon—is phylogenetically restricted to Neoaves. All surveyed adult neoavians showed the absence of a distinct hemipterygoid, and consistently illustrated a clear change in the morphology of the pterygoid process of the palatine throughout ontogeny consistent with its incorporation of the hemipterygoid (figs. 17-20). Importantly, our observations reveal that the hemipterygoid of most neoavians forms only the dorsal portion of the pterygoid process, with the ventral portion of this structure formed by the palatine at an early point in development. As such, in these instances, the adult pterygoid process of the palatine constitutes a structure of hybrid origin, which sometimes exhibits two distinct condyle-like structures originating from the hemipterygoid (dorsally) and the palatine (ventrally). Thus, in most adult neoavian birds this pterygoid-palatine joint consists of the intrapterygoid and the pterygoid/palatine contact (the “*articulationes pterygopalatina et intrapterygoidea”* of Zusi and Livezey (2006)) (but see Zusi and Livezey (2006) for a summary of other contact types). This hybrid articulation of Neoaves is therefore only partially homologous with those of other neognaths. In Anseres, and likely in Anhimidae and Megapodiidae, the rostral projection of the pterygoid does not segment (hence we refer to these processes as ‘hemipterygoid processes’), such that in Anseres the mobile palatal articulation is formed exclusively between the palatine and the pterygoid, while in Anhimidae and Megapodiidae the articulation is either between the pterygoid and palatine or between the hemipterygoid and palatine. In Galliformes a single articular condyle is present on the palatine. If, as suggested by Jollie (1957) this condyle derives from the hemipterygoid embryonically, it would imply that galliforms exclusively exhibit an intrapterygoid articulation, homologous with the dorsal portion of the articulation in Neoaves (but apparently formed in the absence of post-hatching pterygoid segmentation).

The nature of the mobile palatal articulation is, in any case, remarkably conserved across Neoaves, with only minor variation across the surveyed subclades (apart from parrots as noted below) (figs. 14-16). This observation is in line with the suggestion that variation in the neoavian suspensory system is limited by strong functional and developmental constraints (Plateau et al., 2026). This comparatively low degree of disparity in Neoaves, an extraordinarily diverse clade comprising over 95% of extant bird diversity (e.g., Somogyi et al. (2026)) stands in stark contrast to the high degree of variability of the palatine and pterygoid in palaeognaths (figs. 5, 6, s1, s2) and the divergent pterygoid configurations of galloanserans (figs. 8, 11, s4, s5, s7), both of which are far less species-rich than Neoaves. The bizarre, highly derived palatal anatomy observed in parrots (figs. 16, 20, s9, s11) is strikingly divergent with respect to that of all other Neoaves and is likely related to the well-documented and extreme mobility and kinetic capabilities of the psittaciform beak (e.g., Tokita (2003); Young et al. (2023)).

The timing and duration of the process of pterygoid segmentation is likely variable across neoavian diversity, and may covary in line with the altricial-precocial spectrum (Ducatez & Field, 2021). However, it is remarkable that only one specimen from our broad sample, an immature *Gallirallus australis*, exhibited an isolated, fully independent (discrete) hemipterygoid detached from both the pterygoid and the palatine (Stage 3). The hemipterygoid of all other surveyed immature neoavians was either still attached to the pterygoid (either unsegmented at Stage 1, or partially segmented at Stage 2), or partially fused to the palatine (Stage 4). This observation may suggest that a transient, fully independent hemipterygoid only appears during a brief window of post-hatching development; alternatively, and perhaps more likely, it may be the case that fusion of the hemipterygoid and palatine commences prior to the full separation of the hemipterygoid from the pterygoid in many taxa, bypassing Stage 3 entirely. Indeed, our examined immature *Egretta* specimen (suppl. table 1) may illustrate the latter case. However, future work will be necessary to assess whether and how frequently Stage 3 is bypassed, and the extent to which differences in developmental mode may be related to this phenomenon.

### FOSSILS AND THE ORIGIN OF PTERYGOID SEGMENTATION

Although both the presence of a discrete–though transient–hemipterygoid and the complete process of pterygoid segmentation are unique to Neoaves among extant birds, fossil evidence from near-crown stem birds suggests that the origin of a hemipterygoid most likely predates the origin of pterygoid segmentation. Indeed, discrete hemipterygoids have been recognised in adult representatives of the ornithurine clades Ichthyornithes (Torres et al., 2021) and Hesperornithes ((Elzanowski, 1991); (Bell & Chiappe, 2020)). In contrast to the hemipterygoids of immature neoavians and the putative hemipterygoid processes of megapodes and anseriforms (this study), the hemipterygoids of non-crown ornithurines are highly elongate and overlap along much of the dorsomedial length of the palatine ((Bell & Chiappe, 2020); (Torres et al., 2021); (Elzanowski, 1991)). In Ichthyornithes and Hesperornithes, the position of these elongate hemipterygoids is generally comparable to that of immature neoavians at developmental Stages 2-4 (figs. 21, 22). Although these hemipterygoids are unfused to the pterygoid process of the palatine, the hemipterygoid forms the main or only articulation point for the pterygoid in Ichthyornithes ((Torres et al., 2021); (Benito et al., 2022) and Hesperornithes ((Elzanowski, 1991); (Bell & Chiappe, 2020)). As such, these non-neornithine ornithurines appear to exclusively exhibit a pterygoid-hemipterygoid articulation, though the nature of this contact in Hesperornithes has been difficult to reconstruct (see Bell and Chiappe (2020) and Wilken et al. (2025)). This arrangement is perhaps most similar to the palatal joint of non-megapodiid galliforms, and contrasts with the condition observed in anatids (in which the contact is directly between the pterygoid and the palatine), anhimids and megapodes (with an uncertain contact between either the pterygoid and the palatine or between the hemipterygoid and the palatine) and Neoaves (which generally exhibit a hybrid articulation comprising both intrapterygoid and pterygoid-palatine contacts).

On balance, we consider it likely that the hemipterygoid originated within Avialae as an independent palatal element arising from a discrete ossification centre (i.e. not formed through pterygoid segmentation), that persisted into adulthood. This hypothesised embryonic origin would be consistent with that proposed for *Gallus* ((Jollie, 1957); (Núñez León, 2015)), the lapwing *Vanellus chilensis* and the coot *Fulica armillata* (Núñez León, 2015). Based on the known distribution of hemipterygoids among stem birds, we hypothesise that this independent hemipterygoid arose prior to the origin of Ornithurae (fig. 23); however, as with other aspects of detailed cranial morphology (e.g., Field et al. (2018); Hu et al. (2022); Chiappe et al. (2024)) precisely determining the phylogenetic position of its origin is precluded by the rarity of complete, well-preserved Mesozoic avialan skulls and palates. Only later, either along the crownward portion of the avian stem lineage or among crown birds, did the hemipterygoid fuse to the pterygoid body; subsequently, at least along the neoavian stem lineage, pterygoid segmentation may have arisen from the sequential developmental detachment of the hemipterygoid from the pterygoid body, and fusion with the palatine. Although it is conceivable that the hemipterygoid of non-crown ornithurines also detached from the pterygoid through ontogeny as in Neoaves (i.e. undergone pterygoid segmentation – see Torres et al. (2021)), all known ichthyornithine and hesperornithine hemipterygoids are very large relative to these elements in Neoaves and are unfused to either the pterygoid or palatine. We therefore consider the scenario in which the origin of the hemipterygoid was decoupled from the origin of pterygoid segmentation as most plausible. Together with the recent recognition that a neognath-like mobile palatal articulation is likely ancestral for crown birds, rather than representing a neognath synapomorphy ((Torres et al., 2021); (Benito et al. (2022); (Field et al., 2025); (Benito et al., 2026), but see Wilken et al. (2025); Wilken et al. (2026)) this interpretation suggests that neognath-like prokinesis arose independently of pterygoid segmentation.

**Figure 23:**
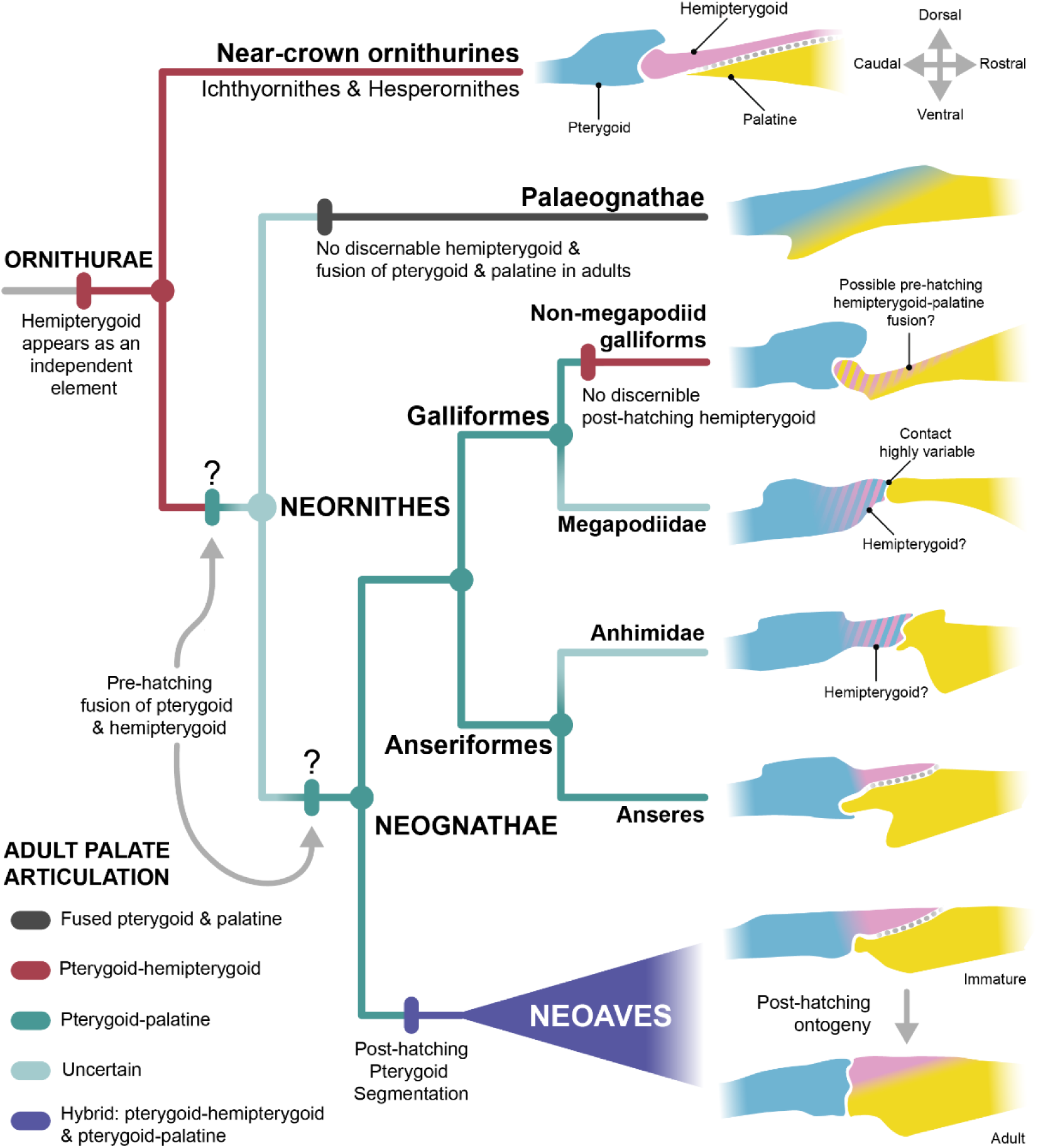
Simplified phylogenetic tree illustrating the evolutionary origin of a discrete hemipterygoid and pterygoid segmentation. We illustrate our preferred evolutionary scenario based on available data, but see discussion for alternative hypotheses. Pink banding marks structures whose identity is uncertain but that might correspond to the hemipterygoid based on current evidence. Tree branches and nodes where the adult palate articulation is indicated as ‘Uncertain’ (light green) might correspond to either pterygoid-hemipterygoid, hemipterygoid-palatine or pterygoid-palatine articulations; current evidence prevents us from discerning among these alternatives. Flat and overlapping, presumably immobile articulations between the hemipterygoid and the palatine are indicated by dashed grey lines to differentiate them from condylar and hinge-like mobile articulations.

Elucidating when the process of segmentation evolved is complicated by our hypothesis that the rostral projections of Anseriformes and Megapodiidae (our ‘hemipterygoid processes’) are homologous with the hemipterygoids of Neoaves and non-crown ornithurines. The presence of this putative hemipterygoid process in anseriforms and megapodes suggests that fusion between the hemipterygoid and the pterygoid is likely plesiomorphic for Galloanserae. This hypothesis is supported by the presence of short and robust rostral projections in stem-group Anseriformes like *Nettapterornis* ((Olson, 1999); (Benito et al., 2022)) and *Presbyornis,* which most likely correspond to hemipterygoid processes and are especially reminiscent of the condition in megapodes (fig. 24). Furthermore, these fossil taxa appear to retain a range of other galloanseran and neornithine cranial plesiomorphies (e.g., Kuo et al. (2023); Field et al. (2024); Arnaout et al. (2025); Arnaout et al. (2026); Kuo et al. (2026)).

**Figure 24:**
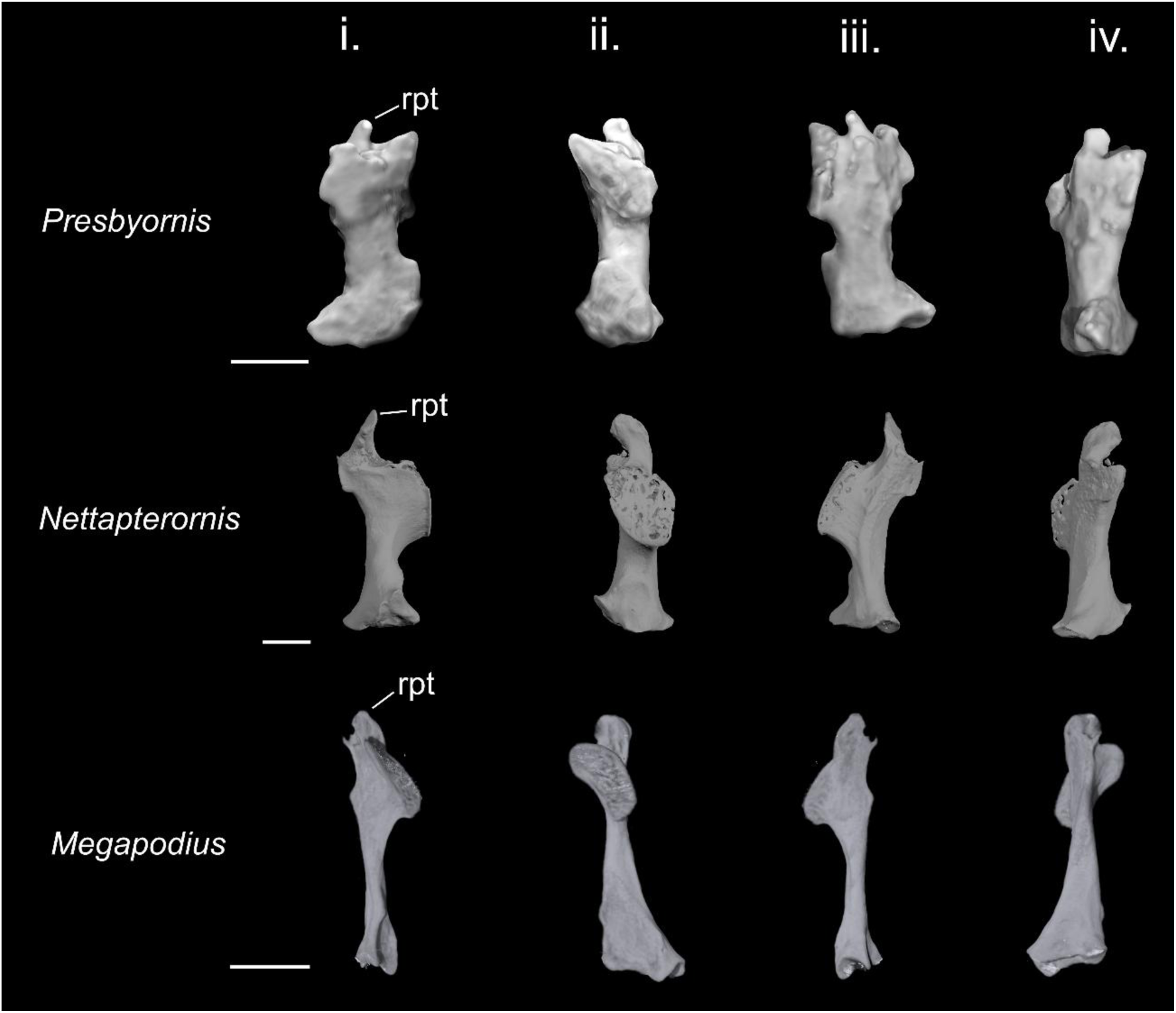
Comparison of the rostral projection of the pterygoid in two fossil total-group Anseriformes (*Presbyornis* (USNM PAL 299846) and *Nettapterornis* (NHMUK PVA 5922)) and the mature megapode *Megapodius nicobariensis*. The pterygoid represents a left element for *Presbyornis* and right for the latter two taxa. Abbreviations: rpt, rostral projection of the pterygoid. i - iv represent the following orientations: i.- dorsal, ii.- medial, iii.- ventral and iv.- lateral. All scale bars equal 2.5 mm.

Important questions remain regarding the evolution of the hemipterygoid process characteristic of Galloanserae considering alternative viable interpretations of macroevolutionary transformation series. The galloanseran hemipterygoid process may represent an intermediate, ‘incomplete’, step in the evolution of pterygoid segmentation, in which case we would interpret a galloanseran-like hemipterygoid process fused to the pterygoid to have evolved along the stem lineage of Neognathae, with full detachment of the hemipterygoid and its fusion to the palatine evolving along the stem lineage of Neoaves. By contrast, if full pterygoid segmentation represents the ancestral condition for Neognathae, the hemipterygoid process would most likely represent an autapomorphy arising along the galloanseran stem lineage from truncation of the process of segmentation at Stage 1 (figs. 21, 22). However, given that we only identify direct evidence for the full process of pterygoid segmentation in neoavians, we consider it most likely that pterygoid segmentation represents a synapomorphy of Neoaves (fig. 23).

Other fossil birds of more uncertain affinities exhibit rostral projections potentially homologous to the galloanseran hemipterygoid process, including the putative crown bird *Vegavis* from the Maastrichtian of Antarctica ((Clarke et al., 2016); (Irazoqui et al., 2026)) and the pseudotoothed Pelagornithidae ((Harrison & Walker, 1976); (Mourer-Chauviré & Geraads, 2008); (Benito et al., 2022)). The rostral projections in these taxa are minute, however, often representing little more than a raised crest on the rostral surface of the pterygoid. Clarifying the phylogenetic position of these enigmatic fossil taxa remains the subject of considerable active research (e.g., Bourdon (2005); Mayr (2011); Torres et al. (2025)), and resolving the question of whether these taxa represent members of major crown bird subclades, or even represent crown birds at all (e.g., Field (2025); Crane et al. (2026)), will clarify the sequence by which a minute hemipterygoid process, and the complete process of pterygoid segmentation, evolved.

Notably, non-megapode galliforms are unique among extant neognathous birds in lacking any unambiguously discernible vestige of a hemipterygoid throughout their post-hatching ontogeny. Jollie (1957) suggested that a hemipterygoid (which ossifies separately to the pterygoid) is present very early in the embryonic ontogeny of *Gallus* and undergoes immediate fusion with the palatine (also see Núñez León (2015)). The pterygoid of cracids and Phasianoidea, which is characterised by a tubular shape, a large concave articular cotyle, and the absence of a rostral projection, is remarkably similar to that of the ichthyornithine *Janavis,* while the morphology of the pterygoid process of the palatine in these galliforms is very reminiscent of the caudal region of the hemipterygoid of *Ichthyornis* ((Torres et al., 2021); (Benito et al., 2022); (Field et al., 2024); (Benito et al., 2026)). As such, the palatal morphology of these non-neornithine ornithurines would seem to provide tentative corroboration of Jollie’s (1957) hypothesis that the hemipterygoid of phasianoid Galliformes fuses with the caudal end of the palatine early in embryonic ontogeny. In this scenario, both stem ornithurines and phasianoid galliforms would be the only avian groups exhibiting an exclusively intrapterygoid palatal joint, without the participation of the palatine, supporting other lines of evidence indicating that the galliform skull is generally plesiomorphic in nature (e.g., Field et al. (2020); Benito et al. (2022); Kuo et al. (2023); Field et al. (2024); Crane et al. (2026); Arnaout et al. (2026)). Further research into the pre-hatching ontogeny of Galliformes will be necessary to test this hypothesis, and, at present, the interpretation that non-megapodiid galliforms simply lack a hemipterygoid entirely, with their caudal palatine converging on a similar morphology to the ichthyornithine hemipterygoid, cannot be refuted.

As discussed above, the presence of a hemipterygoid in extant palaeognaths is highly uncertain, with little evidence to suggest that the pointed rostral projections of certain palaeognath pterygoids (e.g., *Struthio*) are homologous with the hemipterygoid processes of Galloanserae or the hemipterygoids of neoavians or crownward stem birds. Furthermore, the extremely divergent contacts between the pterygoid and the palatine in various palaeognaths raises doubts as to the accuracy and conceptual utility of the oft-touted ‘palaeognath condition’ supposedly characterising the palates of all extant members of the clade. Fossil evidence is lacking to clarify the origin of the palaeognath palate from an ancestral condition presumably more similar to that of extant neognaths ((Torres et al., 2021); (Benito et al., 2022); (Field et al., 2025); (Benito et al., 2026) – but see also Wilken et al. (2025); Wilken et al. (2026)) – precluding the assessment of whether any form of pterygoid segmentation existed in the earliest palaeognaths. Indeed, known stem palaeognaths (representatives of the lithornithid assemblage) appear to exhibit fused palatine-pterygoid complexes similar to those of extant Struthioniformes and Casuariiformes (see Houde (1988)), yet no detailed study of the palatal anatomy of these fossil birds has yet been undertaken.

## 5. CONCLUSIONS

Fossil evidence from crownward stem birds suggests that at least some aspects of galloanseran pterygoid, ‘hemipterygoid’, and palatine morphology probably reflect the plesiomorphic condition of the crown bird palate, yet it remains unclear whether the evolutionary origin of pterygoid segmentation constitutes a neornithine, neognath or neoavian synapomorphy. Regardless, the present study illustrates that the full process of pterygoid segmentation is restricted to Neoaves. We hypothesise that the rostral projections of the pterygoid observable in Anseriformes and Megapodiidae represent ‘hemipterygoid processes’, homologous with the hemipterygoids of neoavians, though they do not detach from the pterygoid body during ontogeny. The presence of discrete hemipterygoids in stem group birds that remain unfused with the palatine suggest a decoupling of the origin of a discrete hemipterygoid and the origin of pterygoid segmentation. The recognition of a neognath-like mobile palate articulation in some stem birds (e.g., see Benito et al. (2022)) would suggest that neognath-like prokinesis arose prior to the origin of pterygoid segmentation.

Recent work illustrates that the arrangement of the neoavian palate is subject to important constraints in which development is channelled along specific morphological axes (Plateau et al., 2026), though the extent to which segmentation alone dictates this constrained pattern of development remains uncertain. The interplay between the onset of pterygoid segmentation and the expansion of avian developmental modes in Neoaves ((Starck & Ricklefs, 1998); (Ducatez & Field, 2021)) may underpin the diversity of ontogenetic trajectories in palate morphology in this clade, with differences in the degree of ossification and a chick’s size at hatching affecting the timing of pterygoid segmentation (e.g., Starck (1993); Starck and Ricklefs (1998); Botelho et al. (2015)). Further insight into the evolutionary history of pterygoid segmentation will necessitate new observations from embryonic development and the fossil record.

## Supporting information

Revised supplementary information Hunt et al

## Author contributions

AKH, DJF and JB conceived and designed the project, AU contributed materials, AKH performed digital segmentation of the bones (excluding *Presbyornis* and *Nettapterornis* which was by JB), AKH and JB designed figures, AKH wrote the original manuscript draft with input from DJF, OP and JB, editing and preparation of the final version of the manuscript was performed by AKH, DJF, JB and OP.

## Acknowledgements

We thank K. Smithson (Cambridge Biotomography Centre) for access to micro-CT scanning facilities. We thank L. Fisher (University of Cambridge) for early assistance with segmentation and interpretation. We thank Helen James and Holly Little (National Museum of Natural History, Smithsonian Institution) for access to CT-data of *Presbyornis* (USNM PAL 299846). This work was funded by UKRI grant MR/X015130/1 (D.J.F) and the SNSF grant P500PN_214284 (O.P). For the purpose of open access, the authors have applied a Creative Commons Attribution (CC BY) licence to any Author Accepted Manuscript version arising.

## Data availability statement

The CT data and 3D models generated during this study will be deposited on MorphoSource upon acceptance of the manuscript [doi to be added upon acceptance].

## Ethics approval statement

n/a

## Permission to reproduce material from other sources

n/a

