## Supplementary material for "Developmental variation in pterygoid segmentation clarifies patterns of avian bony palate evolution": Revised supplementary information Hunt et al

### SUPPLEMENTARY FIGURES AND TABLES

#### Supplementary figure legends:

**Supplementary figure 1:**

Comparative morphology of the palatine in select immature palaeognaths, oriented for the right side of the skull (*Struthio*, *Apteryx* and *Megalapteryx* elements are reflected). Abbreviations: clf, caudolateral flange of pterygoid process of palatine; D, dorsal; L, lateral; pp, pterygoid process of palatine; pp(dl) dorsolateral branch of pterygoid process of palatine; pp(vm) ventromedial branch of pterygoid process of palatine; R, rostral; rmp, rostromedial process of palatine; rp, rostral process of palatine. Images oriented with respect to the skull. All scale bars represent 2.5 mm.

**Supplementary figure 2:**

Comparative morphology of the pterygoid in select immature palaeognaths, oriented for the right side of the skull (*Struthio*, *Apteryx* and *Megalapteryx* elements are reflected). To optimise informativeness of individual images, here orientations for dorsal, medial, ventral, lateral, rostral and caudal (and associated directional arrows) do not necessarily correspond exactly to these orientations with respect to the skull itself. Abbreviations: bpc, basipterygoid process contact surface; D, dorsal; L, lateral; pc, palatine contact; qas, quadrate articular surface; R, rostral; rpt, rostral projection of pterygoid. Scale bars represent 0.5 mm for rostral and caudal views in all taxa apart from the ostrich and moa, which equal 2.5 mm. All other scale bars represent 2.5 mm.

**Supplementary figure 3:**

Comparative morphology of the palatine in select immature anseriforms, oriented for the right side of the skull (*Chauna*, *Anser*, *Aythya*, *Malacorhynchus*, *Spatula* and *Anas* elements are reflected). Abbreviations: cc, caudolateral corner of palatine; D, dorsal; L, lateral; pp, pterygoid process of palatine; R, rostral; rp, rostral process of palatine; rmp, rostromedial process of palatine. Images oriented with respect to the skull. All scale bars represent 2.5 mm.

**Supplementary figure 4:**

Comparative morphology of the pterygoid in select immature anseriforms (part 1), oriented for the right side of the skull (*Chauna* and *Anser* elements are reflected). To optimise informativeness of individual images, here orientations for dorsal, medial, ventral, lateral, rostral and caudal (and associated directional arrows) do not necessarily correspond exactly to these orientations with respect to the skull itself. Abbreviations: bpc, basipterygoid process contact surface; D, dorsal; L, lateral; pc, palatine contact surface; qas, quadrate articular surface; R, rostral; rpt, rostral projection of pterygoid. Scale bars by rostral view images represent 0.5 mm for all taxa and apply to rostral and caudal views. All other scale bars represent 2.5 mm.

**Supplementary figure 5:**

Comparative morphology of the pterygoid in select immature anseriforms (part 2), oriented for the right side of the skull (*Aythya*, *Malacorhynchus*, *Spatula* and *Anas* elements are reflected). To optimise informativeness of individual images, here orientations for dorsal, medial, ventral, lateral, rostral and caudal (and associated directional arrows) do not necessarily correspond exactly to these orientations with respect to the skull itself. Abbreviations: bpc, basipterygoid process contact surface;

D, dorsal; L, lateral; pc, palatine contact surface; qas, quadrate articular surface; R, rostral; rpt, rostral projection of pterygoid. Scale bars by rostral view images represent 0.5 mm for all taxa and apply to rostral and caudal views. All other scale bars represent 2.5 mm.

**Supplementary figure 6:**

Comparative morphology of the palatine in select immature galliforms, oriented for the right side of the skull (*Megapodius*, *Numida*, *Rollulus*, *Coturnix* and *Gallus* (domestic) elements are reflected). Abbreviations: D, dorsal; L, lateral; pp, pterygoid process of palatine; R, rostral; rp, rostral process of palatine; rmp, rostromedial process of palatine. Images oriented with respect to the skull. All scale bars represent 2.5 mm.

**Supplementary figure 7:**

Comparative morphology of the pterygoid in select immature galliforms, oriented for the right side (*Megapodius*, *Numida*, *Rollulus*, *Coturnix* and *Gallus* (domestic) elements are reflected). To optimise informativeness of individual images, here orientations for dorsal, medial, ventral, lateral, rostral and caudal (and associated directional arrows) do not necessarily correspond exactly to these orientations with respect to the skull itself. Abbreviations: D, dorsal; L, lateral; qas, quadrate articular surface; R, rostral. Scale bars represent 0.5 mm for rostral and caudal views in all taxa. All other scale bars represent 2.5 mm.

**Supplementary figure 8:**

Comparative morphology of the palatine in select immature neoavians (part 1), oriented for the right side of the skull (*Apus* and *Otidiphaps* elements are reflected). Cladogram on the left illustrates the phylogenetic relationships of the select neoavians (higher order relationships follow Prum et al. (2015); detailed interrelationships among subclades follow Cooney et al. (2017)). **A** – Strisores; **B** – Columbaves; **C** – Gruiformes. Numbers correspond to the following taxa: 1. *Apus apus;* 2. *Tauraco erythrolophus;* 3. *Musophaga violacea;* 4. *Otidiphaps nobilis;* 5. *Columba livia;* 6. *Gallirallus australis*. Abbreviations: D, dorsal; L, lateral; pp, pterygoid process of palatine; R, rostral; rp, rostral process of palatine; rmp, rostromedial process of palatine. Images oriented with respect to the skull. All scale bars represent 2.5 mm.

**Supplementary figure 9:**

Comparative morphology of the palatine in select immature neoavians (part 2) (Inopinaves), oriented for the right side of the skull (*Opisthocomus*, *Pteroglossus* and *Lorius* elements are reflected). Cladogram on the left illustrates the phylogenetic relationships of the select neoavians (higher order relationships follow Prum et al. (2015); detailed interrelationships among subclades follow Cooney et al. (2017)). Numbers correspond to the following taxa: 1. *Opisthocomus hoazin;* 2. *Vultur gryphus;* 3. *Pteroglossus viridis;* 4. *Lorius lory;* 5. *Ara macao.* Abbreviations: D, dorsal; L, lateral; pp, pterygoid process of palatine; R, rostral; rp, rostral process of palatine; rmp, rostromedial process of palatine. Images oriented with respect to the skull. All scale bars represent 2.5 mm.

**Supplementary figure 10**:

Comparative morphology of the pterygoid and hemipterygoid in select immature neoavians (part 1), oriented for the right side of the skull (*Apus* and *Otidiphaps* elements are reflected). Cladogram on the left illustrates the phylogenetic relationships of the select neoavians (higher order relationships follow Prum et al. (2015); detailed interrelationships among subclades follow Cooney et al. (2017)). **A** – Strisores; **B** – Columbaves; **C** – Gruiformes. Numbers correspond to the following taxa: 1. *Apus apus*, 2. *Tauraco erythrolophus*, 3. *Musophaga violacea,* 4. *Otidiphaps nobilis,* 5. *Columba livia,* 6. *Gallirallus australis.* To optimise informativeness of individual images, here orientations for dorsal, medial, ventral, lateral, rostral and caudal (and associated directional arrows) do not necessarily correspond exactly to these orientations with respect to the skull itself. Abbreviations: D, dorsal; hmp (ST.1), unsegmented hemipterygoid (Stage 1 of pterygoid segmentation); hmp (ST.2), partially segmented hemipterygoid (Stage 2 of pterygoid segmentation); hmp (ST.3), fully segmented hemipterygoid (Stage 3 of pterygoid segmentation); L, lateral; pc, palatine contact surface; qas, quadrate articular surface; R, rostral. Dashed lines to 'pc' indicate the contact is present on the side that is hidden from view. Scale bars by rostral view images represent 0.5 mm and apply to rostral and caudal views. All other scale bars represent 2.5 mm.

**Supplementary figure 11**:

Comparative morphology of the pterygoid and hemipterygoid in select immature neoavians (part 2) (Inopinaves), oriented for the right side of the skull (*Opisthocomus*, *Pteroglossus* and *Lorius* elements are reflected). Cladogram on the left illustrates the phylogenetic relationships of the select neoavians (higher order relationships follow Prum et al. (2015); detailed interrelationships among subclades follow Cooney et al. (2017)). Numbers correspond to the following taxa: 1. *Opisthocomus hoazin;* 2. *Vultur gryphus;* 3. *Pteroglossus viridis;* 4. *Lorius lory;* 5. *Ara macao.* To optimise informativeness of individual images, here orientations for dorsal, medial, ventral, lateral, rostral and caudal (and associated directional arrows) do not necessarily correspond exactly to these orientations with respect to the skull itself. Abbreviations: D, dorsal; hmp (ST.1), unsegmented hemipterygoid (Stage 1 of pterygoid segmentation); hmp (ST.2), partially segmented hemipterygoid (Stage 2 of pterygoid segmentation); L, lateral; pc, palatine contact surface; qas, quadrate articular surface; R, rostral. Dashed lines to 'pc' indicate the contact is present on the side that is hidden from view. Scale bars by rostral view images represent 0.5 mm and apply to rostral and caudal views. All other scale bars represent 2.5 mm.

**Supplementary figure 12**:

Comparative morphology of the pterygoid and hemipterygoid in select immature neoavians (part 3) (Aequorlitornithes), oriented for the right side of the skull (*Phoenicopterus*, *Calidris*, *Spheniscus* and *Macronectes* elements are reflected). Cladogram on the left illustrates the phylogenetic relationships of the select neoavians (higher order relationships follow Prum et al. (2015); detailed interrelationships among subclades follow Cooney et al. (2017)). Numbers correspond to the following taxa: 1. *Phoenicopterus ruber,* 2. *Calidris pugnax,* 3. *Recurvirostra avosetta,* 4. *Spheniscus demersus,* 5. *Diomedea exulans,* 6. *Macronectes giganteus.* To optimise informativeness of individual images, here orientations for dorsal, medial, ventral, lateral, rostral and caudal (and associated directional arrows) do not necessarily correspond exactly to these orientations with respect to the skull itself. Abbreviations: D, dorsal; hmp (ST.2), partially segmented hemipterygoid (Stage 2 of pterygoid segmentation); hmp (ST.4), discrete hemipterygoid with partial fusion to palatine (Stage 4 of pterygoid segmentation); L, lateral; pc, palatine contact surface; qas, quadrate articular surface; R, rostral. Dashed lines to ‘pc’ indicate the contact is present on the side that is hidden from view. Scale bars by rostral view images represent 0.5 mm and apply to rostral and caudal views. All other scale bars represent 2.5 mm.

**Supplementary figure 13**:

Comparative morphology of the pterygoid and hemipterygoid in select immature neoavians (part 4) (Aequorlitornithes), oriented for the right side of the skull. Cladogram on the left illustrates the phylogenetic relationships of the select neoavians (higher order relationships follow Prum et al. (2015); detailed interrelationships among subclades follow Cooney et al. (2017)). Numbers correspond to the following taxa: 1. *Fulmarus glacialis,* 2. *Morus bassanus,* 3. *Egretta garzetta,* 4. *Nycticorax nycticorax,* 5. *Ardea cinerea.* To optimise informativeness of individual images, here orientations for dorsal, medial, ventral, lateral, rostral and caudal (and associated directional arrows) do not necessarily correspond exactly to these orientations with respect to the skull itself. Abbreviations: D, dorsal; hmp (ST.2), partially segmented hemipterygoid (Stage 2 of pterygoid segmentation); hmp (ST.4), discrete hemipterygoid with partial fusion to palatine (Stage 4 of pterygoid segmentation); L, lateral; pc, palatine contact surface; pc (pf), partial fusion of hemipterygoid with palatine at contact surface; qas, quadrate articular surface; R, rostral. Dashed lines to ‘pc’ indicate the contact is present on the side that is hidden from view. Scale bars by rostral view images represent 0.5 mm and apply to rostral and caudal views (apart from *Nycticorax* which represents 2.5 mm). All other scale bars represent 2.5 mm.

**Supplementary figure 14:**

Comparative morphology of the palatine in select immature neoavians (Aequorlitornithes) (part 3), oriented for the right side of the skull (*Phoenicopterus*, *Calidris*, *Spheniscus* and *Macronectes* elements are reflected). Cladogram on the left illustrates the phylogenetic relationships of the select neoavians (higher order relationships follow Prum et al. (2015); detailed interrelationships among subclades follow Cooney et al. (2017)). Numbers correspond to the following taxa: 1, *Phoenicopterus ruber;* 2. *Calidris pugnax;* 3. *Recurvirostra avosetta;* 4. *Spheniscus demersus;* 5. *Diomedea exulans;* 6. *Macronectes giganteus;* 7. *Fulmarus glacialis;* 8*. Morus bassanus;* 9. *Egretta garzetta;* 10*. Nycticorax nycticorax;* 11. *Ardea cinerea.* Abbreviations: D, dorsal; L, lateral; pp, pterygoid process of palatine; R, rostral; rp, rostral process of palatine; rmp, rostromedial process of palatine. Images oriented with respect to the skull. All scale bars represent 2.5 mm.

**Supplementary figure 1:**

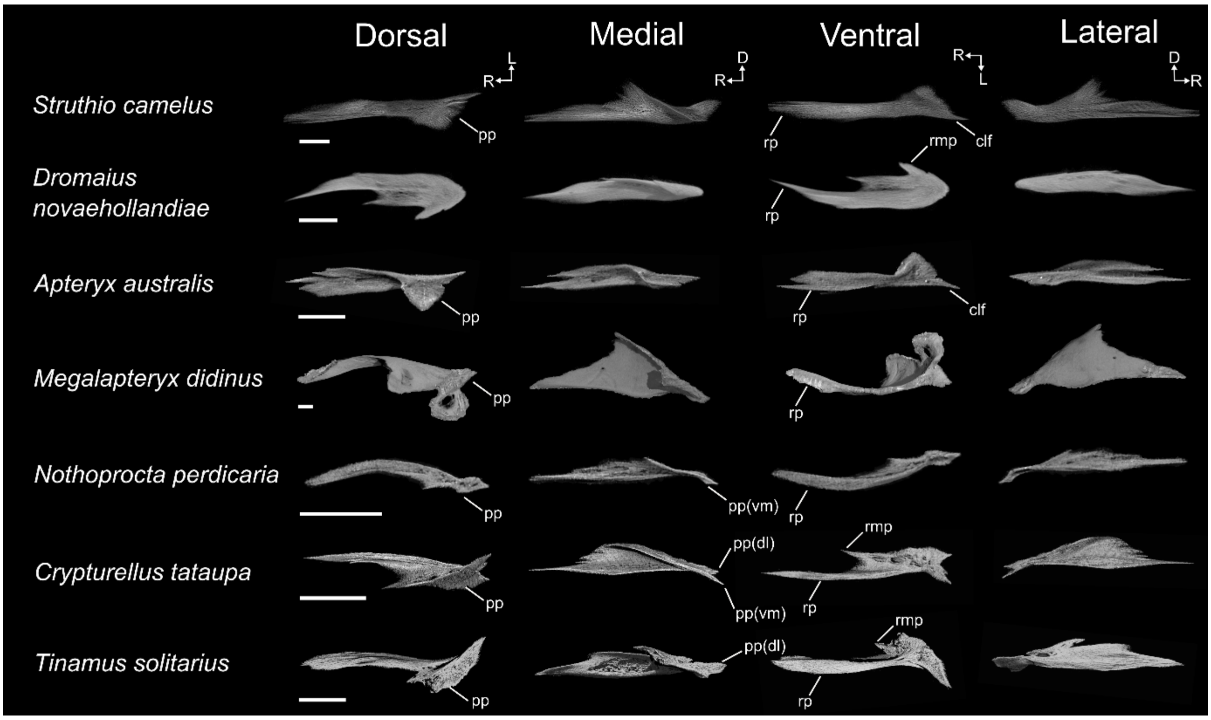

**Supplementary figure 2:**

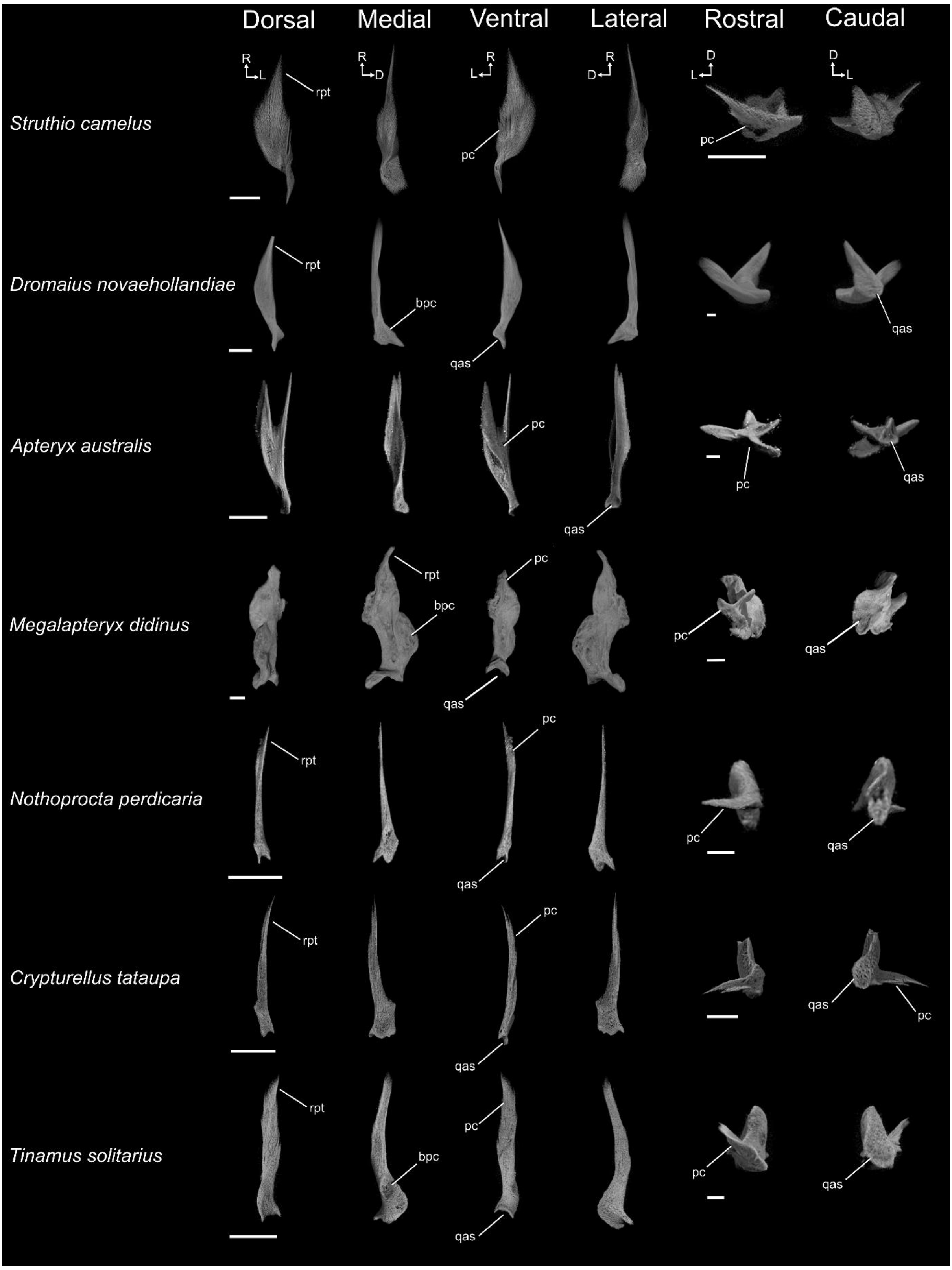

#### Supplementary figure 3:

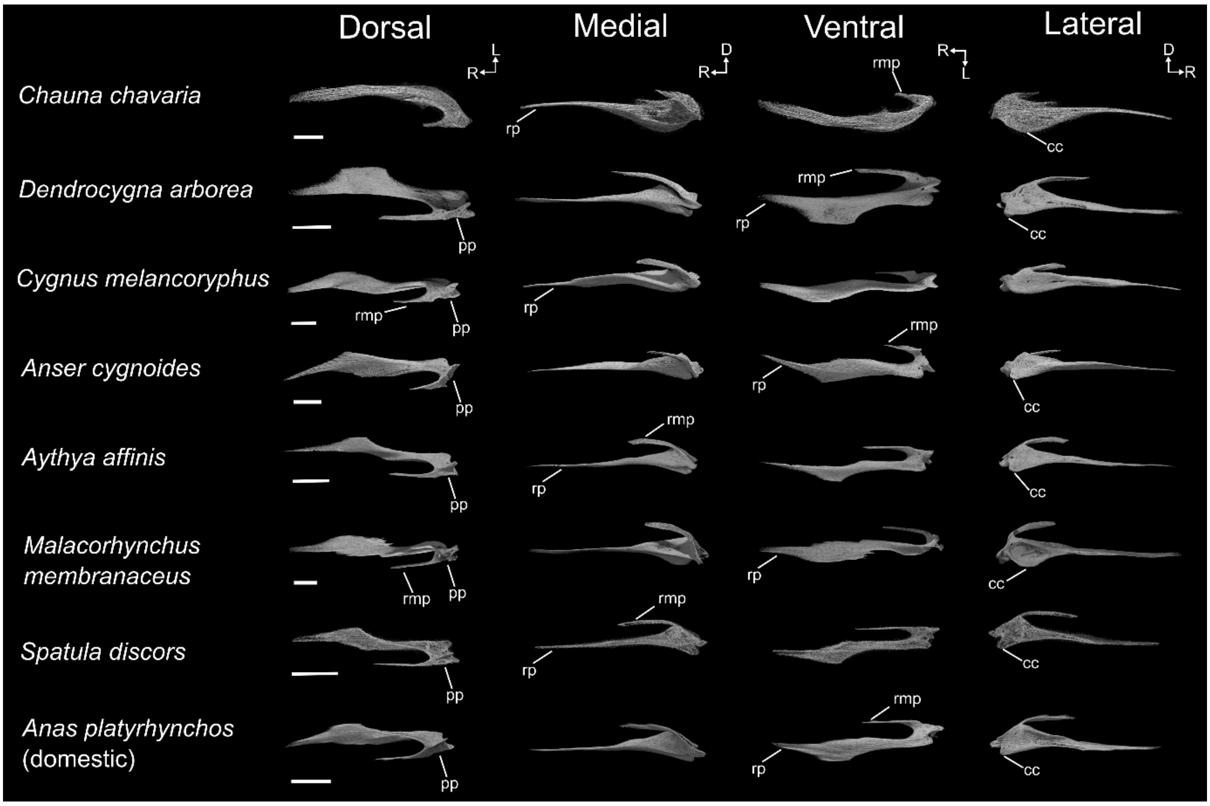

#### Supplementary figure 4:

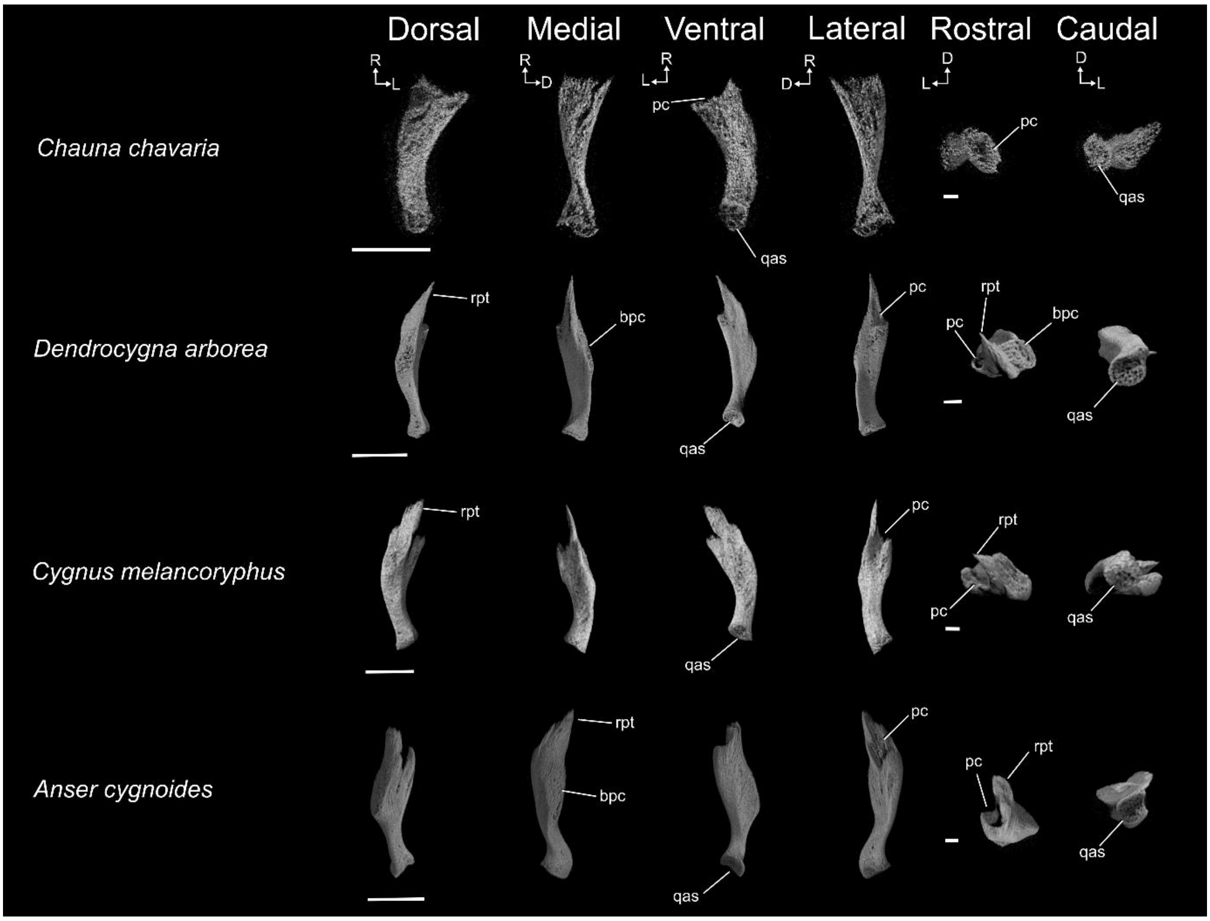

#### Supplementary figure 5:

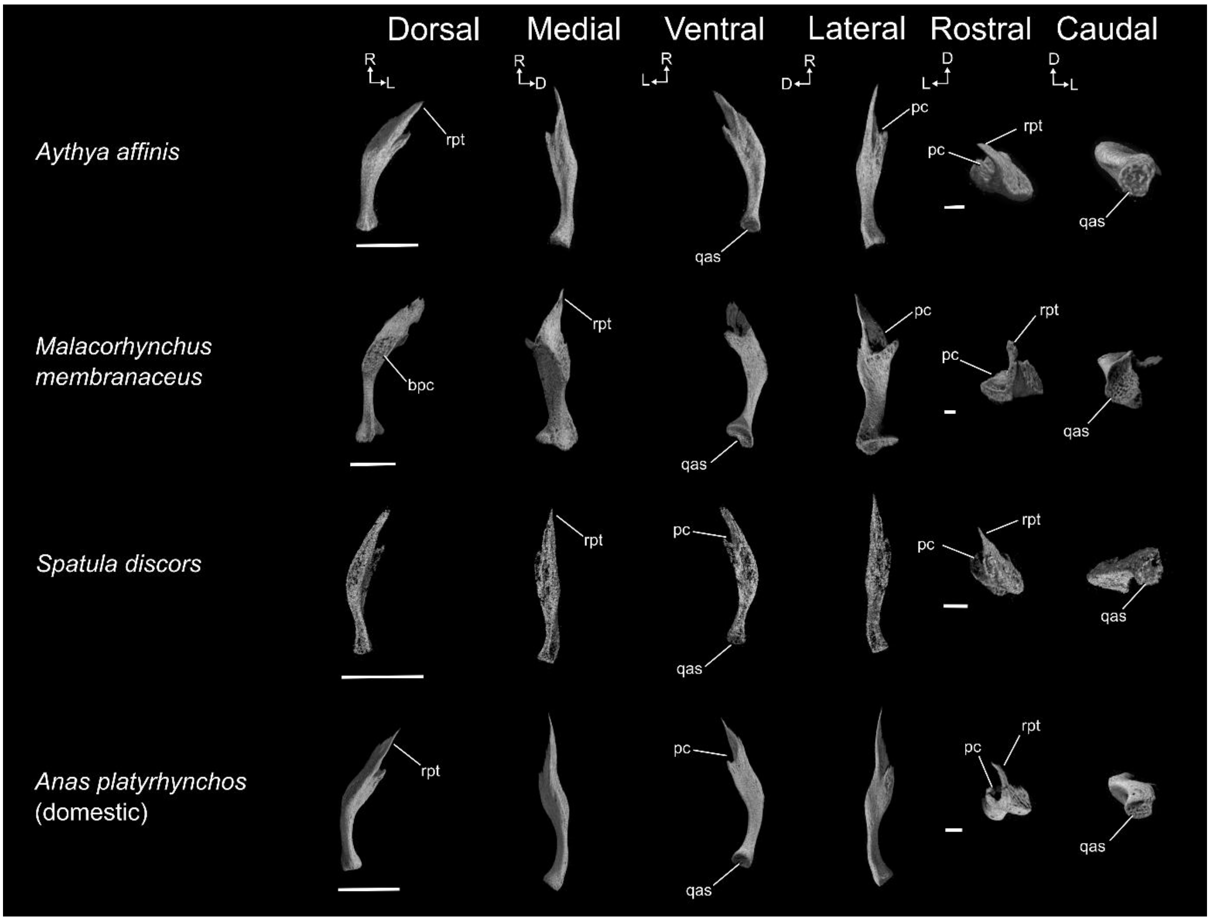

#### Supplementary figure 6:

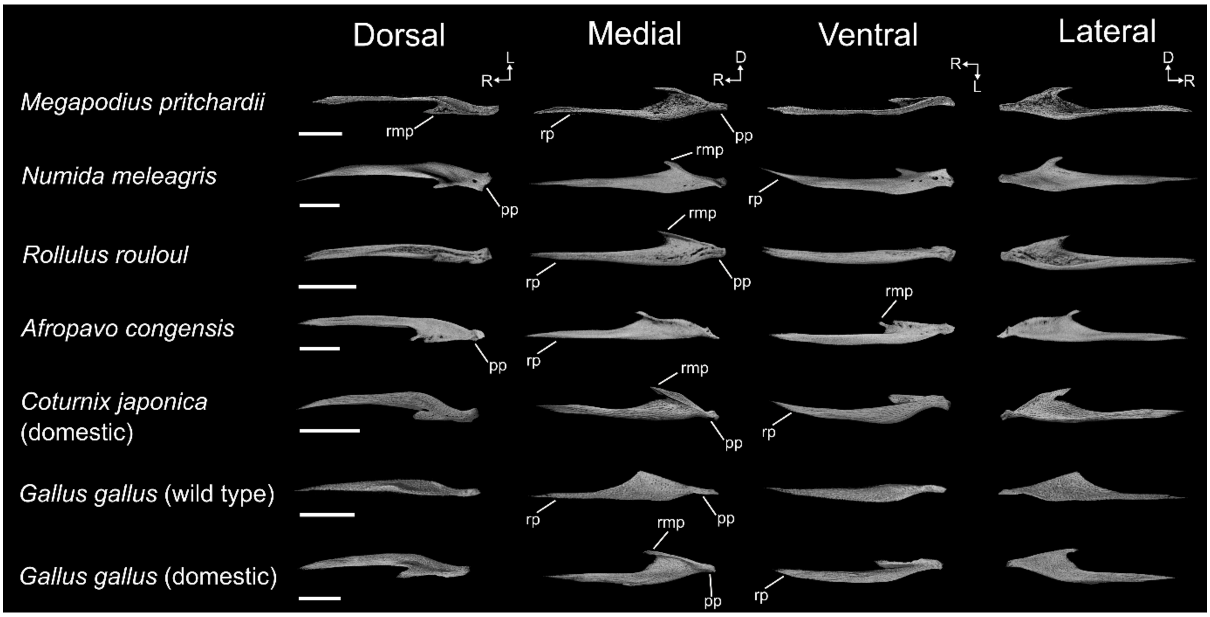

**Supplementary figure 7:**

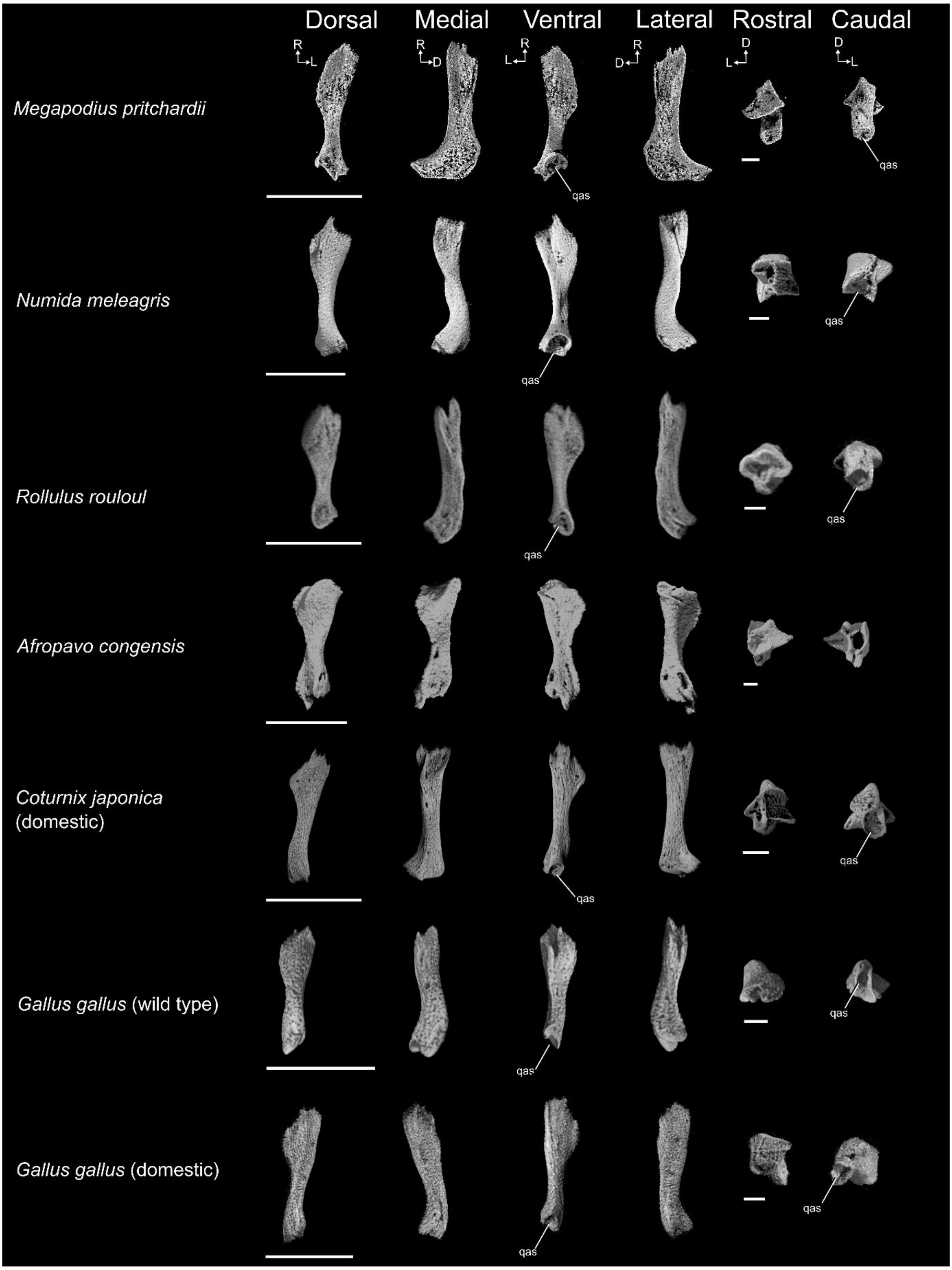

#### Supplementary figure 8:

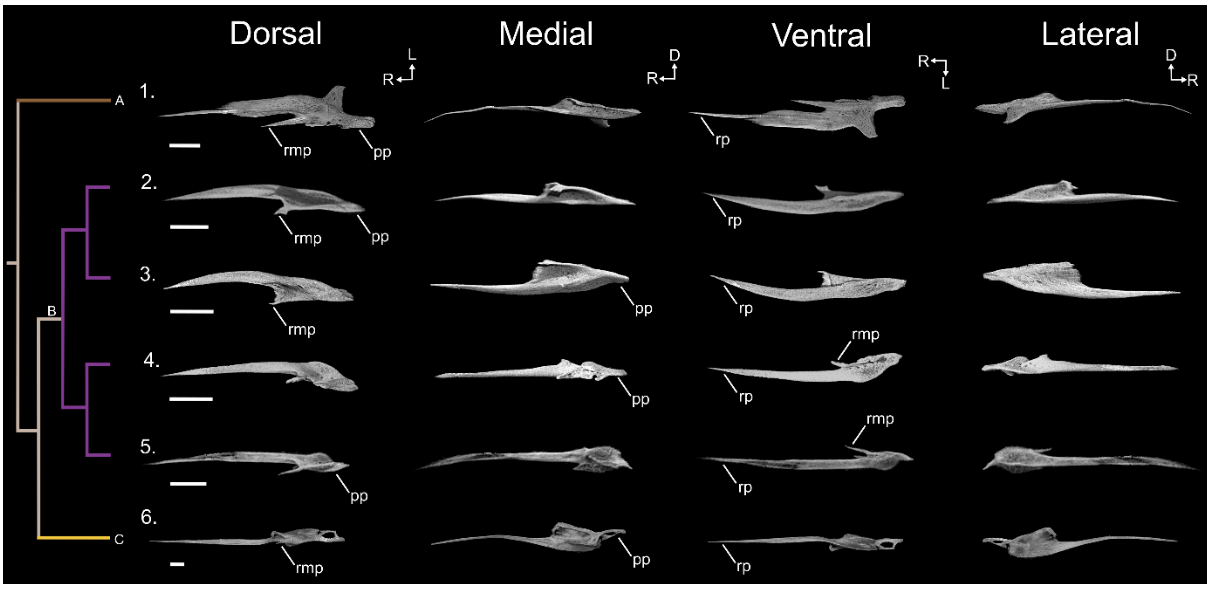


#### Supplementary figure 9:

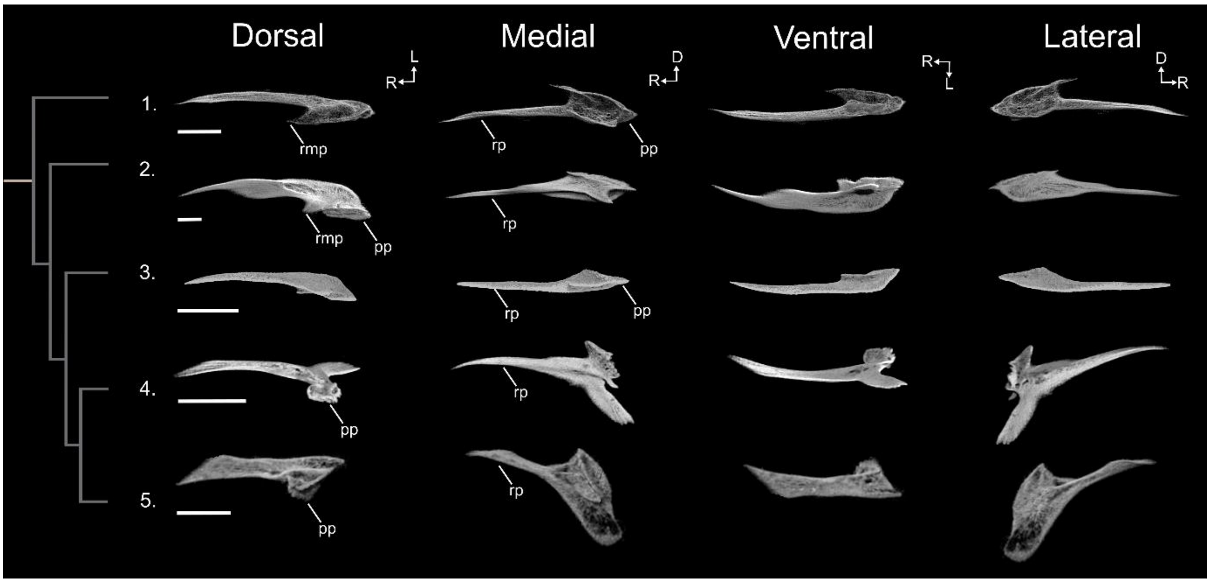

#### Supplementary figure 10:

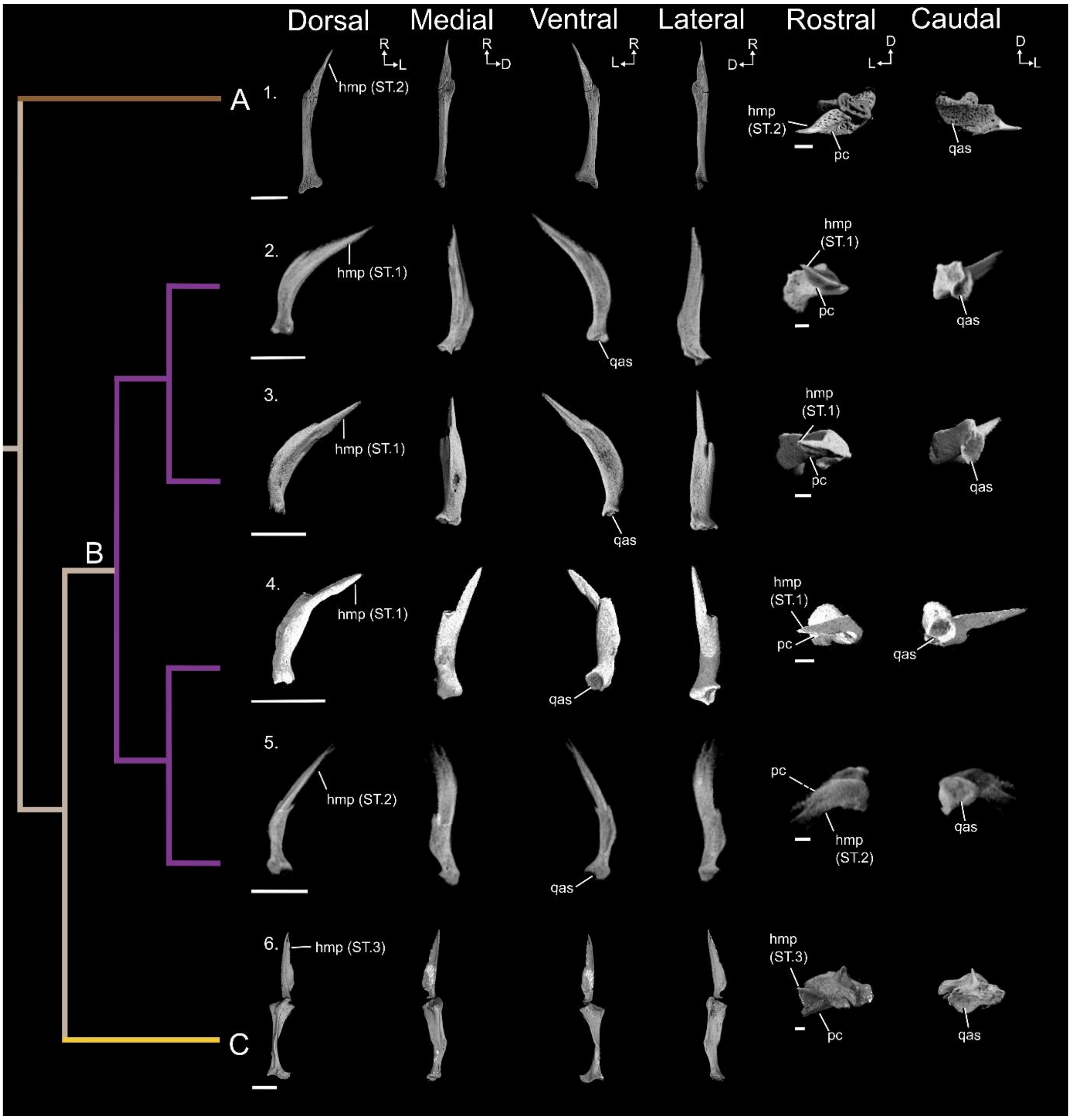

#### Supplementary figure 11:

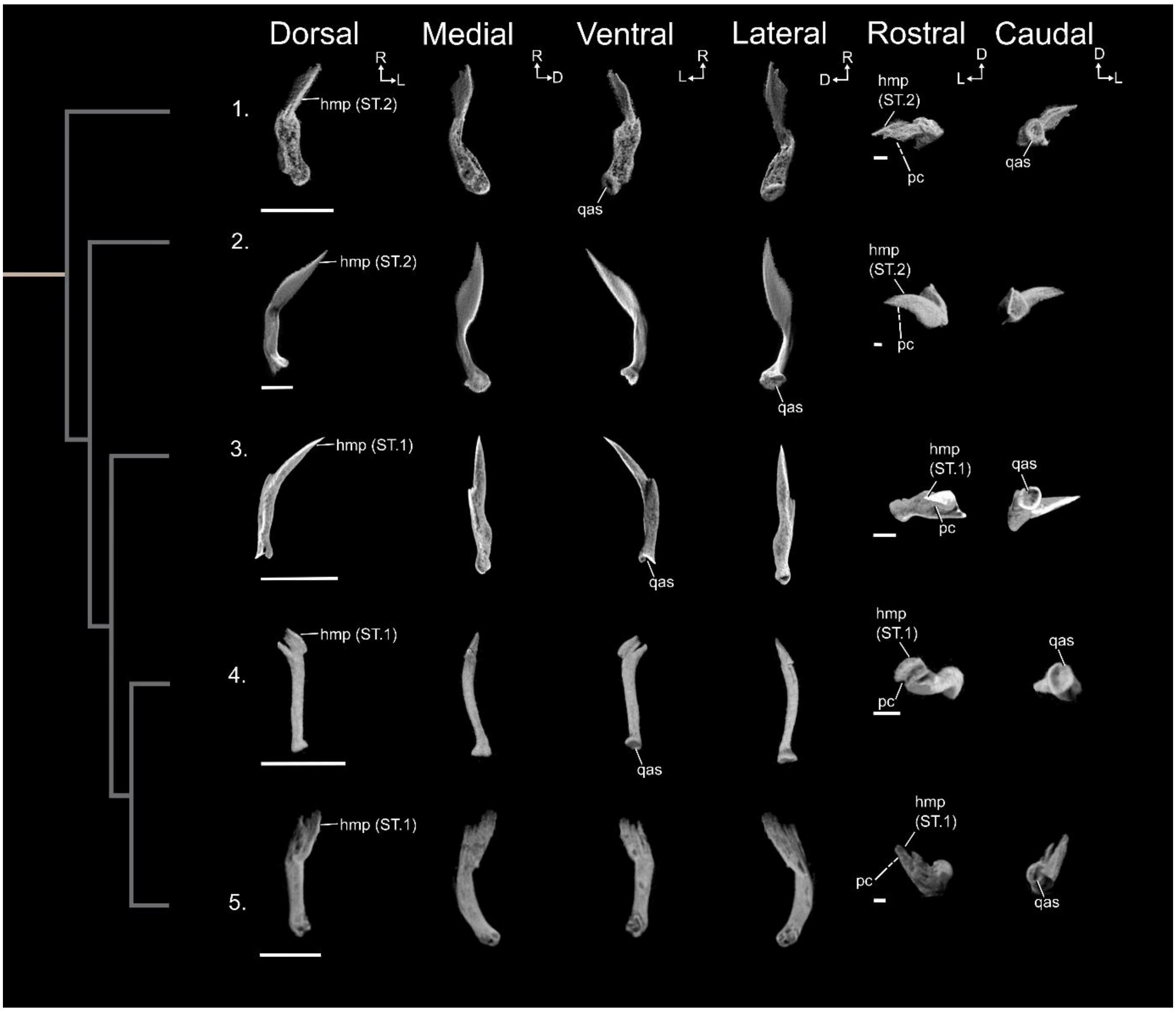

#### Supplementary figure 12:

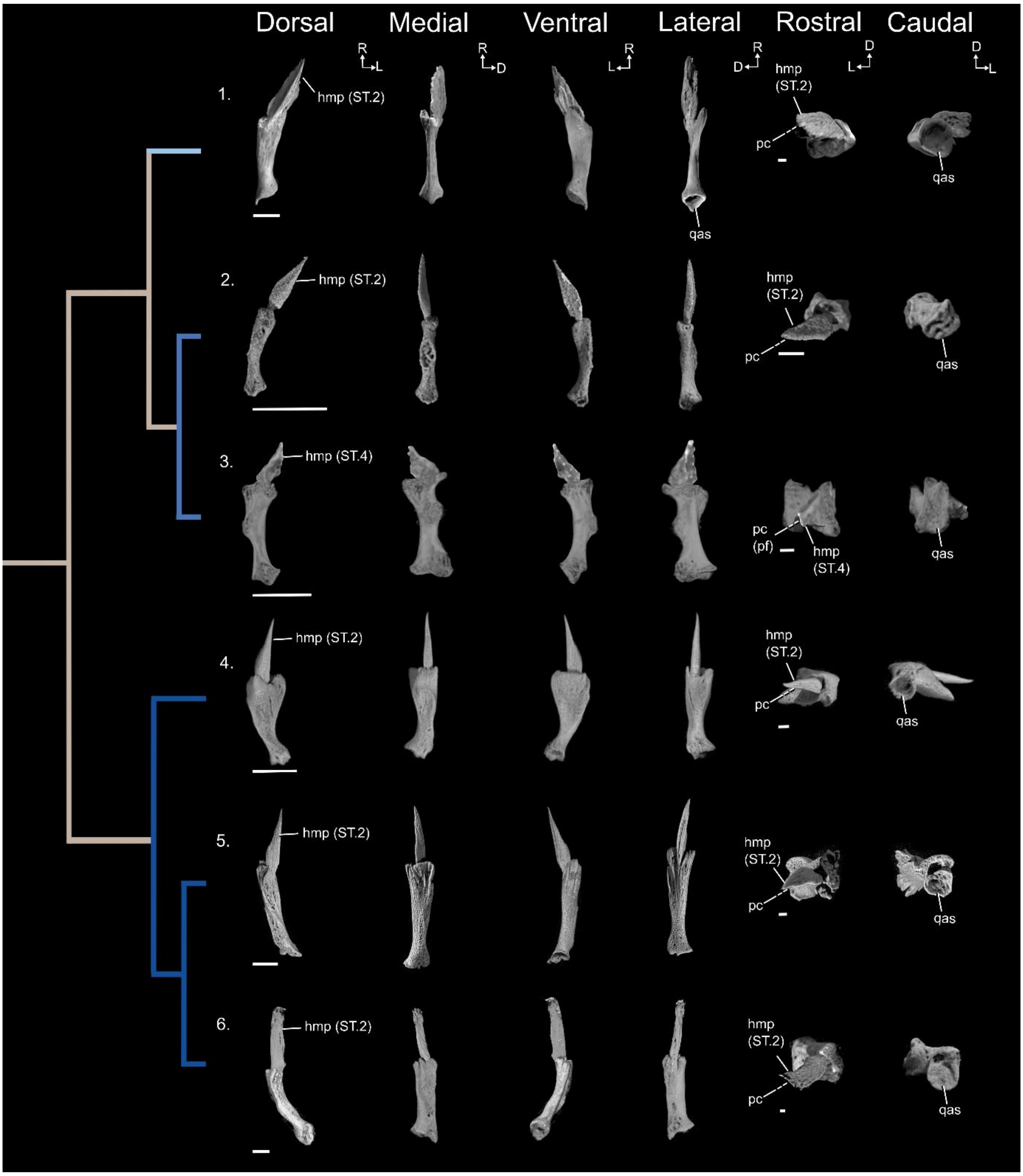

#### Supplementary figure 13:

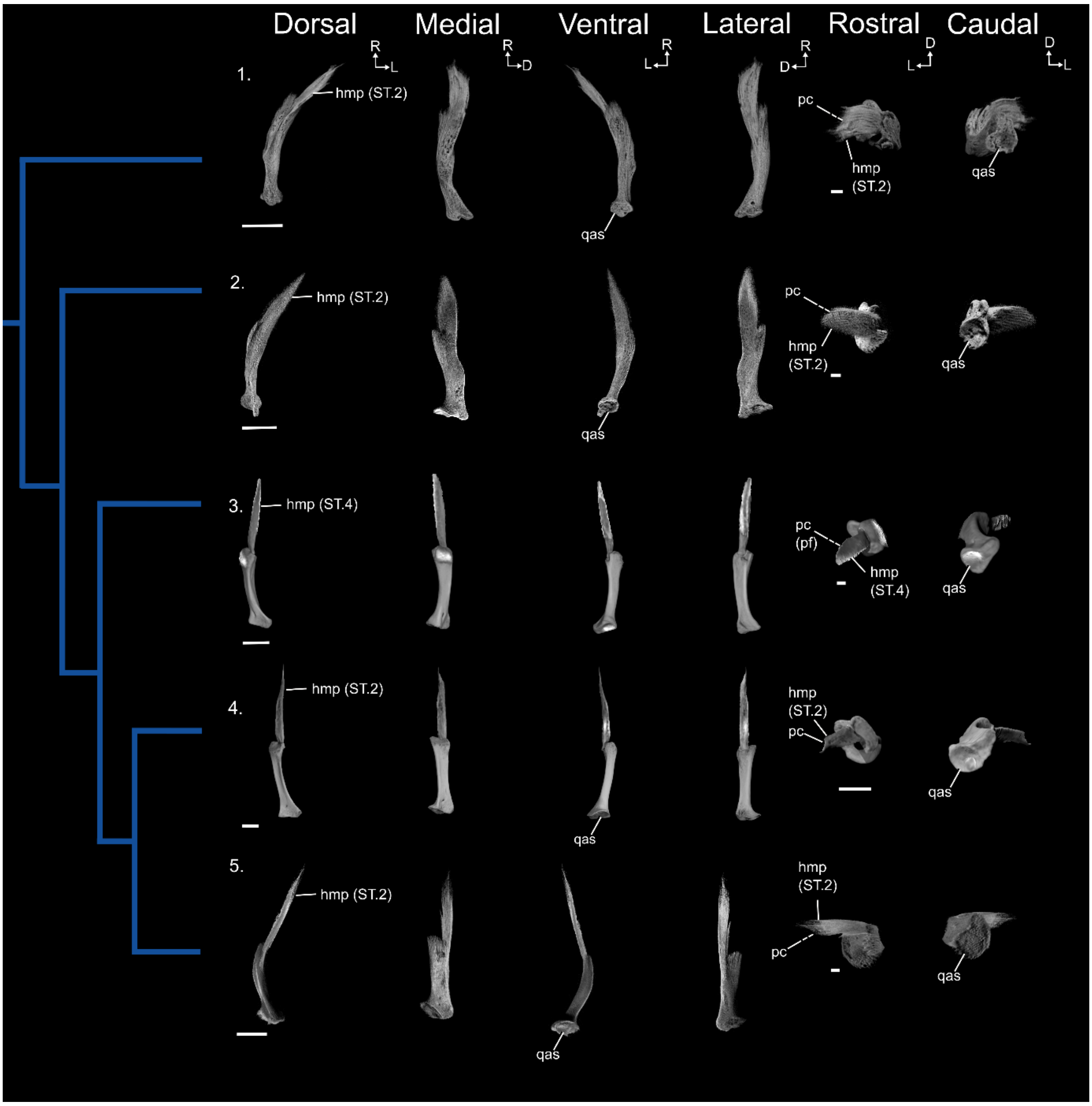

#### Supplementary figure 14:

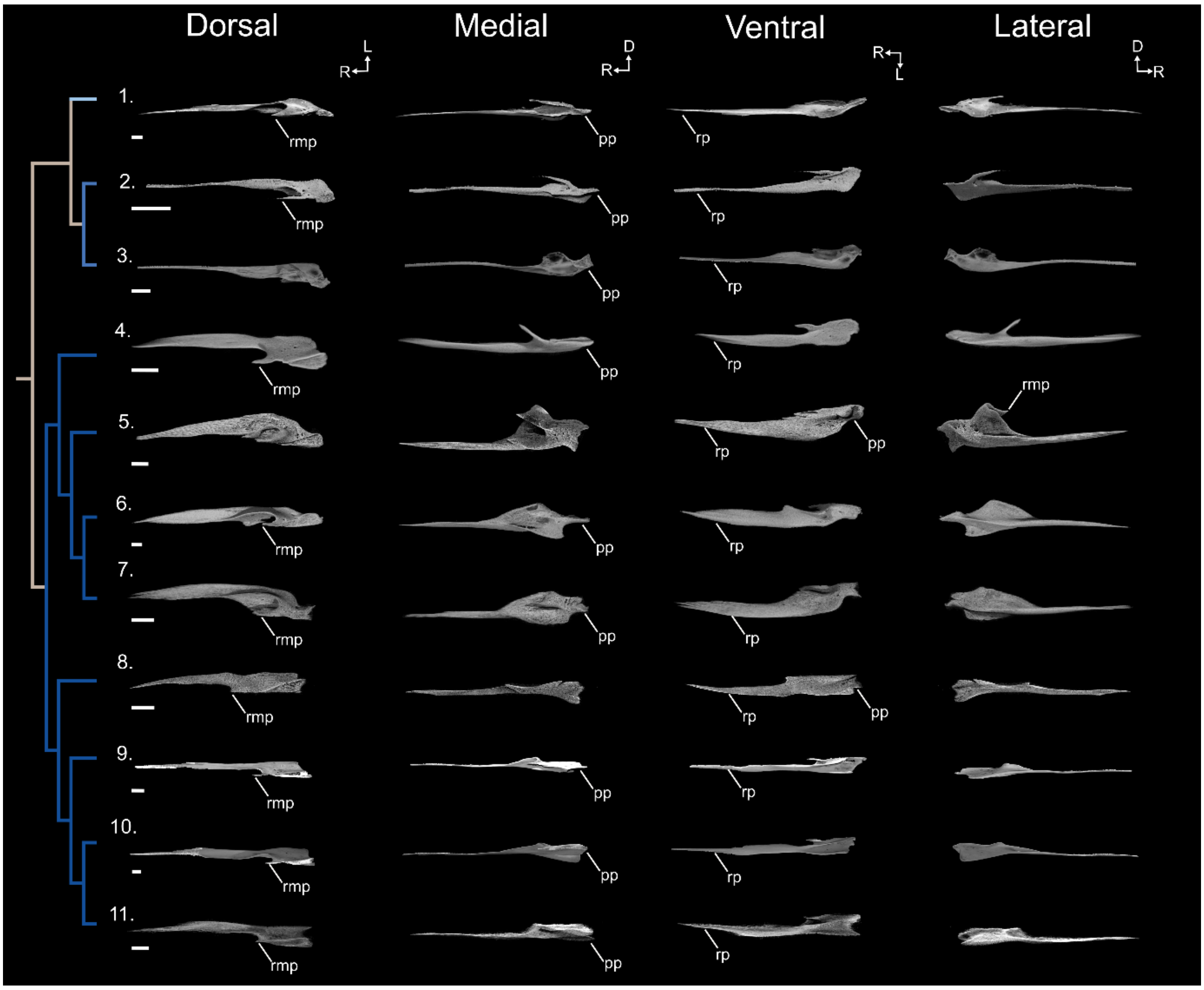


#### Supplementary table legends:

**Supplementary table 1**: List of the taxa digitally segmented and figured in this study. Abbreviations: BMNH, British Museum of Natural History, London (now NHM); FMNH, Field Museum of Natural History, Chicago; NMBE, Naturhistorisches Museum Bern; NHM, Natural History Museum, London; MNHF, Musée d'Histoire Naturelle Fribourg; UMZC, University Museum of Zoology, Cambridge; USNM, National Museum of Natural History, Washington, DC.

| ORDER | TAXON | COMMON NAME | COLLECTION NUMBER | TYPE | ONTOGENETIC STAGE | MORPHOSOURCE ID |
|---|---|---|---|---|---|---|
| Tinamiformes | *Tinamus solitarius* | Solitary Tinamou | UMZC 2022.11.49 | Wet | Embryo | |
| Tinamiformes | *Crypturellus tataupa* | Tataupa Tinamou | UMZC 2022.11.35 | Wet | Embryo | |
| Tinamiformes | *Crypturellus tataupa* | Tataupa Tinamou | UMZC uncatalogued | Osteo | Adult | |
| Tinamiformes | *Nothoprocta perdicaria* | Chilean Tinamou | UMZC uncatalogued | Wet | Embryo | |
| Casuariiformes | *Dromaius novaehollandiae* | Emu | UMZC uncatalogued | Wet | Embryo | |
| Casuariiformes | *Dromaius novaehollandiae* | Emu | UMZC 362 | Osteo | Subadult | |
| Apterygiformes | *Apteryx australis* | Southern Brown Kiwi | UMZC 5/Apt/1/a/9 | Wet | Hatchling | |
| Apterygiformes | *Apteryx australis* | Southern Brown Kiwi | FMNH 391011 | Osteo | Subadult | ID: 000108118 |
| Struthioniformes | *Struthio camelus* | Common Ostrich | UMZC uncatalogued | Wet | Embryo | |
| Struthioniformes | *Struthio camelus* | Common Ostrich | UMZC uncatalogued | Osteo | Subadult | |
| Dinornithiformes | *Megalapteryx didinus* | Upland Moa | NHM A16 | Desiccated | Subadult | |
| Galliformes | *Gallus gallus* | Red junglefowl | UMZC 14/Pha/24/a/30 | Wet | Embryo | |
| Galliformes | *Gallus gallus* (domestic) | Domesticated Red Junglefowl | UMZC 400 | Osteo | Adult | |
| Galliformes | *Gallus gallus* (domestic) | Domesticated Red Junglefowl | UMZC uncatalogued | Wet | Embryo | |
| Galliformes | *Rollulus rouloul* | Crested Partridge | UMZC 2022.11.40 | Wet | Hatchling | |
| Galliformes | *Afropavo congensis* | Congo Peacock | UMZC 2022.11.33 | Wet | Hatchling | |
| Galliformes | *Coturnix japonica* (domestic) | Domesticated Japanese Quail | UMZC uncatalogued | Wet | Hatchling | |
| Galliformes | *Megapodius pritchardii* | Tongan Megapode | UMZC 14/Meg/6/g/2 | Wet | Hatchling | |
| Galliformes | *Alectura lathami* | Australian Brushturkey | NHMUK S-2010.1.31 | Osteo | Adult | |
| Galliformes | *Leipoa ocellata* | Malleefowl | UMZC 391.A | Osteo | Adult | |
| Galliformes | *Megapodius nicobariensis* | Nicobar Megapode | UMZC 14/Meg/6/f/3 | Skin | Adult | |
| Galliformes | *Numida meleagris* | Helmeted Guineafowl | UMZC 393 | Osteo | Adult | |
| Galliformes | *Numida meleagris* | Helmeted Guineafowl | UMZC uncatalogued | Wet | Hatchling | |
| Anseriformes | *Chauna chavaria* | Northern Screamer | UMZC 211.A | Osteo | Adult | |
| Anseriformes | *Chauna chavaria* | Northern Screamer | BMNH 1923.5.24.30 | Wet | Hatchling | |
| Anseriformes | *Anhima cornuta* | Horned Screamer | NHMUK S/2009.2.2 | Osteo | Adult | |
| Anseriformes | *Anas platyrhynchos domesticus* | Domesticated Duck | UMZC uncatalogued | Wet | Embryo | |
| Anseriformes | *Malacorhynchus membranaceus* | Pink-eared Duck | UMZC uncatalogued | Frozen | Juvenile | |
| Anseriformes | *Anser cygnoides* | Swan Goose | UMZC uncatalogued | Wet | Hatchling | |
| Anseriformes | *Aythya ferina* | Common Pochard | NHMUK:1851.12.23.15 | Osteo | Adult | ID: 000125870 |
| Anseriformes | *Aythya affinis* | Lesser Scaup | UMZC uncatalogued | Wet | Hatchling | |
| Anseriformes | *Cygnus melancoryphus* | Black-necked Swan | UMZC uncatalogued | Wet | Hatchling | |
| Anseriformes | *Dendrocygna arborea* | West Indian Whistling Duck | UMZC uncatalogued | Wet | Hatchling | |
| Anseriformes | *Spatula discors* | Blue-winged Teal | UMZC uncatalogued | Wet | Hatchling | |
| Total-group Anseriformes | *Presbyornis pervetus* | NA | USNM PAL 299846 | Fossil | Adult | |
| Total-group Anseriformes | *Nettapterornis oxfordi* | NA | NHMUK PVA 5922 | Fossil | Adult | |
| Caprimulgiformes | *Apus apus* | Common Swift | BMNH 1893.1.24.3 | Skin | Hatchling | |
| Caprimulgiformes | *Apus apus* | Common Swift | NMBE 189/83 | Osteo | Adult | |
| Musophagiformes | *Tauraco erythrolophus* | Red-crested Turaco | UMZC uncatalogued | Frozen | Adult | |
| Musophagiformes | *Tauraco erythrolophus* | Red-crested Turaco | UMZC 2022.11.8 | Wet | Embryo | |
| Musophagiformes | *Musophaga violacea* | Violet Turaco | UMZC 2022.11.38 | Wet | Hatchling | |
| Columbiformes | *Otidiphaps nobilis* | Pheasant Pigeon | UMZC 2022.11.34 | Wet | Embryo | |
| Columbiformes | *Columba livia* (domestic) | Domestic Rock Pigeon | UMZC 721 | Osteo | Adult | |
| Columbiformes | *Columba livia* | Rock Pigeon | UMZC uncatalogued | Frozen | Hatchling | |
| Gruiformes | *Gallirallus australis* | Weka | UMZC 15/Ral/20/a/10 | Skin | Juvenile | |
| Gruiformes | *Gallirallus australis* | Weka | BMNH S/1955.5.16 | Osteo | Adult | |
| Phoenicopteriformes | *Phoenicopterus ruber* | American Flamingo | NMBE 178/2/72 | Osteo | Juvenile | |
| Phoenicopteriformes | *Phoenicopterus ruber* | American Flamingo | NMBE 297/82 | Osteo | Adult | |
| Charadriiformes | *Calidris pugnax* | Ruff | UMZC 2022.11.43 | Wet | Hatchling | |
| Charadriiformes | *Recurvirostra avosetta* | Pied Avocet | UMZC 16/Rec/4/c/5 | Skin | Juvenile | |
| Charadriiformes | *Recurvirostra avosetta* | Pied Avocet | NHMUK 1962.10.1 | Osteo | Adult | ID: 000092729 |
| Sphenisciformes | *Spheniscus demersus* | African Penguin | UMZC 2022.11.1 | Wet | Hatchling | |
| Sphenisciformes | *Spheniscus demersus* | African Penguin | BMNH S/2010.1.72 | Osteo | Adult | |
| Procellariiformes | *Diomedea exulans* | Snowy Albatross | UMZC 9/Dio/1/h/8 | Wet | Embryo | |
| Procellariiformes | *Macronectes halli* | Northern Giant Petrel | UMZC uncatalogued | Frozen | Adult | |
| Procellariiformes | *Macronectes giganteus* | Southern Giant Petrel | UMZC uncatalogued | Frozen | Hatchling | |
| Procellariiformes | *Fulmarus glacialis* | Northern Fulmar | UMZC 9/Pro/4/a/13 | Wet | Hatchling | |
| Suliformes | *Morus bassanus* | Northern Gannet | UMZC 10/Sul/1/a/6 | Wet | Juvenile | |
| Suliformes | *Morus bassanus* | Northern Gannet | UMZC 262.E | Osteo | Adult | |
| Pelecaniformes | *Egretta garzetta* | Little Egret | NMBE 174/72 | Osteo | Juvenile | |
| Pelecaniformes | *Nycticorax nycticorax* | Black-crowned Night Heron | NMBE 180/72 | Osteo | Adult | |
| Pelecaniformes | *Nycticorax nycticorax* | Black-crowned Night Heron | NMBE 176/72 | Osteo | Juvenile | |
| Pelecaniformes | *Ardea cinerea* | Grey Heron | UMZC 11/Ard/2/a/5 | Wet | Hatchling | |
| Opisthocomiformes | *Opisthocomus hoazin* | Hoatzin | UMZC 14/Opi/1/a/1 | Wet | Embryo | |
| Opisthocomiformes | *Opisthocomus hoazin* | Hoatzin | MNHF 1033857 | Osteo | Adult | |
| Accipitriformes | *Vultur gryphus* | Andean Condor | UMZC 2022.11.31 | Wet | Embryo | |
| Accipitriformes | *Vultur gryphus* | Andean Condor | UMZC 519.AA | Osteo | Adult | |
| Piciformes | *Pteroglossus viridis* | Green Aracari | UMZC 2022.11.3 | Wet | Embryo | |
| Piciformes | *Pteroglossus viridis* | Green Aracari | BMNH S/1993.5.4 | Osteo | Adult | |
| Psittaciformes | *Lorius lory* | Black-capped Lory | UMZC 2022.11.6 | Wet | Embryo | |
| Psittaciformes | *Ara ambiguus* | Great Green Macaw | UMZC uncatalogued | Frozen | Adult | |
| Psittaciformes | *Ara macao* | Scarlet Macaw | UMZC 2022.11.13 | Wet | Embryo | |

**Supplementary table 2:** List of unfigured comparative specimens examined in this study**.** Abbreviations: NHM, Natural History Museum, London; UMMZ, University of Michigan Museum of Zoology, Michigan; UMZC, University Museum of Zoology, Cambridge.

| ORDER | TAXON | COMMON NAME | COLLECTION NUMBER | TYPE | ONTOGENETIC STAGE | MORPHOSOURCE ID |
|---|---|---|---|---|---|---|
| Tinamiformes | *Tinamus major* | Great Tinamou | NHMUK S/1968.7.1 | Osteo | Adult | |
| Tinamiformes | *Nothura maculosa* | Spotted Nothura | NHMUK S/2008.1.5 | Osteo | Adult | |
| Tinamiformes | *Eudromia elegans* | Elegant Crested-Tinamou | NHMUK S/1930.3.24.7 | Osteo | Subadult | |
| Tinamiformes | *Rhynchotus rufescens* | Red-winged Tinamou | NHMUK S/2010.1.1 | Osteo | Adult | |
| Tinamiformes | *Nothoprocta perdicaria* | Chilean Tinamou | UMZC 824 | Osteo | Adult | |
| Tinamiformes | *Nothoprocta perdicaria* | Chilean Tinamou | UMZC uncatalogued | Frozen | Subadult | |
| Anseriformes | *Chauna torquata* | Southern Screamer | NHMUK S-2012.31.1 | Osteo | Adult | |
| Anseriformes | *Anas platyrhynchos* | Mallard | UMZC 225 | Osteo | Adult | |
| Anseriformes | *Cygnus olor* | Mute Swan | NHMUK 1854.6.25.2 | Osteo | Adult | ID:000157589 |
| Anseriformes | *Dendrocygna bicolor* | Fulvous Whistling-Duck | UMMZ 219885 | Osteo | Adult | ID:000109604 |
| Anseriformes | *Malacorhynchus membranaceus* | Pink-eared Duck | UMZC 12-Ana-33-a-1 | Skin | Adult | |
| Anseriformes | *Anseranas semipalmata* | Magpie Goose | NHMUK:1852.7.22.1 | Osteo | Adult | ID: 000083594 |
| Galliformes | *Ortalis ruficauda* | Rufous-vented Chachalaca | UMZC uncatalogued | Frozen | Juvenile | |
| Galliformes | *Ortalis ruficauda* | Rufous-vented Chachalaca | UMMZ 155489 | Osteo | Adult | ID: 000092950 |
| Galliformes | *Coturnix coturnix* | Common Quail | UMMZ 224005 | Osteo | Adult | ID: 000109575 |
| Galliformes | *Rollulus rouloul* | Crested Partridge | NHMUK 1871.7.20.87 | Osteo | Adult | ID: 000092768 |
| Psittaciformes | *Psittacus erithacus* | Gray Parrot | NHMUK S/1992.41.60 | Osteo | Adult | ID: 000092678 |
